# Genomic evidence for within-species thermal adaptation in collagen genes

**DOI:** 10.64898/2026.10.23.761307

**Authors:** Joana L. Santos, Peter D. Fields, Luca Cornetti, Dieter Ebert

**Author notes:** Correspondence: Joana L. Santos. Department of Aquatic Ecology, Swiss Federal Institute of Aquatic Science and Technology (Eawag), Ueberlandstrasse 133, 8600 Dübendorf, Switzerland (Joana L. Santos). Syngenta Crop Protection AG, Schaffhauserstrasse 101, 4332 Stein, Switzerland.

## Abstract

Collagen is the most abundant protein in animals. Its thermal stability depends on its specific amino acid composition within the repeated Gly-_Xaa_-Y_aa_ motifs forming the triple helix structure. Proline increases collagen rigidity, while serine enhances its elasticity—properties that contribute to collagen function and that vary with temperature. Although variation in collagen composition has been linked to temperature adaptation across evolutionary distinct taxa, its role within a species remains unclear. Here, we tested whether collagen genes across populations of the freshwater crustacean, *Daphnia magna*, show a signature of local adaptation to the ambient temperature. Analysing 39 collagen genes from 215 *D. magna* populations across three phylogeographic lineages spawning four continents, we found that animals from populations in warmer climates have higher proline and lower serine content in collagen—consistent with enhanced protein stability at higher temperatures. This variation was also associated with experimentally measured upper temperature tolerance. Parallel directional patterns of amino acid change appeared across genetically distinct lineages. Collagen genes also showed strong signatures of positive selection, comparable to that of immune-related genes, supporting a scenario of polygenic molecular role in adaptive responses. Our findings provide evidence of intraspecific molecular evolution in collagen amino acid composition driven by environmental temperature. Additionally, our study bridges macroevolutionary patterns with microevolutionary processes, posing the first intraspecific evidence that collagen genes can contribute to responses to environmental gradients at the molecular level, and thus, advancing our understanding of molecular adaptation of constrained structural proteins in an environmental context.

## Introduction

As the principal structural protein in metazoans, collagen accounts for about 30% of all animal proteins and is essential for maintaining the strength and structure of bones, skin, cartilage and blood vessels (Kielty and Grant 2002). The collagen gene family is highly diverse and different proteins fulfil various functions (Haq et al. 2019). Despite this diversity, the structure of collagen proteins is conserved among taxa and genes, characterized by a triple helix comprised of a repeated glycine followed by two other amino acids (Gly-X_aa_-Y_aa_) (Kielty and Grant 2002). While this structure is conserved, the amino acid composition can vary among animals, being shaped to some degree by the environment (Szpak 2011).

Within the collagen triple helix, the amino acid proline is one of the most abundant amino acids and a key predictor of protein stability (Shoulders and Raines 2009). The biophysical properties of collagen are well described: collagen stability declines with increasing temperatures, but a higher proline content can enhance thermal stability by increasing rigidity (Rigby 1968; Shoulders and Raines 2009; Fujii et al. 2022). Accordingly, it has been hypothesized that proline (and its hydroxylated form, hydroxyproline, produced by a post-translational modification) content in collagen may evolve in response to the environmental conditions in which an organism is found (Fujii et al. 2022; Tassara et al. 2023), such that higher proline content is expected in animals living in warmer environments (Akita et al. 2020). This proline–temperature hypothesis is supported by comparative studies across animal taxa. One hydrothermal annelid species, for example, has a significantly higher proline content than its cold-water relatives (Sicot et al. 2000). Similarly, fish collagen contains less proline and hydroxyproline than terrestrial mammals, and cold-water fish collagen has less proline than tropical fish collagen (Lin and Liu 2006; Szpak 2011; Akita et al. 2020). Accordingly, cold-water fish tend to have the lowest collagen thermal stability, followed by other ectotherms (Rigby 1968; Harvey et al. 2021). Another important amino acid in temperature regulation is serine, which shows a strong negative linear association with proline (Rigby 1967). In contrast to proline, serine increases protein elasticity (Liao et al. 2019) and is found more abundantly in the collagen of animals from colder environments (Lin and Liu 2006; Szpak 2011; Akita et al. 2020). Together, these findings suggest that environmental temperature can shape collagen amino acid composition through selection to balance proline and serine content.

Macroevolutionary patterns across taxa imply adaptation of collagen amino acid composition to environmental temperature. It remains unclear whether such patterns arise through similar processes within species and across populations. At the intraspecific level, evidence for thermal adaptation at the protein levels has so far come mainly from studies based on stress-response proteins, with heat shock proteins being the most studied example (Chen et al. 2018). These differences in population thermal tolerance associated with changes in heat shock proteins expression are typically mediated by changes in gene regulation or expression, rather than by changes in the amino acid sequence (Chen et al. 2018). In fact, heat shock proteins are highly conserved at the sequence level. In contrast, evidence for intraspecific thermal adaptation involving structural proteins is comparatively scarce. One notable exception are antifreeze proteins in cold-adapted fish species, where populations can differ in gene copy number, allelic composition, and expression levels along thermal gradients, resulting in local adaptation in cold resistance (Cheng 1998; Graham and Davies 2024). Whether similar forms of intraspecific molecular adaptation occur in other structural proteins under strong functional constraint, remains largely unexplored. Moreover, in structural proteins, with a key role in maintaining cell integrity, amino acids substitutions can have profound effect on protein stability and function, and thus, it is expected that at the sequence level these proteins evolve more slowly (Shoulders and Raines 2009). Altogether, this raises the question if changes in proline and serine content in collagen can also evolve within populations of a species in response to environmental temperatures, as is observed across divergent taxa.

Here, we explore whether collagen genes show a signature of local adaptation across genetically distinct populations of a widespread pond-dwelling planktonic species, *Daphnia magna*, which inhabits a wide range of thermal environments (Ebert 2022), by identifying changes in collagen amino acid composition at the sequence level. We use whole-genome sequence data from 215 populations across the Holarctic and representing the three major phylogeographic lineages (Fields et al. 2018; Cornetti et al. 2019), to test the prediction that environmental temperature associates positively with proline and negatively with serine content of collagens, as predicted by the proline-temperature hypothesis. We further examine whether these patterns are associated with experimentally measured thermal tolerance of this crustacean (Seefeldt and Ebert 2019), and test for signatures of selection in collagen genes by conducting a population genetic analysis. By integrating environmental, phenotypic, and genomic data, our approach enables to assess if molecular evolution in collagen genes leads to local adaptation and follows parallel trajectories on different continents. If this is confirmed, it would indicate that even highly constrained structural proteins can contribute to intraspecific local thermal adaptation.

## Materials and Methods

### Study design

To evaluate the amino acid composition—specially of proline and serine amino acids—of collagen genes, whole genome sequences of 215 populations from the freshwater crustacean, *D. magna*, spanning four continents, three distinct genetic lineages and distinct thermal gradients were used (Fig. 1 and Table S1). The samples belong to the *Daphnia magna Diversity Panel*, a standing collection of hundreds of genotypes each from distinct populations and fully sequenced, available at the Ebert-research group at the University of Basel. From the *D. magna* genom,e collagen genes were identified and, the amino acid content within the Gly-X_aa_-Y_aa_ motifs was estimated for each genotype. Proline and serine content were related to the local mean annual temperature from genotype’s origin and associated with temperature tolerance. Note that we could not account for hydroxyproline content with our approach, since this it is not a genetically encoded amino acid, but a post-translational modification of proline (Fujii et al. 2022). These associations were further evaluated for correcting phylogenetic structure. Population genetic statistics were conducted to identify signatures of selection.

**Figure 1.**
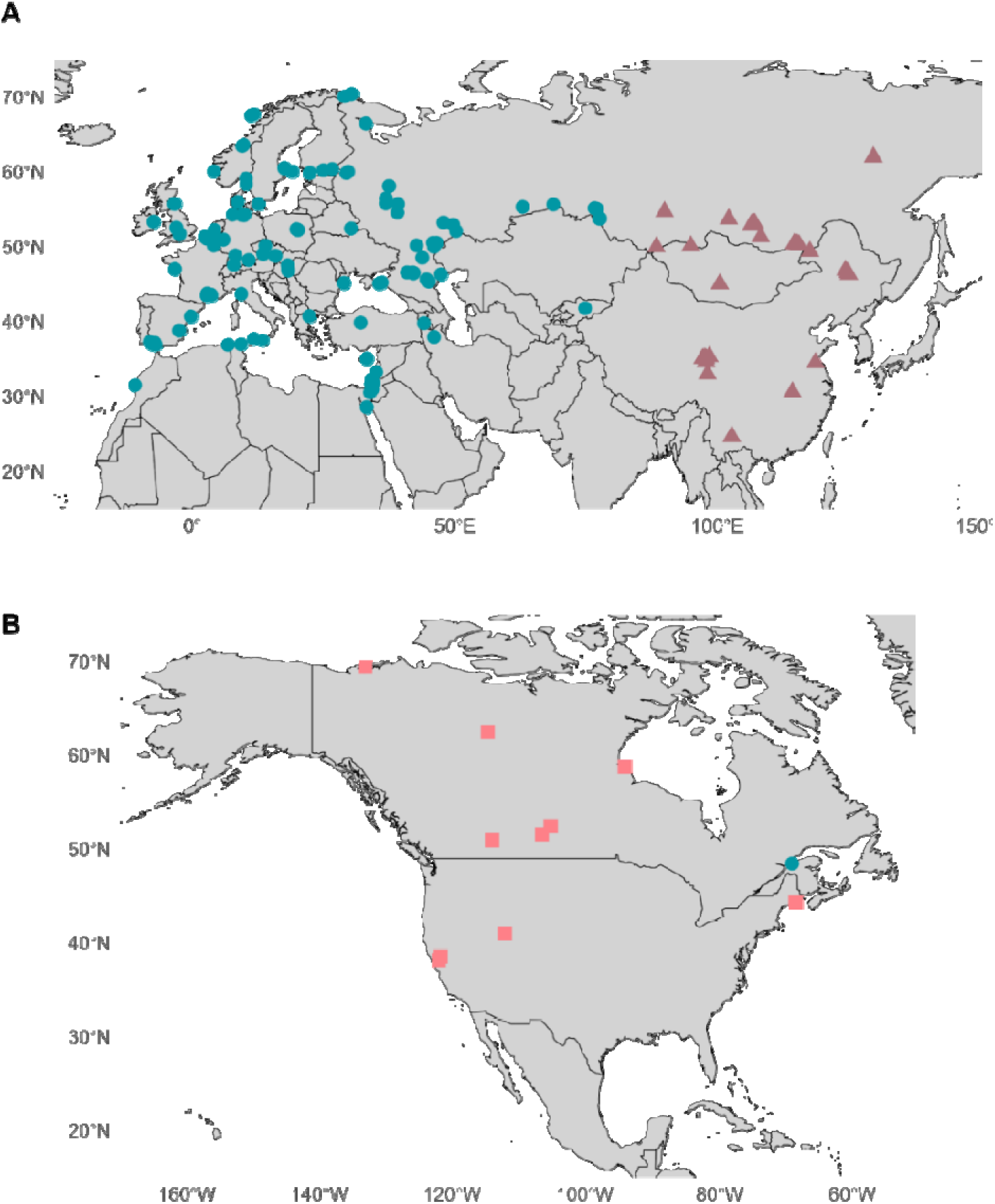
Samples distribution. Geographic distribution of the 215 *D. magna* genotypes used in this study across Eurasia (A) and North America (B). Each dot represents one genotype from a single population. Dots are colored according to genetic lineage, with the Western Eurasian lineage in blue, East Asian lineage in purple and the North American lineage in pink.

### Whole genome sample data

Whole genome paired-end Illumina sequencing was conducted for 215 *D. magna* genotypes from three genetic lineages: Western Eurasia (n=161), East Asia (n=35) and North America (n=19) (Fig. 1 and Table S1; NCBI Database; BioProject ID PRJNA1059512). The low sample size of *D. magna* from North America, is explained by the scarcity of the species on this continent. DNA library preparation, sequencing and quality assessment are described in Fields *et al*. (2022). Sequence reads were mapped to the v.3.0 *D. magna* reference genome (NCBI Database; BioProject ID PRJNA624896), and single nucleotide polymorphisms (SNPs) were identified using GATK v.3.8 (McKenna et al. 2010) ‘HaplotypeCaller’ according to GATK Best Practices recommendations. Filtering was applied with BCFtools (Danecek et al. 2021) and VCFtools v.0.1.16 (Danecek et al. 2011) to include only biallelic positions and ensure a minimum depth of coverage of 10 and quality thresholds—minimum genotype quality of 20 and minor allele count of 4. The final variant calling file (VCF) contained 6,603,086 SNPs.

### Amino acid content in collagen genes

Collagen genes were identified in the *D. magna* reference genome using *hmmsearch* (Eddy and HMMER development team 2020), based on the repetitive Gly-X_aa_-Y_aa_ motifs. Of the 94 potential collagen genes identified, 50 single isoform genes contained sequences with more than 20 repetitive Gly-X_aa_-Y_aa_ motifs and were shared across all individuals (Table S2).

For each collagen gene, the coding sequences of each genotype and respective haplotype were extracted and converted into protein sequences based on the gene annotations of the *D. magna* reference genome (NCBI Database; BioProject ID PRJNA624896) using BCFtools *consensus* v1.9 (Danecek et al. 2021), BEDtools *getfasta* v2.29.0 (Quinlan 2014) with the *-s* flag for genes in the reverse direction, and Transeq v6.6.0.0 (Rice et al. 2000). To better estimate haplotype sequences, the VCF file was previously phased using Beagle v5.3 (Browning et al. 2021).

Amino acid content was calculated within repeated regions of at least 20 Gly-X_aa_-Y_aa_ motifs, which sometimes identified multiple regions within a gene (*see* Table S2). Protein-coding regions were identified with a custom script (*available in* https://github.com/JLS52/col_dmdp082023.git), and extracted using seqkit subseq v2.3.1 (Shen et al. 2016). For each genotype, the amino acid content was calculated per 1000 residues for each region of interest and inferred haplotype, then averaged across regions (with equal contribution) and haplotypes using matrixStats package v0.61.0 (Bengtsson et al. 2021) in R v4.0.2 (R Core Team 2021). The overall amino acid content estimated for *D. magna* resulted from an averaged value across all collagen genes and genotypes used in this study.

Eleven collagen genes with high-impact mutations observed at the start and/or stop codon level were excluded from analysis as they could potentially affect translation and results interpretation (Table S2). Therefore, the main analysis included a total of 39 genes, which are present in the 215 genotypes included in the study and contain a complete protein fasta sequence to allow comparison across genotypes. Although, the complete dataset (50 collagen genes) analysis is included in supplementary material (Table S5). All genes from the complete dataset, except one of the eleven excluded genes had previously been shown to be expressed in *D. magna* (Orsini et al. 2016).

### Environmental temperature data

Annual mean temperature at the genotype’s site of origin was retrieved from the Bioclim dataset (available at https://www.worldclim.org/data/bioclim.html) of WorldClim v2 (Bioclim 1) database(Fick and Hijmans 2007) for 1 km^2^ resolution using the R package foreach V1.5.2 (Daniel et al. 2022) (Table S1). For sites located near the coast or on small islands, climate data were obtained from the nearest land-based GPS coordinates no further than 5 km from the original site.

### Statistical data analysis

Separate multiple linear regression models (MLRM) were used to assess the overall proline and serine contents as a function of annual mean temperature (Bioclim 1), incorporating phylogenetic lineage as a factor with the *lm* function. Separate models for each lineage were performed due to significant interactions. The mean values of amino acid content were reported together with the standard deviation. Model validation, with regard to data linearity, independence, normality, homoscedastic and influencing points, and data visualization was conducted using the R packages ggplot2 v3.3.5 (Wickham et al. 2020), data.table v1.14.2 (Dowle et al. 2019) and dplyr v1.0.7 (Wickham et al. 2021).

Phylogenetic generalized least squares (PGLS) analysis accounted for the phylogenetic signal in the data. A phylogenetic tree was generated from 100,000 random SNPs filtered for quality using VCFtools v.0.1.16 (Danecek et al. 2011) according to the following criteria: Minor allele frequency (MAF) > 0.05, at least 90% of the individuals with a genotype present, minimum genotype quality of 30, minimum depth of 10 and maximum depth of two alleles. Later, it was converted to phylip format using the vcf2phylip.py script (https://github.com/edgardomortiz/vcf2phylip). Maximum likelihood trees were built using RAxML-NG v.1.1.0 (Kozlov et al. 2019) under GTGTR substitutional model for unphased, diploid genotypes with the maximum likelihood-based best fit tree estimated allowing 25 replicate starting trees, and topological convergence assessed by Robinson-Foulds (RF) metric (Robinson and Foulds 1981). PGLS analysis input the phylogenetic tree and variables of interest (i.e., bioclimatic data, phylogenetic lineage and amino acid content) using caper v1.0.1(Orme et al. 2018) R package. The evolutions of both proline and serine content were mapped as continuous traits using phytools *contMap* v1.2.0 R package (Revell 2022).

### Temperature tolerance associations

Proline and serine content were associated with published temperature tolerance data for 72 Western Eurasian summer-active genotypes (Table S1) (Seefeldt and Ebert 2019), using similar MLRM as described above. Temperature tolerance was measured in a ramp experiment by increasing temperature in clonal asexually reproducing populations by 1 °C every two weeks until they died(Seefeldt and Ebert 2019).

### Detecting positive selection in collagen genes

To estimate the impact of selection on collagen genes, we calculated the ratio of nucleotide diversity at non-synonymous and synonymous sites (π_N_/π_S_) for the 39 collagen genes, and two other gene sets, the immune-related gene set originally described for *Drosophila* (see: https://www.epfl.ch/labs/lemaitrelab/lemaitre-lab/resources/list-of-drosophila-genes-potentially-involved-in-the-immune-response/), and the Arthropods BUSCO gene set (Benchmarking Universal Single-Copy Ortholog) (*see* Table S8). Immune genes are expected to show elevated π_N_/π_S_ due to rapid evolution (Shultz and Sackton 2019), while BUSCO genes are highly conserved with low π_N_/π_S_ under purifying selection (Waterhouse et al. 2010). Only Western Eurasian genotypes were analyzed to minimize bias due to population structure.

DNA alignments generated from inferred haplotypes from the same VCF file used to estimate amino acid content were analyzed with the selectionStats.py script (https://github.com/tatumdmortimer/popgen-stats). The overall nucleotide diversity (π), and the nucleotide diversity at the nonsynonymous (π_N_) and synonymous (π_S_) sites were calculated per gene of the three gene sets (Collagen n=39, Immune=298, BUSCO=797; Table S2 and S8), and gene set comparisons were done using Wilcoxon rank tests. Analyses were repeated excluding rock pool populations (130 genotypes remained for analysis) to account for the potential πN/πS overestimation due to the accumulation of mutations from repeated genetic bottlenecks in these populations.

## Results

### Amino acid content in collagen genes

A total of 50 collagen genes were identified in 215 *D. magna* genotypes originating from four continents: Europe, Africa, Asia and North America (Fig. 1 and Table S1). The main analysis included only 39 genes that were complete for all genotypes (i.e., without high-impact mutations), which comprised a hundred regions containing more than 20 continuous and repetitive Gly-X_aa_-Y_aa_ motifs (*see* Methods section; Table S2). As expected, glycine accounted for about one-third of the amino acid content within the Gly-X_aa_-Y_aa_ motifs of the collagen genes (overall mean: 341.56 ± 0.14 (STD) per 1000 residues) (Table S3 and S4). Proline was the second most common amino acid (overall mean: 211.26 ± 0.73 per 1000 residues); its content varied among genotypes in 38 genes, while one gene contained no proline (snap_masked-000011F-processed-gene-20.3-mRNA-1 gene). Serine content was 28.19 (± 0.36) per 1000 residues.

### Amino acid variation shows an association with environmental temperature

The annual mean temperature of origin for the *D. magna* genotypes ranged from negative 10.18 °C to positive 20.01 °C (mean 7.62 ± 6.63 °C) (Table S1). As predicted, there was an overall positive association between proline content and annual mean temperature, explaining 18 % of the variance (MLRM: DF = 1, F-statistics = 100.61, *p*-value <2.2×10-16, slope = 0.08; Fig. 2A and Table 1). Also consistent with our prediction, serine content showed a negative association with annual mean temperature (MLRM: Adjusted-R^2^ = 0.02, DF = 1, F-statistics = 8.08, *p*-value = 4.9×10^-3^, slope = -0.03; Fig. 2B and Table 1). The effects of temperature on proline and serine content were dependent on phylogenetic lineage (MLMR: Interaction annual mean temperature and phylogenetic lineage – DF = 2, F = 10.46, *p* = 4.7×10^-5^ for proline and DF=2, F = 4.64, *p* = 0.01 for serine). Specifically, proline and serine associated with annual mean temperature for the Western Eurasian and the East Asian lineages, but not for the North American lineage (Table 1). The North American lineage also had the lowest proline and highest serine average values (Fig. 2). These findings were consistent for all 50 collagen genes identified in the *D. magna* genome (*see* Methods section and Table S5). After correcting for the phylogenetic signal (PGLS analysis), the associations between annual mean temperatures and proline and serine content retained the same direction (positive and negative, respectively) but remained significant only for proline (Table S6, Fig. S1 and S2). In addition, proline and serine content were strongly and negatively associated (LRM: Adjusted-R^2^ = 0.62, DF = 119, F = 352.7, *p* < 2.2×10^-16^, slope = -0.39).

**Figure 2.**
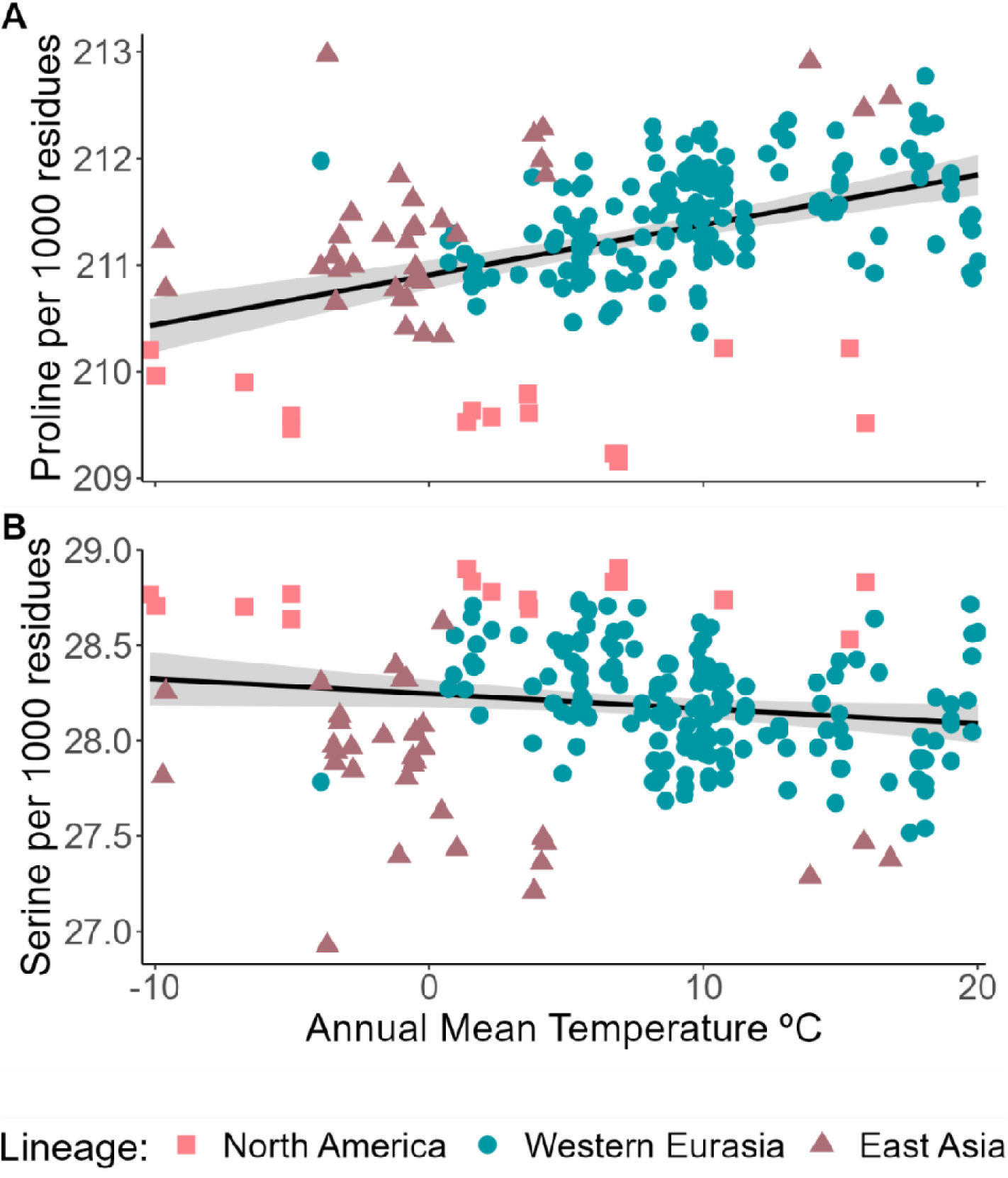
Amino acid content of collagen genes related to annual mean temperature at the *Daphnia magna* genotype’s site of origin. Variation in the number of proline (A) and serine (B) amino acids per 1000 residues in relation to annual mean temperature for 39 collagen genes from the 125 D. magna genotypes used in this study. Each dot represents one sample, colored according to genetic lineage. The black line represents the linear regression model between the two variables including the 95 % confidence interval of the mean, shaded in grey.

**Table 1.** Description of proline and serine amino acid content across 39 *D. magna* collagen genes, and its association with annual mean temperature. Summary statistics show the sample size (= number of genotypes) (N), mean number of an amino acid per 1000 bp (Mean), standard deviation of the mean (STD), maximum (Max) and minimum (Min) values. A multiple regression model was performed on the entire dataset (All) using annual mean temperature at the site of genotype origin (Temp.), phylogenetic lineage and their interaction (Int.) as independent factors. Independently, a linear regression model was also conducted for each phylogeographic lineage: Western Eurasia (WA), East Asia (EA) and North America (NA). The degrees of freedom (Df), the F-value, p-value and percentage of explained variance (Exp. Var; in %) are shown for each regression model. Significant p-values of a threshold at 0.05 are in bold.

|  |  |  |  |  |  | Regression model |  |  |  |  |  |
| --- | --- | --- | --- | --- | --- | --- | --- | --- | --- | --- | --- |
|  | Data | N | Mean (STD) | Max | Min | Factors | Df | F-value | p-value | slope | Exp. Var. |
| Proline | All | 215 | 211.26±0.73 | 212.97 | 209.16 | Temp. | 1 | 100.61 | <2.2x10<sup>-16</sup> | 0.08 | 17.93 |
|  |  |  |  |  |  | Lineage | 2 | 115.34 | <2.2x10<sup>-16</sup> | - | 41.10 |
|  |  |  |  |  |  | Int. | 2 | 10.46 | 4.7x10<sup>-5</sup> | - | 3.73 |
|  | WA | 161 | 211.45±0.49 | 212.77 | 210.37 | Temp. | 1 | 40.37 | 2.1x10<sup>-9</sup> | 0.04 | 20.25 |
|  | EA | 35 | 211.33±0.71 | 212.97 | 210.34 | Temp. | 1 | 20.52 | 7.3x10<sup>-5</sup> | 0.08 | 20.52 |
|  | NA | 19 | 209.60±0.36 | 210.22 | 209.16 | Temp. | 1 | 1.03 | 0.32 | -0.01 | 1.03 |
| Serine | All | 215 | 28.19±0.36 | 28.90 | 26.93 | Temp. | 1 | 8.08 | 4.9x10<sup>-3</sup> | -0.03 | 2.05 |
|  |  |  |  |  |  | Lineage | 2 | 84.04 | <2.2x10<sup>-16</sup> | - | 42.61 |
|  |  |  |  |  |  | Int. | 2 | 4.64 | 0.01 | - | 2.35 |
|  | WA | 161 | 28.19±0.27 | 28.74 | 27.52 | Temp. | 1 | 20.58 | 1.1x10<sup>-5</sup> | -0.02 | 11.46 |
|  | EA | 35 | 27.85±0.39 | 28.62 | 26.93 | Temp. | 1 | 10.01 | 3.3x10<sup>-3</sup> | -0.03 | 23.37 |
|  | NA | 19 | 28.78±0.10 | 28.90 | 28.53 | Temp. | 1 | 0.31 | 0.58 | 1.7x10<sup>-3</sup> | 1.82 |

631 \*p<0.05

### Amino acid content predicts temperature tolerance

For 72 *D. magna* genotypes from the summer-active populations of the Western Eurasian lineage, the maximal tolerated survival temperature was previously experimentally assessed and ranges from 27.5 to 32.5 °C (mean of 30.86 ± 1.33) (Seefeldt and Ebert 2019). These estimates of temperature tolerance associated positively with proline and negatively with serine content, explaining 8.4 % and 6.3 % of the variation, respectively (LRM: DF = 1, F = 7.51, *p* = 7.8×10^-3^ for proline and DF = 1, F = 5.7, *p* = 0.02 for serine).

### Genetic diversity and signatures of positive selection in collagen genes

Our findings suggest that collagen genes are diverse and under positive local selection. To scale these features, we compared them with a group of highly conserved genes (BUSCO genes set— Benchmarking Universal Single-Copy Ortholog genes) and with a group of genes known to evolve rapidly (immune gene sets). Nucleotide diversity was generally low in BUSCO genes (2.8×10^-3^ ± 0.002) and significantly higher in immune genes (7.6×10^-3^ ± 0.01) and collagen genes (7.9×10^-3^ ± 0.004). The πN/πS ratios in collagen genes (0.27 ± 0.18) and immune genes (0.30 ± 0.28) were rather high and similar (Wilcoxon test: W = 5537, p-value = 0.93) (Fig. 3), and were significantly higher than the mean πN/πS for the BUSCO genes (0.14 ± 0.22) (Wilcoxon test: W = 174528, p-value < 2.2×10^-16^, and W = 23256, p-value = 9.1×10^-9^, respectively) (Fig. 3). These results remained unchanged when the genotypes from rock pool populations were excluded (*see* Table S7), which are known for their high genetic load.

**Figure 3.**
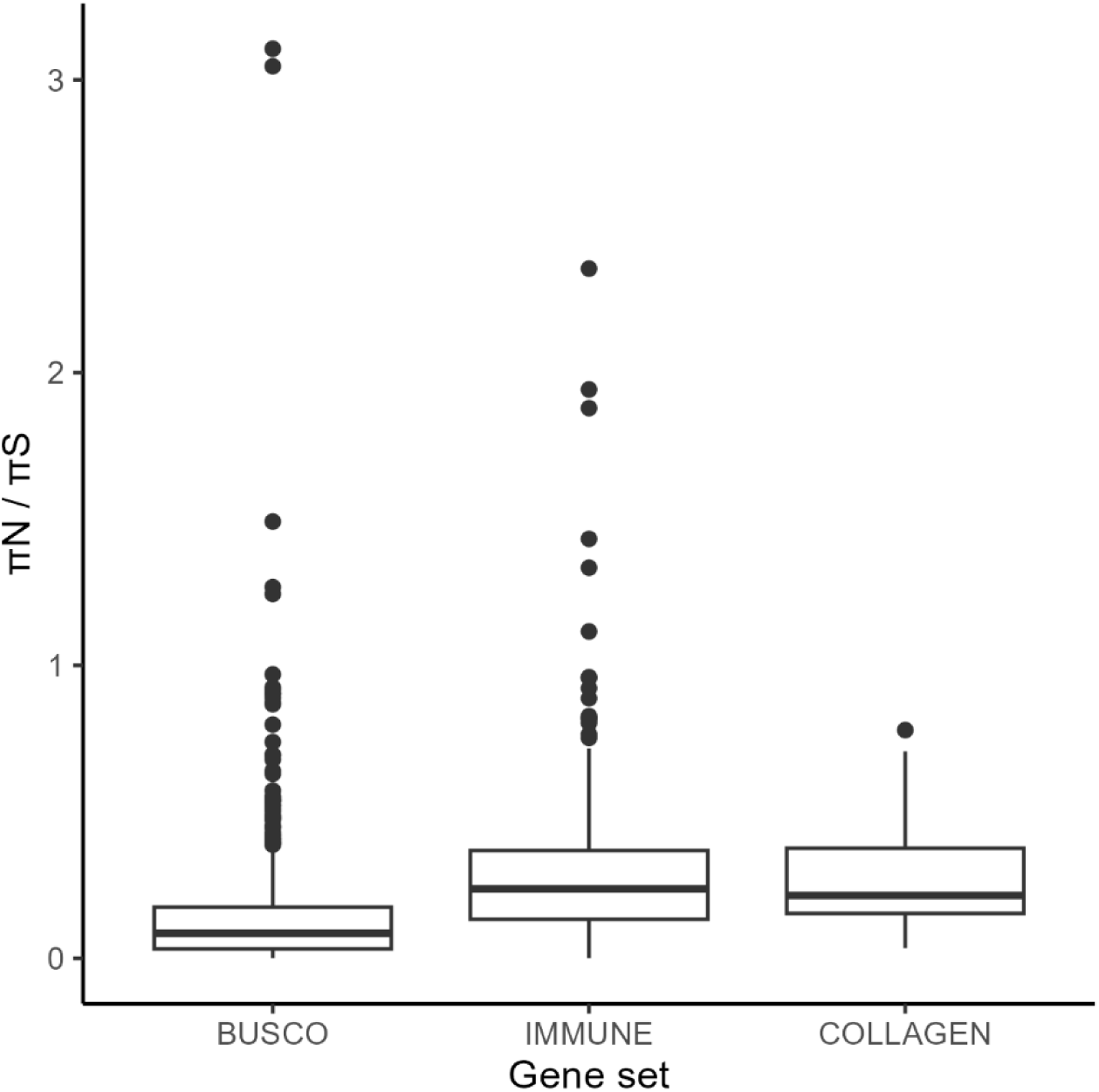
Nucleotide diversity among gene sets. Comparison of estimated nucleotide diversity ratios (πN/πS) for BUSCO (n=797), immune (n=298) and collagen (n=39) genes for the Western Eurasian phylogenetic lineage. Boxplots show the median, first and third quantiles. Whiskers extend to 1.5 times the upper and lower limits of the interquartile range. Dots are outside the 1.5 times from the interquartile range. BUSCO genes have a significantly lower median than the other two gene sets, which do not differ significantly from each other.

## Discussion

The thermal stability of collagen, an important structural protein of all animals, depends on the amino acid composition within its triple helix structure (Shoulders and Raines 2009). Our genomic analysis evaluated how environmental temperature shapes evolution in the amino acid composition of collagen genes across distinct phylogenetic lineages of a single freshwater species. As predicted warmer environments favoured an increased proportion of proline, while the serine content increased in cooler conditions. This finding is further supported by similar associations of proline and serine content with published records of experimentally assessed temperature tolerance (2019), and by strong signatures of positive selection found in collagen genes. Our results further showed that variation in collagen gene composition, especially of proline and serine content, can evolve independently in distinct phylogeographic lineages of *D. magna*, providing evidence of directional and repeatable molecular evolution at the intraspecific level. The changes in amino acid composition were evident across 39 collagen genes emphasizing the polygenic nature of temperature adaptation. Altogether, these findings reveal that even highly conserved structural proteins such as collagen, typically thought to be evolutionary constrained, evolve as an intraspecific adaptive response to environmental temperature, which had previously only been documented in macroecological comparisons across taxa (Szpak 2011).

### Temperature shapes collagen amino acid composition

Our study supports the proline hypothesis for temperature adaptation within the widespread planktonic species *D. magna*. Beyond temperature associations, several lines of evidence support this hypothesis. In particular, studies across taxa have shown similar patterns of thermal adaptation among organisms from diverse temperature environments, including comparisons between cold- and warm-blooded organisms (Lin and Liu 2006; Szpak 2011; Akita et al. 2020; Harvey et al. 2021). The averaged values reported in our study for proline are in the range of those found in bone and skin collagens of mammals, while serine it is slightly lower than in fish species, as reported in the study of Szpak (2011), a compilation of existing literature. Our study demonstrates that the convergence of the identified pattern at inter- and intraspecific levels shows how selection by temperature shapes collagen amino acid composition, even at a microevolutionary level. Additionally, the outcome of the trade-off between proline and serine content aligns with the known biophysical properties of these amino acids, which are crucial to the structural function of collagen and its temperature dependence. Proline, by imposing conformational constraints and increasing collagen’s melting point, leads to an increase in collagen rigidity, stabilizing protein structure and helping it withstand elevated temperatures that may causes protein denaturation (Liao et al. 2019; Fujii et al. 2022). In contrast, serine enhanced elasticity and protects against collagen rigidity in colder environments (Fields 2001; Fujii et al. 2022). In line with this, our study also revealed a clear negative association between proline and serine content. Finally, we show that the empirically assessed maximum tolerated temperature for each *D. magna* genotype associates positively with proline (Seefeldt and Ebert 2019), linking the molecular pattern to the predicted phenotype.

### Parallel and polygenic molecular evolution

Our Holarctic sampling scheme enabled us to test for repeated and parallel evolution across three continental temperature gradients. Both the Western Eurasian and East Asian phylogenetic lineages, which have been isolated approximately 1.2 million years ago (Cornetti et al. 2019), showed positive associations between temperature and proline content (and inversely for serine). However, the more recent post-glacial colonization of northern habitats (Fields et al. 2018) likely drove the current temperature adaptation patterns in *D. magna*, as is consistent with the evolutionary histories of many other species that expanded and diversified from southern refugia when the ice retracted about 10,000 years ago in the Northern Hemisphere (Hewitt 1999). Indeed, the Western Eurasian lineage seems to have used a single refugia during the last ice-age in the Middle East (Fields et al. 2015). The relatively subtle genetic footprints of thermal adaptation observed here for *D. magna* may indicate that the adaptive process has not yet reached its optimum. This parallelism supports the idea that thermal adaptation in collagen genes follows a predictable trajectory, reinforcing how adaptive evolution can be repetitive under similar environmental pressures.

Our findings hold up within a phylogenetic context. We observed not only the association between amino acid composition and environmental temperature in two lineages, but also that this relationship persisted even after correcting for the phylogenetic structure of the dataset. Although, this was only marginally significant for serine, possibly reflecting the conservative nature of the implemented phylogenetic correction which may have overcorrected our dataset. In *D. magna*, geography shapes both climate and genetic relatedness within each lineage (isolation by distance (Fields et al. 2015)), which reduces the power to unveil signatures of thermal adaptation, especially in traits with smaller effect such as serine content. Nonetheless, the consistent direction of the significant associations observed after phylogenetic correction indicate that, despite the divergence of both lineages, the pattern of collagen amino acid variation is similar. This pattern is compatible with parallel evolution along similar temperature gradients, rather than shared ancestry, although not fully excluding the potential contribution of the phylogenetic context. Together, these results highlight that adaptive and historical processes are not mutually exclusive and show the need of interpreting phylogenetically corrected models cautiously in spatially structured systems.

The link between several collagen genes and temperature emphasizes the polygenic nature of thermal adaptation in *D. magna*. Many studies have reported that genomic responses to temperature variation is a polygenic trait in diverse organisms, including associations with genes from various families (De La Torre et al. 2021). Amino acid variation and positive selection in collagen genes in response to environmental conditions have also been documented (Yudin et al. 2017), emphasizing the dynamic role of collagen in thermal adaptation, despite its apparent conserved status among animal taxa (Haq et al. 2019). It is likely that alongside collagen, other gene families may also contribute to temperature adaptation in *D. magna*, as the case of heat shock protein expression (Scheffer et al. 2022).

The evolutionary change in proline and serine content may be facilitated by the proximity of their codons in the genetic code. A single point mutation at the first codon position can switch between proline (CCx, where x is U, C, A or G) or serine (UCx). This may explain the strong negative association between proline and serine amino acids ((Rigby 1967); and this study). In addition, the high mutation rate estimated for the nuclear genome of *D. magna*—6.66–11.97 x 10^-9^/bp/generation (95% CI)(Ho et al. 2020)—combined with the large number of asexual generations per year (around 10 in most permanent water bodies and 2-5 in temporary ponds (Ebert 2022)), and the often large population sizes (e.g. larger than 10^8^ reported in Ameline *et al*.(2022)), create ample opportunities for beneficial mutations to arise across the many sites of the varied collagen genes.

Because the observed variation in proline and serine content among genotypes is modest, it may seem opposed to the idea of a strong selective process. However, given the polygenic context of collagen adaptation, we expect very small selection coefficients for individual sites, potentially so low that genetic drift could render selection inefficient (John and Stephan 2020). Indeed, while many *D. magna* populations are large (Ameline et al. 2022), others may often experience bottlenecks during the colonization of empty habitat patches, which can impede adaptive evolution (Angst et al. 2022). Thus, collagen adaptation to local temperatures is likely a slow and highly variable process across locations and demographic scenarios. The high-expected number of potentially beneficial mutations ameliorates the weak effect of selection, although it is likely that the current picture may not reflect the fitness peak of local populations in each habitat. Temperature adaptation of collagen could be lagging, and this time lag could become even more pronounced with the changing climate.

### Exceptions and limitations of general patterns

Interestingly, the observed pattern of amino acid composition across temperatures was not detected in the North American lineage, which consistently showed lower proline and higher serine contents than the Eurasian lineages, and possibly indicating adaptation to low temperatures (Fig. 2). Because of our small sample size for this lineage, we cannot draw clear conclusions, but several non-mutually exclusive processes could explain these results. Lineage-specific responses may be mediated by phylogenetic constraints, demographic history, or differences in available genetic variation (Rosenblum et al. 2014). On one hand, historical processes may contribute to this pattern. We hypothesize that the North American lineage may have survived in a cold, but ice-free refugia (e.g., Alaska (Löfverström and Liakka 2016)) during ice ages and spread more recently. On the other hand, we cannot exclude that population dynamics and selection processes can limit the detection of the amino acid composition pattern across temperatures. For example, strong population genetic structure coupled with isolation by distance may reduce effective recombination and limit the emergence of clear patterns. Conversely, ongoing gene flow among populations occupying different thermal regimes, although not expected here, could weaken local adaptation. While the North American findings challenge the generalized patterns of parallel evolution and predictability, it also stresses the importance of replicating environmental associations across multiple lineages and ensuring proper sample sizes to identify universal trends. Other well-studied systems have also shown that parallel evolution seems prevalent but not universal (Stuart et al. 2022).

### Signatures of positive selection support adaptive evolution

The elevated πN/πS ratio in collagen genes as opposed to housekeeping genes (here represented by the BUSCO genes set), for which a slow evolution rate is typically assumed (Waterhouse et al. 2010), suggests directional selection within populations and divergence across them (Kawecki and Ebert 2004). In *D. magna,* the rate of collagen evolution seems to be similar to that of immune genes, which are notorious for their fast evolution (Shultz and Sackton 2019). These findings further support that amino acid changes in collagen genes are driven by selection and not due to relaxed constraint or drift. Populations with variable habitat conditions and moderate gene flow, such as *D. magna* populations, often show a geographic pattern of isolation by distance, which facilitates adaptive evolution by spreading new mutations across near-by habitats without homogenizing gene pools (Kawecki and Ebert 2004; Fields et al. 2015). The substantial standing genetic variation coupled with pronounced habitat structure (Chaturvedi et al. 2021; Fields et al. 2022) explains this species’ ability to adapt to environmental conditions. Previous studies have documented such adaptability capacity of *Daphnia* for traits such as salinity tolerance (Santos et al. 2024), heavy metals resistance(Agra et al. 2010), cyanobacteria (Schwarzenberger et al. 2020) and the onset of diapause (Santos and Ebert 2022).

### Broader implications

Differences in collagen amino acid composition across taxa have long been interpreted as adaptations to varying thermal environments. However, the microevolutionary processes behind this macroevolutionary pattern, which we have undertaken to focus here, have remained largely underexplored. Our results indicate that microevolutionary processes can generate repeatable molecular trends, even in highly constrained structural proteins at the intraspecific level. Parallel patterns of molecular evolution in collagen composition across multiple lineages suggest directional tendencies when functional constraints and mutational opportunities are similar, while exceptions (e.g., North America lineage) highlight the influence of demography, population dynamics and lineage-specific constraints. Altogether, it also highlights how microevolutionary changes can contribute to broader evolutionary trends.

Our findings further provide an example of how genomic variation in coding regions can adjust the adaptive responses of complex protein families to environmental conditions, even in a polygenic trait. Given the great importance of collagen in animals, our genome-based approach for estimating collagen composition is easily adaptable to other species with available genomic data, offering a valuable tool for studying adaptation across environments at both species and population levels. Specifically, our results suggest that collagen evolution may serve as molecular marker of thermal adaptation. This is not only relevant to evolutionary ecology, as it can inspire future research in species inhabiting variable thermal habitats, but also to biotechnology and industry (e.g., medicine, cosmetic and food industries), looking for thermally resilient collagen properties (Zheng et al. 2023).

Finally, by demonstrating the repeatable, directional adaptability in structural proteins, as collagen, to temperature variation, our study contributes to a deeper understanding of the molecular basis of thermal adaptation, specifically in a keystone species from freshwater ecosystems, where the impacts of environmental changes have been severe. Generally, identifying the adaptive mechanisms enabling species to adapt to varying environments is crucial for predicting their responses to climate change. Species with such flexibility have higher chances of successfully colonizing new environments, highlighting the importance of such adaptations for ecosystem resilience and biodiversity conservation.

## Data and code availability

Raw genomic data and the reference genome of *D. magna* are deposited in NCBI database (BioProject ID PRJNA1059512 and PRJNA624896, respectively). The filtered VCF file used in the analysis will be available upon acceptance. The remaining data produced in this study and required for replication is available in the Supplementary Information file. All the code used for data analysis and to produce figures and tables is available in Github (https://github.com/JLS52/col_dmdp082023).

## Author Contributions

JLS and DE designed the project; DE curated the *Daphnia magna* diversity panel; JLS and PDF performed the data analysis; LC, DE and PDF provided guidance with data analysis; PDF produced the genome sequences; JLS wrote the manuscript with comments from all authors; DE supervised this project.

## Acknowledgments

We thank Pascal Angst and Jonathon H. Stillman for feedback during this project, and Louis du Pasquier for his expertise regarding genes with immune functions. Thanks also to members of the Ebert group for valuable insights on the manuscript draft. Suzanne Zweizig improved the language of the manuscript.

