## Supplementary Material for "Genomic evidence for within-species thermal adaptation in collagen genes"

Title:

### **Table S1.** Geographic coordinates and annual mean temperature (°C) of the locations from which the D. magna genotypes used in this study originated. The first two letters of the genotype label indicate the country. Samples with an asterisk (*) are originated from rockpools populations. In samples with an (^+^) previous estimates of maximal tolerated temperature are available in Seefeldt and Ebert^1^.

| **Genotype** | **Lineage** | **Latitude** | **Longitude** | **Annual Mean Temperature** |
| --- | --- | --- | --- | --- |
| AM-AR1-1 | Western Eurasia | 39.75169 | 44.8064 | 12.33 |
| BE-HO-1^+^ | Western Eurasia | 50.14506 | 5.077083 | 9.70 |
| BE-OHZ-T10^+^ | Western Eurasia | 50.83949 | 4.655044 | 10.24 |
| BE-OM-1^+^ | Western Eurasia | 50.86319 | 4.723872 | 10.39 |
| BE-T1-2 | Western Eurasia | 50.82305 | 4.593936 | 10.02 |
| BE-WE-59^+^ | Western Eurasia | 51.06778 | 3.773611 | 10.74 |
| BE-WH1-2^+^ | Western Eurasia | 51.33632 | 3.348728 | 10.27 |
| BE-WH2-11 | Western Eurasia | 51.33478 | 3.348134 | 10.27 |
| BE-WTE-1-1^+^ | Western Eurasia | 50.82836 | 4.639056 | 10.22 |
| BY-G-9^+^ | Western Eurasia | 52.42146 | 31.01378 | 6.95 |
| CA-RI-1*^+^ | Western Eurasia | 48.44487 | -68.5847 | 3.26 |
| CH-H-1^+^ | Western Eurasia | 47.55777 | 8.862608 | 9.42 |
| CY-PA1-1 | Western Eurasia | 35.03148 | 33.97376 | 19.03 |
| CY-PA2-1 | Western Eurasia | 35.03276 | 33.95505 | 19.04 |
| CY-PA3-1 | Western Eurasia | 35.03406 | 33.95493 | 19.03 |
| CY-PA3-2 | Western Eurasia | 35.03406 | 33.95493 | 19.03 |
| CZ_DO_1 | Western Eurasia | 49.00592 | 14.44142 | 8.62 |
| CZ_SUR_1 | Western Eurasia | 49.01075 | 14.43742 | 8.63 |
| CZ-KO-1^+^ | Western Eurasia | 50.12536 | 14.86867 | 9.32 |
| CZ-LE1-1^+^ | Western Eurasia | 48.7833 | 16.7899 | 9.83 |
| CZ-N1-1^+^ | Western Eurasia | 48.77532 | 16.72353 | 9.92 |
| CZ-N2-6^+^ | Western Eurasia | 48.76664 | 16.74248 | 9.93 |
| CZ-NVR-1 | Western Eurasia | 49.00661 | 14.44203 | 8.64 |
| DE_GB_1 | Western Eurasia | 54.32851 | 10.6274 | 8.71 |
| DE_SPO_1 | Western Eurasia | 54.2861 | 8.650398 | 8.77 |
| DE-G1-106^+^ | Western Eurasia | 54.28193 | 10.96651 | 8.65 |
| DE-K35-Mu10^+^ | Western Eurasia | 48.20665 | 11.70972 | 8.70 |
| DE-KA-28^+^ | Western Eurasia | 50.93536 | 6.927944 | 10.81 |
| DE-KN1-4^+^ | Western Eurasia | 54.17691 | 10.80668 | 8.37 |
| DE-R1-1^+^ | Western Eurasia | 54.2076 | 10.4236 | 8.43 |
| DE-S1-8^+^ | Western Eurasia | 48.77987 | 9.182967 | 9.73 |
| DE-S2-1^+^ | Western Eurasia | 48.80603 | 9.172572 | 9.35 |
| DE-S3-3^+^ | Western Eurasia | 48.80631 | 9.171758 | 9.35 |
| DK-RL-3^+^ | Western Eurasia | 55.96417 | 9.596389 | 7.37 |
| DZ-JV-2 | Western Eurasia | 36.88033 | 7.757 | 17.94 |
| EG_ELIAS_1 | Western Eurasia | 28.54411 | 33.97395 | 14.82 |
| ES-D-BDE1-F9 | Western Eurasia | 37.14807 | -6.03655 | 17.91 |
| ES-D-CC-NF7 | Western Eurasia | 37.07111 | -6.27241 | 18.10 |
| ES-D-H-F3 | Western Eurasia | 36.87183 | -5.86155 | 17.53 |
| ES-DO1-1 | Western Eurasia | 36.97836 | -6.47764 | 17.87 |
| ES-HT-1 | Western Eurasia | 38.77519 | -1.41022 | 13.08 |
| ES-HY-1 | Western Eurasia | 38.77931 | -1.435 | 13.05 |
| ES-OM-1 | Western Eurasia | 37.25435 | -6.96911 | 18.10 |
| ES-OM-5 | Western Eurasia | 37.25435 | -6.96911 | 18.10 |
| ES-RO-1 | Western Eurasia | 37.12793 | -6.4816 | 18.10 |
| ES-RV-1 | Western Eurasia | 40.65943 | 0.774614 | 16.78 |
| FI-FAT-1-3*^+^ | Western Eurasia | 60.02165 | 19.90212 | 5.83 |
| FI-FAV-1-1*^+^ | Western Eurasia | 60.02161 | 19.90251 | 5.83 |
| FI-FSP1-1*^+^ | Western Eurasia | 60.18436 | 25.79478 | 5.02 |
| FI-FUT1-2-1*^+^ | Western Eurasia | 60.34705 | 27.47848 | 4.53 |
| FI-OER-3-3*^+^ | Western Eurasia | 59.78859 | 23.17414 | 5.50 |
| FI-SK-17-1*^+^ | Western Eurasia | 59.83122 | 23.2577 | 5.50 |
| FI-SK-58-2c*^+^ | Western Eurasia | 59.83305 | 23.25743 | 5.50 |
| FI-SKW-2-1*^+^ | Western Eurasia | 59.83297 | 23.25626 | 5.50 |
| FR-C-1 | Western Eurasia | 43.59158 | 4.591517 | 14.97 |
| FR-LR5-1 | Western Eurasia | 43.51503 | 3.82309 | 14.98 |
| FR-LR6-1 | Western Eurasia | 43.4518 | 3.80755 | 15.14 |
| FR-LR7-1 | Western Eurasia | 43.49367 | 3.79486 | 15.03 |
| FR-LR9-1 | Western Eurasia | 43.56712 | 3.96982 | 14.82 |
| FR-NPM19-1^+^ | Western Eurasia | 46.99861 | -2.23944 | 12.77 |
| FR-NPM21-1^+^ | Western Eurasia | 46.99889 | -2.235 | 12.77 |
| FR-SA-1 | Western Eurasia | 43.48012 | 4.647302 | 14.95 |
| FR-TR-1 | Western Eurasia | 43.73831 | 3.863302 | 14.31 |
| GB-C1-1^+^ | Western Eurasia | 51.73441 | -1.33626 | 9.86 |
| GB-EK1-32^+^ | Western Eurasia | 55.70241 | -2.34083 | 8.15 |
| GB-EK2-6^+^ | Western Eurasia | 55.69766 | -2.34343 | 8.20 |
| GB-EP-1^+^ | Western Eurasia | 52.45358 | -1.91944 | 9.33 |
| GB-FML-1^+^ | Western Eurasia | 52.53106 | -1.95587 | 9.35 |
| GB-LK1-1^+^ | Western Eurasia | 55.70765 | -2.19038 | 8.28 |
| GB-S17-7^+^ | Western Eurasia | 51.62046 | -1.38622 | 10.05 |
| GR-K1-1 | Western Eurasia | 40.704 | 23.14422 | 14.49 |
| HU-AG-03 | Western Eurasia | 47.51458 | 19.08129 | 11.46 |
| HU-CR3-01 | Western Eurasia | 47.07374 | 19.13603 | 10.76 |
| HU-K-6^+^ | Western Eurasia | 46.79639 | 19.185 | 10.76 |
| HU-KEL-01 | Western Eurasia | 46.80277 | 19.16442 | 10.76 |
| IE_DUB_1 | Western Eurasia | 53.32674 | -6.23408 | 10.20 |
| IE-DUB-5 | Western Eurasia | 53.32674 | -6.23408 | 10.20 |
| IL-BM-1 | Western Eurasia | 30.51134 | 34.6121 | 16.41 |
| IL-EY-17 | Western Eurasia | 30.93801 | 35.04054 | 20.01 |
| IL-G-3 | Western Eurasia | 32.22884 | 34.83114 | 19.65 |
| IL-M1-8 | Western Eurasia | 31.77821 | 35.2206 | 16.25 |
| IL-PS-2 | Western Eurasia | 32.25571 | 34.85327 | 19.80 |
| IL-PS-6 | Western Eurasia | 32.25571 | 34.85327 | 19.80 |
| IL-RAM-4 | Western Eurasia | 33.23007 | 35.76511 | 15.60 |
| IL-SH-4 | Western Eurasia | 31.70103 | 34.71376 | 19.74 |
| IL-TY-10 | Western Eurasia | 32.24366 | 34.85448 | 19.81 |
| IL-YERU | Western Eurasia | 30.98986 | 34.89108 | 18.49 |
| IR-GG1-102 | Western Eurasia | 37.91898 | 46.707 | 8.86 |
| IT-ISR1-2 | Western Eurasia | 43.69194 | 10.28889 | 14.18 |
| IT-MDV-1 | Western Eurasia | 37.68547 | 12.61753 | 17.85 |
| IT-PER-2 | Western Eurasia | 37.5192 | 14.30735 | 14.13 |
| KG-SK-1 | Western Eurasia | 41.75573 | 75.23239 | -3.94 |
| MA-ES-3 | Western Eurasia | 31.49071 | -9.76443 | 17.83 |
| NL_LA_1 | Western Eurasia | 52.2154 | 5.185928 | 9.65 |
| NO-AA-1*^+^ | Western Eurasia | 60.05095 | 5.074448 | 7.59 |
| NO-F-1 | Western Eurasia | 63.58765 | 10.72897 | 5.38 |
| NO-LADE-1^+^ | Western Eurasia | 63.44899 | 10.4529 | 5.64 |
| NO-M3-1* | Western Eurasia | 59.09718 | 11.19665 | 6.73 |
| NO-RO-1*^+^ | Western Eurasia | 67.52741 | 12.12679 | 5.73 |
| NO-V-7*^+^ | Western Eurasia | 67.68696 | 12.67197 | 5.47 |
| NO-VADSO-2 | Western Eurasia | 70.06469 | 29.76428 | 6.73 |
| NO-VAR1-5* | Western Eurasia | 70.38769 | 31.09082 | 1.60 |
| NO-VAR4-3* | Western Eurasia | 70.4045 | 31.0644 | 1.55 |
| NO-VAR5-6* | Western Eurasia | 70.40545 | 31.06488 | 1.55 |
| NO-VAR6-5* | Western Eurasia | 70.40566 | 31.06724 | 1.64 |
| PL-KNP-4^+^ | Western Eurasia | 52.32272 | 20.73051 | 7.79 |
| PL-W1-1^+^ | Western Eurasia | 52.21092 | 20.99654 | 8.37 |
| PL-W2-1 | Western Eurasia | 52.23041 | 21.02841 | 8.35 |
| RO-DS-3 | Western Eurasia | 45.14712 | 29.67817 | 11.48 |
| RU_KA2_6 | Western Eurasia | 45.61639 | 45.31333 | 10.63 |
| RU_KA3_2 | Western Eurasia | 45.33 | 45.7275 | 10.79 |
| RU_KAI_105 | Western Eurasia | 45.59944 | 45.2975 | 10.61 |
| RU_MA21_2 | Western Eurasia | 46.45806 | 42.81778 | 9.98 |
| RU-AS2-1 | Western Eurasia | 46.30252 | 48.01356 | 10.18 |
| RU-B3-01 | Western Eurasia | 50.21602 | 46.86357 | 6.50 |
| RU-B5-1^+^ | Western Eurasia | 49.98643 | 46.69301 | 6.75 |
| RU-BN-BB6^+^ | Western Eurasia | 50.15552 | 43.3894 | 7.13 |
| RU-BOL1-1*^+^ | Western Eurasia | 66.42572 | 33.83683 | 0.72 |
| RU-BOR-1 | Western Eurasia | 58.07542 | 38.19938 | 3.78 |
| RU-BP-1^+^ | Western Eurasia | 58.07542 | 38.19938 | 3.78 |
| RU-C10-06 | Western Eurasia | 50.5007 | 47.15726 | 6.58 |
| RU-C12-1^+^ | Western Eurasia | 50.49643 | 47.20594 | 6.53 |
| RU-C20-1^+^ | Western Eurasia | 50.50979 | 46.51903 | 6.76 |
| RU-FR-LR8-1 | Western Eurasia | 43.51476 | 3.8334 | 14.95 |
| RU-KOR1-1*^+^ | Western Eurasia | 66.45189 | 33.79901 | 0.72 |
| RU-KUI-2^+^ | Western Eurasia | 55.30383 | 63.40533 | 2.29 |
| RU-MA10-3 | Western Eurasia | 46.49156 | 42.89798 | 9.82 |
| RU-MA13-3 | Western Eurasia | 46.44378 | 42.8091 | 9.97 |
| RU-MA20-10 | Western Eurasia | 46.54028 | 42.64028 | 9.80 |
| RU-MA23-8 | Western Eurasia | 46.55278 | 41.60306 | 9.86 |
| RU-MA3-5 | Western Eurasia | 46.46327 | 42.78464 | 9.99 |
| RU-MA4-3 | Western Eurasia | 46.43908 | 42.74524 | 10.00 |
| RU-MA8-01 | Western Eurasia | 46.47187 | 42.81004 | 9.93 |
| RU-NOV1-04 | Western Eurasia | 55.1278 | 77.03733 | 0.89 |
| RU-NOV2-01 | Western Eurasia | 53.75382 | 77.89404 | 1.70 |
| RU-NOV-3-2 | Western Eurasia | 55.03217 | 77.46633 | 0.94 |
| RU-NOV-4-7 | Western Eurasia | 53.774 | 78.00783 | 1.74 |
| RU-NOV5-17 | Western Eurasia | 53.74617 | 78.0525 | 1.84 |
| RU-P1-01 | Western Eurasia | 54.53611 | 39.79041 | 5.18 |
| RU-PIK-2^+^ | Western Eurasia | 59.99491 | 30.47308 | 4.87 |
| RU-R2-1^+^ | Western Eurasia | 56.425 | 37.60267 | 4.35 |
| RU-RM1-2^+^ | Western Eurasia | 55.76347 | 37.58158 | 5.47 |
| RU-RO1-3^+^ | Western Eurasia | 55.68332 | 39.88246 | 4.62 |
| RU-RT1-135^+^ | Western Eurasia | 45.22167 | 36.80833 | 11.55 |
| RU-RT21-1^+^ | Western Eurasia | 45.2221 | 36.6848 | 11.55 |
| RU-SAM2-1 | Western Eurasia | 52.08709 | 50.86831 | 5.55 |
| RU-SAM5-1 | Western Eurasia | 52.92296 | 50.31727 | 4.88 |
| RU-SPB-35^+^ | Western Eurasia | 59.81111 | 30.13306 | 4.86 |
| RU-SYR1-1^+^ | Western Eurasia | 53.21292 | 48.48821 | 5.25 |
| RU-TU1-1^+^ | Western Eurasia | 55.62433 | 69.20867 | 1.30 |
| RU-VOL-36 | Western Eurasia | 48.53 | 44.48694 | 8.31 |
| SE-BY-6^+^ | Western Eurasia | 55.67545 | 13.54507 | 7.76 |
| SE-G1-9c*^+^ | Western Eurasia | 60.42173 | 18.51015 | 5.50 |
| SE-G2-8c*^+^ | Western Eurasia | 60.43215 | 18.52237 | 5.50 |
| SE-G4-20*^+^ | Western Eurasia | 60.4152 | 18.56895 | 5.50 |
| SE-GN1-3d3*^+^ | Western Eurasia | 60.49717 | 18.43103 | 5.64 |
| SE-GN2-3A10*^+^ | Western Eurasia | 60.49713 | 18.43156 | 5.64 |
| SE-H1-4^+^ | Western Eurasia | 58.34233 | 11.21803 | 7.48 |
| TN-RA-2 | Western Eurasia | 36.96 | 10.22356 | 18.46 |
| TR-EG-1^+^ | Western Eurasia | 39.82361 | 32.83056 | 10.71 |
| UA-KRI-7^+^ | Western Eurasia | 45.09371 | 36.31065 | 11.54 |
| CA_TUK1_7* | North America | 69.44308 | -133.031 | -10.17 |
| CA_TUK2_4* | North America | 69.43722 | -133.016 | -9.96 |
| CA_YE1_1 | North America | 62.46889 | -114.495 | -5.03 |
| CA_YE2_6 | North America | 62.46861 | -114.496 | -5.03 |
| CA-CAL1-1 | North America | 50.93119 | -113.865 | 3.60 |
| CA-CAL2-1 | North America | 50.93656 | -113.869 | 3.63 |
| CA-CH-1* | North America | 58.77098 | -93.8508 | -6.74 |
| CA-LL1-1 | North America | 52.40856 | -105.006 | 1.38 |
| CA-LL2-2 | North America | 52.45734 | -105.049 | 1.57 |
| CA-SS-1 | North America | 51.56034 | -106.357 | 2.29 |
| US-D-1 | North America | 38.53014 | -121.611 | 15.93 |
| US-FAR1-1 | North America | 40.95826 | -111.916 | 10.74 |
| US-SP15-1* | North America | 44.3331 | -68.0629 | 6.73 |
| US-SP163-1* | North America | 44.3336 | -68.0636 | 6.91 |
| US-SP2-1* | North America | 44.33307 | -68.0629 | 6.73 |
| US-SP22-1-1* | North America | 44.3343 | -68.0648 | 6.91 |
| US-SP6-10* | North America | 44.33357 | -68.0635 | 6.91 |
| US-SP7-1* | North America | 44.33249 | -68.0612 | 6.73 |
| US-SU-1 | North America | 38.13773 | -121.908 | 15.33 |
| CN-DONG-1 | East Asia | 35.29854 | 98.67672 | -2.83 |
| CN-FUX-3 | East Asia | 24.63344 | 102.908 | 15.86 |
| CN-JIANG1-2 | East Asia | 34.61881 | 98.05581 | -3.25 |
| CN-JIANG3-1 | East Asia | 34.61914 | 98.05501 | -3.25 |
| CN-MADU1-1 | East Asia | 35.03803 | 97.3978 | -3.48 |
| CN-MADU3-1 | East Asia | 35.09329 | 97.76693 | -3.42 |
| CN-MADU4-2 | East Asia | 35.03306 | 98.05064 | -3.40 |
| CN-SHIQU1-3 | East Asia | 32.97159 | 98.33859 | 1.02 |
| CN-W1-1 | East Asia | 30.56237 | 114.3901 | 16.83 |
| CN-WON-1 | East Asia | 46.82502 | 124.379 | 4.10 |
| CN-XIU-1 | East Asia | 46.5276 | 124.7968 | 4.25 |
| CN-YUE-1 | East Asia | 49.37319 | 117.6624 | 0.45 |
| CN-ZHANG-1 | East Asia | 34.48546 | 118.7207 | 13.90 |
| CN-ZONG-1 | East Asia | 46.29601 | 125.0217 | 4.15 |
| MN-DM-1 | East Asia | 45.03271 | 100.6605 | 3.83 |
| RU_IRK1_10 | East Asia | 53.07415 | 106.9375 | -0.81 |
| RU_IRK5_72 | East Asia | 52.83539 | 106.5804 | -0.85 |
| RU_IRK9_1 | East Asia | 53.694 | 102.3513 | -1.10 |
| RU-AL1-11 | East Asia | 49.96783 | 88.692 | -3.94 |
| RU-BAL1-2 | East Asia | 53.01778 | 106.8858 | -0.60 |
| RU-BAYA1-1 | East Asia | 53.01783 | 106.8868 | -0.60 |
| RU-BU2-2 | East Asia | 51.315 | 108.4105 | -2.77 |
| RU-HA2-12 | East Asia | 54.67267 | 90.22683 | 0.50 |
| RU-IRK1-12 | East Asia | 53.07415 | 106.9375 | -0.81 |
| RU-IRK2-14 | East Asia | 52.98949 | 106.721 | -1.22 |
| RU-IRK3-1 | East Asia | 53.10517 | 107.2543 | -1.66 |
| RU-IRK4-10 | East Asia | 52.87683 | 106.5863 | -1.00 |
| RU-TY4-1 | East Asia | 50.1005 | 95.15017 | -3.71 |
| RU-YAK1-6 | East Asia | 61.96405 | 129.631 | -9.61 |
| RU-YAK3-25 | East Asia | 61.93746 | 129.6382 | -9.71 |
| RU-ZB1-1 | East Asia | 50.41317 | 114.7247 | -0.19 |
| RU-ZB2-22 | East Asia | 50.34718 | 114.8258 | -0.23 |
| RU-ZB4-12 | East Asia | 50.32433 | 115.0997 | -0.50 |
| RU-ZB5-1 | East Asia | 50.31817 | 115.3052 | -0.50 |
| RU-ZB6-03 | East Asia | 50.22585 | 115.6419 | -0.52 |

^1.^ L. Seefeldt, D. Ebert, Temperature- versus precipitation-limitation shape local temperature tolerance in a Holarctic freshwater crustacean. *Proc. R. Soc. B* **286**, 20190929 (2019).

### **Table S2.** Description of the 50 collagen genes identified in the D. magna genome. Genes included in the main analysis are highlighted with a “Y” (n=39), otherwise “N” (n=11) in “Main”. The coordinates based on the D. magna reference genome are displayed with the chromosome (Chr), contig number, and start and end position (Pos) information. The length and number of Gly-X_aa_-Y_aa_ repetitive motifs are shown per gene in the last column. In genes with multiple motifs (i.e., with a non-Gly-X_aa_-Y_aa_ signature in between) each region is listed. Overall, the regions of interest in the protein sequence (i.e., at least 20 Gly-X_aa_-Y_aa_ repetitive motifs) of the 50 collagen genes comprise 15726 bp, from which 12498 bp (i.e., from the 39 complete collagen genes) were included in the main analysis.

| **Gene Id** | **Main** | **Chr** | | **Contig** | | **Start Pos** | | **End Pos** | | **Repetitive motifs** |
| --- | --- | --- | --- | --- | --- | --- | --- | --- | --- | --- |
| maker-000008F-augustus-gene-18.25-mRNA-1 | Y | 1 | 000008F | | 1676224 | | 1688246 | | 21+25+35+44 | |
| augustus_masked-000010F-processed-gene-17.22-mRNA-1 | Y | 1 | 000010F | | 1858008 | | 1869057 | | 20 | |
| maker-000016F-augustus-gene-0.27-mRNA-1 | N | 1 | 000016F | | 37140 | | 42652 | | 61+63 | |
| maker-000000F-augustus-gene-29.50-mRNA-1 | Y | 2 | 000000F | | 831877 | | 836400 | | 338 | |
| maker-000004F-augustus-gene-34.44-mRNA-4 | Y | 2 | 000004F | | 2980082 | | 2991082 | | 37+24+52 | |
| maker-000004F-augustus-gene-34.47-mRNA-3 | N | 2 | 000004F | | 3384060 | | 3389473 | | 63+46 | |
| maker-000004F-augustus-gene-34.59-mRNA-1 | Y | 2 | 000004F | | 3403085 | | 3405604 | | 38+24+51 | |
| maker-000004F-augustus-gene-39.19-mRNA-1 | Y | 2 | 000004F | | 3444220 | | 3453295 | | 40+24+23+22+20+59+22+24+23+26 | |
| maker-000004F-augustus-gene-39.23-mRNA-1 | Y | 2 | 000004F | | 3903607 | | 3912448 | | 38+26+32+22+23+32+67+22+43+40+27 | |
| maker-000004F-snap-gene-34.24-mRNA-1 | Y | 2 | 000004F | | 3882307 | | 3891772 | | 37+24+51+37+24+51+24+42+37+24+36 | |
| maker-000004F-snap-gene-34.25-mRNA-1 | N | 2 | 000004F | | 3416718 | | 3419715 | | 37+24+51 | |
| maker-000004F-snap-gene-35.52-mRNA-1 | Y | 2 | 000004F | | 3421772 | | 3433551 | | 115+106+114 | |
| snap_masked-000004F-processed-gene-34.95-mRNA-1 | Y | 2 | 000004F | | 3565392 | | 3575641 | | 36+76 | |
| snap_masked-000004F-processed-gene-34.96-mRNA-1 | N | 2 | 000004F | | 888181 | | 890538 | | 37+24+52 | |
| augustus_masked-000036F-processed-gene-8.49-mRNA-1 | N | 2 | 000036F | | 818988 | | 820622 | | 37+24+50 | |
| maker-000036F-augustus-gene-8.91-mRNA-1 | Y | 2 | 000036F | | 822898 | | 826238 | | 37+24+51 | |
| maker-000036F-augustus-gene-8.94-mRNA-1 | Y | 2 | 000036F | | 828346 | | 830000 | | 37+24+51 | |
| maker-000036F-augustus-gene-8.95-mRNA-1 | Y | 2 | 000036F | | 3391531 | | 3393043 | | 37+94 | |
| maker-000036F-augustus-gene-8.96-mRNA-1 | Y | 2 | 000036F | | 3395679 | | 3397918 | | 37+24+51 | |
| maker-000011F-augustus-gene-19.87-mRNA-1 | Y | 4 | 000011F | | 1357865 | | 1362384 | | 33 | |
| snap_masked-000011F-processed-gene-20.3-mRNA-1 | Y | 4 | 000011F | | 798468 | | 802168 | | 20 | |
| augustus_masked-000028F-processed-gene-13.51-mRNA-1 | Y | 4 | 000028F | | 1960595 | | 1961936 | | 23+24 | |
| genemark-000028F-processed-gene-8.23-mRNA-1 | Y | 4 | 000028F | | 1330115 | | 1331202 | | 49 | |
| maker-000028F-augustus-gene-13.116-mRNA-1 | Y | 4 | 000028F | | 1353021 | | 1353928 | | 23 | |
| maker-000028F-augustus-gene-13.121-mRNA-1 | Y | 4 | 000028F | | 1332761 | | 1334940 | | 24 | |
| maker-000028F-snap-gene-13.68-mRNA-5 | Y | 4 | 000028F | | 2016669 | | 2017439 | | 25 | |
| snap_masked-000001F-processed-gene-14.25-mRNA-1 | Y | 5 | 000001F | | 113392 | | 125151 | | 199+138 | |
| maker-000108F-augustus-gene-0.59-mRNA-2 | Y | 5 | 000108F | | 1424939 | | 1434664 | | 23+23+36 | |
| maker-000012F-augustus-gene-19.14-mRNA-1 | Y | 6 | 000012F | | 1971241 | | 1973791 | | 34+22+42 | |
| maker-000012F-augustus-gene-21.72-mRNA-1 | Y | 6 | 000012F | | 2097919 | | 2105278 | | 37+24+50 | |
| maker-000082F-snap-gene-1.69-mRNA-1 | Y | 6 | 000082F | | 212269 | | 225294 | | 26+34+20 | |
| maker-000003F-augustus-gene-12.90-mRNA-1 | N | 7 | 000003F | | 238647 | | 245968 | | 23+20 | |
| maker-000003F-augustus-gene-4.97-mRNA-1 | Y | 7 | 000003F | | 4213 | | 11524 | | 48 | |
| maker-000003F-snap-gene-42.129-mRNA-1 | N | 7 | 000003F | | 222512 | | 228078 | | 30 | |
| maker-000018F-augustus-gene-4.32-mRNA-1 | Y | 7 | 000018F | | 1242510 | | 1243985 | | 47+25 | |
| augustus_masked-000038F-processed-gene-2.75-mRNA-1 | N | 7 | 000038F | | 402487 | | 411386 | | 25 | |
| genemark-000038F-processed-gene-2.57-mRNA-1 | Y | 7 | 000038F | | 4374221 | | 4381447 | | 28+30+30+29+30+30 | |
| maker-000038F-augustus-gene-2.132-mRNA-1 | Y | 7 | 000038F | | 458741 | | 466065 | | 30 | |
| maker-000038F-augustus-gene-2.134-mRNA-1 | Y | 7 | 000038F | | 229247 | | 230947 | | 32+32 | |
| maker-000038F-augustus-gene-2.135-mRNA-1 | N | 7 | 000038F | | 267057 | | 271057 | | 23 | |
| augustus_masked-000039F-processed-gene-0.41-mRNA-1 | Y | 7 | 000039F | | 272690 | | 274261 | | 30 | |
| maker-000031F-augustus-gene-2.77-mRNA-1 | Y | 8 | 000031F | | 199918 | | 203133 | | 76 | |
| maker-000031F-augustus-gene-3.25-mRNA-1 | Y | 8 | 000031F | | 368034 | | 372954 | | 91 | |
| genemark-000079F-processed-gene-0.29-mRNA-1 | Y | 9 | 000079F | | 86920 | | 89264 | | 83 | |
| maker-000079F-augustus-gene-0.96-mRNA-1 | Y | 9 | 000079F | | 75727 | | 79284 | | 87 | |
| maker-000079F-augustus-gene-0.98-mRNA-1 | N | 9 | 000079F | | 89325 | | 92914 | | 97 | |
| maker-000006F-augustus-gene-22.69-mRNA-1 | Y | 10 | 000006F | | 2269268 | | 2273000 | | 37+24+52 | |
| maker-000006F-augustus-gene-22.74-mRNA-1 | Y | 10 | 000006F | | 2242736 | | 2246726 | | 37+24+40 | |
| maker-000006F-augustus-gene-22.75-mRNA-1 | Y | 10 | 000006F | | 2246895 | | 2248537 | | 37+24+50 | |
| maker-000035F-snap-gene-5.102-mRNA-1 | N | 10 | 000035F | | 524755 | | 528309 | | 37+32 | |

### **Table S3**. Amino acid content (counts per 1000 residues) for each D. magna genotype, averaged among Gly-X_aa_-Y_aa_ motifs of the 39 collagen genes and phased haplotypes.

| **Genotype** | **Ala** | **Arg** | **Asn** | **Asp** | **Cys** | **Glu** | **Gln** | **Gly** | **His** | **Ile** | **Leu** | **Lys** | **Met** | **Phe** | **Pro** | **Ser** | **Thr** | **Trp** | **Tyr** | **Val** |
| --- | --- | --- | --- | --- | --- | --- | --- | --- | --- | --- | --- | --- | --- | --- | --- | --- | --- | --- | --- | --- |
| AM-AR1-1 | 71.6 | 22.6 | 21.1 | 44.4 | 0.2 | 29.4 | 26.3 | 341.5 | 2.0 | 34.2 | 14.4 | 44.5 | 7.7 | 10.1 | 212.0 | 28.0 | 31.5 | 2.3 | 13.8 | 20.6 |
| BE-HO-1 | 72.2 | 22.7 | 21.2 | 44.4 | 0.3 | 29.2 | 26.3 | 341.4 | 2.1 | 34.1 | 14.3 | 44.5 | 7.8 | 10.0 | 211.8 | 27.9 | 31.4 | 2.3 | 13.8 | 20.5 |
| BE-KN2-1 | 71.1 | 22.8 | 21.5 | 44.2 | 0.3 | 29.5 | 26.1 | 341.4 | 2.0 | 34.1 | 14.2 | 44.4 | 7.8 | 10.1 | 211.8 | 28.3 | 31.7 | 2.3 | 13.7 | 20.8 |
| BE-OHZ-T10 | 72.9 | 22.7 | 20.8 | 44.9 | 0.2 | 28.9 | 26.3 | 341.8 | 2.1 | 34.1 | 14.2 | 44.5 | 8.0 | 10.3 | 211.3 | 28.0 | 31.1 | 2.3 | 13.7 | 20.3 |
| BE-OM-1 | 72.0 | 22.8 | 21.2 | 44.3 | 0.2 | 29.3 | 26.1 | 341.6 | 2.2 | 34.1 | 14.5 | 44.5 | 7.9 | 10.0 | 211.8 | 28.1 | 31.3 | 2.3 | 13.7 | 20.3 |
| BE-T1-2 | 72.5 | 22.6 | 20.6 | 45.0 | 0.3 | 29.0 | 26.4 | 341.5 | 2.2 | 34.0 | 14.3 | 44.5 | 7.8 | 10.2 | 211.7 | 27.9 | 31.0 | 2.3 | 13.7 | 20.6 |
| BE-WE-59 | 72.2 | 22.8 | 21.1 | 44.5 | 0.2 | 29.3 | 26.3 | 341.8 | 2.1 | 34.1 | 14.4 | 44.5 | 7.8 | 10.0 | 211.6 | 27.9 | 31.2 | 2.3 | 13.7 | 20.3 |
| BE-WH1-2 | 72.2 | 22.6 | 20.7 | 44.7 | 0.3 | 29.2 | 26.3 | 341.3 | 2.0 | 34.1 | 14.1 | 44.5 | 7.9 | 10.0 | 211.5 | 28.6 | 31.2 | 2.3 | 13.9 | 20.9 |
| BE-WH2-11 | 71.9 | 22.7 | 21.0 | 44.3 | 0.2 | 29.3 | 26.3 | 341.5 | 2.2 | 34.0 | 14.3 | 44.6 | 7.9 | 10.0 | 211.7 | 28.3 | 31.5 | 2.3 | 13.8 | 20.5 |
| BE-WTE-1-1 | 71.7 | 22.7 | 21.7 | 43.8 | 0.3 | 29.5 | 26.2 | 341.6 | 2.0 | 34.1 | 14.4 | 44.5 | 7.8 | 10.0 | 211.9 | 27.9 | 31.4 | 2.3 | 13.8 | 20.5 |
| BY-G-9 | 72.3 | 22.6 | 21.2 | 44.5 | 0.2 | 29.1 | 26.2 | 341.7 | 2.2 | 34.4 | 14.3 | 44.6 | 7.8 | 10.0 | 210.8 | 28.3 | 31.6 | 2.3 | 13.7 | 20.3 |
| CA-CAL1-1 | 73.3 | 22.9 | 20.8 | 45.2 | 0.2 | 28.6 | 26.6 | 341.3 | 2.2 | 33.8 | 14.4 | 44.2 | 8.0 | 10.4 | 209.8 | 28.7 | 31.5 | 2.3 | 13.7 | 20.1 |
| CA-CAL2-1 | 73.4 | 22.9 | 20.8 | 45.3 | 0.2 | 28.6 | 26.6 | 341.3 | 2.3 | 33.8 | 14.4 | 44.2 | 8.0 | 10.4 | 209.6 | 28.7 | 31.7 | 2.3 | 13.7 | 20.2 |
| CA-CH-1 | 73.4 | 22.9 | 20.7 | 45.0 | 0.2 | 28.9 | 26.6 | 341.2 | 2.2 | 33.8 | 14.5 | 44.2 | 8.0 | 10.4 | 209.9 | 28.7 | 31.4 | 2.3 | 13.7 | 20.2 |
| CA-LL1-1 | 73.4 | 22.9 | 20.8 | 45.1 | 0.2 | 28.8 | 26.6 | 341.3 | 2.3 | 33.8 | 14.5 | 44.1 | 8.0 | 10.4 | 209.5 | 28.9 | 31.5 | 2.3 | 13.7 | 20.1 |
| CA-LL2-2 | 73.2 | 22.9 | 20.8 | 45.0 | 0.2 | 28.8 | 26.5 | 341.3 | 2.3 | 33.8 | 14.6 | 44.2 | 8.0 | 10.4 | 209.6 | 28.8 | 31.5 | 2.3 | 13.7 | 20.3 |
| CA-RI-1 | 72.4 | 22.7 | 20.7 | 45.0 | 0.2 | 28.8 | 26.1 | 341.7 | 2.2 | 34.2 | 14.2 | 44.7 | 7.7 | 10.2 | 210.9 | 28.6 | 31.5 | 2.3 | 13.7 | 20.5 |
| CA-SS-1 | 73.4 | 22.9 | 20.7 | 45.1 | 0.2 | 28.9 | 26.6 | 341.2 | 2.3 | 33.8 | 14.6 | 44.1 | 8.0 | 10.4 | 209.6 | 28.8 | 31.5 | 2.3 | 13.7 | 20.1 |
| CA_TUK1_7 | 73.4 | 22.9 | 21.0 | 45.1 | 0.2 | 28.5 | 26.3 | 341.3 | 2.3 | 33.8 | 14.5 | 44.2 | 8.0 | 10.4 | 210.2 | 28.8 | 31.3 | 2.3 | 13.7 | 20.1 |
| CA_TUK2_4. | 73.4 | 22.9 | 20.7 | 45.4 | 0.2 | 28.5 | 26.5 | 341.3 | 2.3 | 33.8 | 14.4 | 44.1 | 8.0 | 10.4 | 210.0 | 28.7 | 31.4 | 2.3 | 13.7 | 20.1 |
| CA_YE1_1 | 73.4 | 22.9 | 20.8 | 45.1 | 0.2 | 28.8 | 26.6 | 341.3 | 2.3 | 33.9 | 14.4 | 44.2 | 8.0 | 10.3 | 209.5 | 28.8 | 31.5 | 2.3 | 13.7 | 20.2 |
| CA_YE2_6 | 73.3 | 22.9 | 20.7 | 45.2 | 0.2 | 28.7 | 26.6 | 341.3 | 2.2 | 33.8 | 14.5 | 44.2 | 8.0 | 10.4 | 209.6 | 28.6 | 31.6 | 2.3 | 13.7 | 20.3 |
| CH-H-1 | 72.2 | 22.7 | 20.9 | 44.6 | 0.2 | 29.0 | 26.2 | 341.7 | 2.1 | 33.9 | 14.2 | 44.6 | 7.8 | 10.0 | 211.5 | 28.1 | 31.7 | 2.3 | 13.7 | 20.8 |
| CH-Z-1 | 72.1 | 22.7 | 20.8 | 44.8 | 0.3 | 29.1 | 26.3 | 341.7 | 2.1 | 33.9 | 14.1 | 44.4 | 7.8 | 10.0 | 211.3 | 28.4 | 31.3 | 2.3 | 13.7 | 21.1 |
| CN-DONG-1 | 72.5 | 22.6 | 21.5 | 44.3 | 0.2 | 29.0 | 26.4 | 341.6 | 2.1 | 33.6 | 14.7 | 44.4 | 7.7 | 10.4 | 211.5 | 28.0 | 31.4 | 2.3 | 13.7 | 20.4 |
| CN-FUX-3 | 72.4 | 22.6 | 21.3 | 44.2 | 0.2 | 29.5 | 26.2 | 341.4 | 2.1 | 33.9 | 14.6 | 44.5 | 7.7 | 10.2 | 212.5 | 27.5 | 30.9 | 2.3 | 13.6 | 20.6 |
| CN-JIANG1-2 | 72.6 | 22.6 | 21.6 | 44.0 | 0.2 | 29.0 | 26.4 | 341.5 | 2.1 | 34.2 | 14.7 | 44.4 | 7.6 | 10.4 | 211.0 | 28.1 | 31.6 | 2.3 | 13.7 | 20.2 |
| CN-JIANG2-3 | 72.4 | 22.6 | 20.9 | 44.6 | 0.2 | 29.0 | 26.4 | 341.5 | 2.1 | 33.8 | 14.3 | 44.6 | 7.7 | 10.2 | 211.2 | 28.2 | 31.7 | 2.3 | 13.7 | 20.8 |
| CN-JIANG3-1 | 72.4 | 22.6 | 21.6 | 44.1 | 0.2 | 29.0 | 26.3 | 341.6 | 2.3 | 34.2 | 14.6 | 44.4 | 7.5 | 10.3 | 211.3 | 28.1 | 31.6 | 2.3 | 13.7 | 20.2 |
| CN-MADU1-1 | 73.4 | 22.7 | 21.1 | 44.7 | 0.2 | 28.9 | 26.2 | 341.6 | 2.2 | 33.9 | 14.7 | 44.4 | 7.6 | 10.2 | 211.1 | 28.0 | 31.4 | 2.3 | 13.7 | 20.1 |
| CN-MADU3-1 | 72.5 | 22.7 | 21.5 | 44.4 | 0.2 | 28.9 | 26.3 | 341.6 | 2.2 | 34.1 | 14.6 | 44.4 | 7.7 | 10.4 | 210.6 | 27.9 | 31.8 | 2.3 | 13.7 | 20.5 |
| CN-MADU4-2 | 72.7 | 22.6 | 21.4 | 44.4 | 0.2 | 29.1 | 26.3 | 341.5 | 2.2 | 34.0 | 14.8 | 44.4 | 7.4 | 10.4 | 210.6 | 27.9 | 31.9 | 2.3 | 13.7 | 20.3 |
| CN-SHIQU1-3 | 72.7 | 22.7 | 21.2 | 44.2 | 0.2 | 29.1 | 26.5 | 341.6 | 2.1 | 34.0 | 14.9 | 44.4 | 7.8 | 10.4 | 211.3 | 27.4 | 31.6 | 2.3 | 13.6 | 20.2 |
| CN-W1-1 | 72.5 | 22.8 | 21.7 | 44.2 | 0.2 | 29.1 | 26.2 | 341.5 | 2.1 | 33.8 | 14.5 | 44.3 | 7.8 | 10.2 | 212.6 | 27.4 | 30.9 | 2.3 | 13.6 | 20.5 |
| CN-WON-1 | 72.7 | 22.7 | 20.8 | 44.9 | 0.2 | 28.7 | 26.3 | 341.6 | 2.1 | 33.9 | 14.6 | 44.6 | 7.7 | 10.2 | 212.0 | 27.4 | 31.8 | 2.3 | 13.7 | 20.1 |
| CN-XIU-1 | 72.7 | 22.6 | 20.7 | 45.0 | 0.2 | 29.0 | 26.2 | 341.6 | 2.1 | 33.8 | 14.6 | 44.6 | 7.8 | 10.3 | 211.8 | 27.5 | 31.8 | 2.3 | 13.6 | 20.0 |
| CN-YUE-1 | 72.6 | 22.9 | 21.4 | 44.3 | 0.2 | 29.2 | 26.1 | 341.6 | 2.2 | 34.1 | 14.6 | 44.5 | 7.8 | 10.3 | 211.4 | 27.6 | 31.4 | 2.3 | 13.7 | 20.2 |
| CN-ZHANG-1 | 72.4 | 22.6 | 21.3 | 44.3 | 0.2 | 29.3 | 26.2 | 341.4 | 2.1 | 33.9 | 14.5 | 44.5 | 7.8 | 10.2 | 212.9 | 27.3 | 30.9 | 2.3 | 13.6 | 20.5 |
| CN-ZONG-1 | 72.4 | 22.7 | 20.8 | 44.7 | 0.2 | 29.1 | 26.3 | 341.6 | 2.0 | 34.0 | 14.7 | 44.5 | 7.7 | 10.2 | 212.3 | 27.5 | 31.4 | 2.3 | 13.6 | 20.1 |
| CY-PA1-1 | 71.8 | 22.6 | 21.1 | 44.4 | 0.2 | 29.2 | 26.1 | 341.5 | 2.1 | 33.9 | 14.4 | 44.4 | 7.6 | 10.2 | 211.8 | 28.2 | 31.8 | 2.3 | 13.7 | 20.7 |
| CY-PA2-1 | 72.2 | 22.6 | 21.0 | 44.8 | 0.2 | 29.1 | 26.1 | 341.4 | 2.0 | 34.0 | 14.1 | 44.5 | 7.6 | 10.1 | 211.7 | 28.1 | 31.8 | 2.3 | 13.7 | 21.0 |
| CY-PA3-1 | 71.5 | 22.7 | 21.2 | 44.3 | 0.2 | 29.4 | 26.1 | 341.5 | 2.1 | 34.0 | 14.5 | 44.5 | 7.7 | 10.3 | 211.9 | 28.1 | 31.6 | 2.3 | 13.6 | 20.7 |
| CY-PA3-2 | 71.9 | 22.6 | 21.2 | 44.4 | 0.2 | 29.3 | 26.2 | 341.5 | 2.0 | 34.1 | 14.1 | 44.4 | 7.6 | 10.2 | 211.8 | 27.9 | 31.7 | 2.3 | 13.6 | 21.0 |
| CZ-KO-1 | 71.7 | 22.7 | 21.6 | 43.9 | 0.2 | 29.5 | 26.2 | 341.7 | 2.1 | 34.0 | 14.1 | 44.4 | 7.8 | 10.1 | 212.1 | 27.7 | 31.2 | 2.3 | 13.8 | 20.9 |
| CZ-LE1-1 | 72.5 | 22.6 | 20.7 | 44.7 | 0.3 | 29.0 | 26.1 | 341.5 | 2.2 | 34.1 | 14.0 | 44.6 | 7.8 | 10.1 | 211.6 | 28.2 | 31.4 | 2.3 | 13.8 | 20.7 |
| CZ-N1-1 | 71.7 | 22.8 | 21.4 | 44.0 | 0.3 | 29.3 | 26.2 | 341.5 | 2.1 | 34.0 | 14.4 | 44.5 | 7.8 | 9.9 | 211.9 | 28.2 | 31.3 | 2.3 | 13.8 | 20.7 |
| CZ-N2-6 | 72.7 | 22.7 | 20.9 | 44.6 | 0.3 | 29.0 | 26.1 | 341.5 | 2.0 | 34.1 | 14.2 | 44.6 | 7.9 | 10.1 | 211.3 | 28.2 | 31.4 | 2.3 | 13.8 | 20.6 |
| CZ-NVR-1 | 72.3 | 22.7 | 21.1 | 44.3 | 0.2 | 29.3 | 26.2 | 341.7 | 2.3 | 34.1 | 14.3 | 44.6 | 7.9 | 10.1 | 211.3 | 27.9 | 31.4 | 2.3 | 13.8 | 20.5 |
| CZ_DO_1 | 72.1 | 22.6 | 20.7 | 44.8 | 0.2 | 29.1 | 26.3 | 341.6 | 2.1 | 34.0 | 14.4 | 44.7 | 7.8 | 10.1 | 211.2 | 28.2 | 31.5 | 2.3 | 13.9 | 20.7 |
| CZ_SUR_1 | 72.4 | 22.5 | 21.1 | 44.3 | 0.2 | 29.3 | 26.1 | 341.7 | 2.2 | 34.1 | 14.3 | 44.5 | 7.8 | 10.2 | 211.6 | 27.7 | 31.4 | 2.3 | 13.8 | 20.6 |
| DE-G1-106 | 71.0 | 22.8 | 21.4 | 43.9 | 0.2 | 29.5 | 26.3 | 341.7 | 2.1 | 34.0 | 14.4 | 44.4 | 7.8 | 10.1 | 211.6 | 28.4 | 31.7 | 2.3 | 13.8 | 20.8 |
| DE-K35-Mu10 | 72.3 | 22.6 | 20.9 | 44.8 | 0.2 | 29.0 | 26.2 | 341.7 | 2.1 | 34.1 | 14.0 | 44.6 | 7.8 | 10.1 | 211.0 | 28.1 | 32.0 | 2.3 | 13.8 | 20.7 |
| DE-KA-28 | 72.3 | 22.6 | 20.7 | 44.7 | 0.3 | 29.1 | 26.3 | 341.6 | 2.2 | 34.0 | 14.0 | 44.5 | 7.7 | 10.1 | 212.0 | 27.9 | 31.6 | 2.3 | 13.8 | 20.6 |
| DE-KN1-4 | 72.0 | 22.6 | 20.7 | 44.6 | 0.3 | 29.0 | 26.2 | 341.5 | 2.2 | 33.9 | 14.3 | 44.8 | 7.8 | 10.0 | 211.5 | 28.1 | 31.6 | 2.3 | 13.8 | 20.8 |
| DE-R1-1 | 71.8 | 22.8 | 21.5 | 44.2 | 0.3 | 29.2 | 26.4 | 341.6 | 2.1 | 34.0 | 14.4 | 44.4 | 7.9 | 10.1 | 211.7 | 27.8 | 31.5 | 2.3 | 13.7 | 20.7 |
| DE-S1-8 | 72.2 | 22.5 | 21.2 | 44.4 | 0.2 | 29.1 | 26.3 | 341.7 | 2.1 | 33.9 | 14.3 | 44.6 | 7.7 | 10.0 | 211.6 | 28.0 | 31.6 | 2.3 | 13.7 | 20.6 |
| DE-S2-1 | 72.7 | 22.7 | 21.2 | 44.4 | 0.2 | 29.1 | 26.1 | 341.7 | 2.1 | 34.1 | 14.1 | 44.5 | 7.7 | 10.1 | 211.3 | 28.0 | 31.7 | 2.3 | 13.8 | 20.5 |
| DE-S3-3 | 72.7 | 22.5 | 21.3 | 44.3 | 0.2 | 29.1 | 26.2 | 341.7 | 2.1 | 34.0 | 14.2 | 44.6 | 7.8 | 10.0 | 211.8 | 27.8 | 31.4 | 2.3 | 13.7 | 20.4 |
| DE_GB_1 | 71.7 | 22.9 | 21.1 | 44.2 | 0.2 | 29.3 | 26.0 | 341.6 | 2.1 | 34.1 | 14.2 | 44.7 | 7.8 | 10.0 | 211.7 | 28.3 | 31.6 | 2.3 | 13.8 | 20.6 |
| DE_SPO_1 | 71.7 | 22.8 | 21.1 | 44.5 | 0.2 | 29.3 | 26.5 | 341.7 | 2.1 | 33.8 | 14.4 | 44.2 | 7.8 | 10.1 | 211.8 | 28.4 | 31.3 | 2.3 | 13.8 | 20.4 |
| DK-RL-3 | 71.7 | 22.6 | 21.4 | 44.0 | 0.2 | 29.5 | 26.4 | 341.6 | 2.0 | 34.1 | 14.4 | 44.6 | 7.8 | 10.1 | 211.7 | 28.1 | 31.4 | 2.3 | 13.8 | 20.5 |
| DZ-JV-2 | 71.1 | 22.6 | 21.6 | 43.6 | 0.2 | 29.4 | 26.1 | 341.5 | 2.2 | 34.1 | 14.6 | 44.7 | 7.8 | 10.0 | 212.3 | 28.0 | 31.5 | 2.3 | 13.7 | 20.8 |
| EG_ELIAS_1 | 72.3 | 22.6 | 20.9 | 44.5 | 0.2 | 29.0 | 26.4 | 341.6 | 2.0 | 33.9 | 14.1 | 44.5 | 7.6 | 10.0 | 211.5 | 28.3 | 31.9 | 2.3 | 13.6 | 20.9 |
| ES-D-BDE1-F9 | 72.4 | 22.7 | 21.0 | 44.5 | 0.2 | 29.0 | 26.1 | 341.5 | 2.2 | 34.2 | 14.4 | 44.7 | 7.5 | 10.0 | 211.8 | 27.9 | 31.7 | 2.3 | 13.8 | 20.5 |
| ES-D-CC-NF7 | 71.6 | 22.5 | 21.4 | 44.0 | 0.2 | 29.5 | 26.0 | 341.8 | 2.0 | 34.4 | 14.2 | 44.7 | 7.7 | 10.1 | 212.8 | 27.5 | 31.4 | 2.3 | 13.6 | 20.5 |
| ES-D-H-F3 | 72.4 | 22.6 | 21.0 | 44.5 | 0.2 | 29.1 | 26.1 | 341.7 | 2.2 | 34.0 | 14.5 | 44.7 | 7.7 | 10.2 | 212.1 | 27.5 | 31.5 | 2.3 | 13.6 | 20.3 |
| ES-DO1-1 | 71.4 | 22.6 | 21.3 | 43.9 | 0.2 | 29.6 | 26.2 | 341.6 | 2.1 | 34.2 | 14.5 | 44.7 | 7.7 | 10.1 | 212.3 | 27.9 | 31.5 | 2.3 | 13.5 | 20.4 |
| ES-HT-1 | 71.8 | 22.6 | 21.2 | 44.1 | 0.2 | 29.2 | 25.8 | 341.6 | 2.0 | 34.1 | 14.4 | 44.8 | 7.8 | 10.0 | 212.4 | 27.7 | 31.6 | 2.3 | 13.8 | 20.6 |
| ES-HY-1 | 72.3 | 22.5 | 20.9 | 44.3 | 0.2 | 29.2 | 25.9 | 341.7 | 2.1 | 34.1 | 14.4 | 44.6 | 7.7 | 10.0 | 212.2 | 28.0 | 31.3 | 2.3 | 13.7 | 20.7 |
| ES-OM-1 | 72.2 | 22.6 | 21.1 | 44.3 | 0.2 | 29.3 | 26.2 | 341.7 | 2.0 | 34.0 | 14.4 | 44.7 | 7.6 | 10.1 | 212.3 | 27.8 | 31.5 | 2.3 | 13.6 | 20.3 |
| ES-OM-5 | 71.8 | 22.6 | 21.2 | 44.3 | 0.2 | 29.3 | 26.1 | 341.7 | 2.1 | 34.1 | 14.5 | 44.7 | 7.6 | 10.2 | 212.0 | 27.7 | 31.8 | 2.3 | 13.7 | 20.5 |
| ES-RO-1 | 71.5 | 22.5 | 21.2 | 44.1 | 0.2 | 29.5 | 25.8 | 341.5 | 2.1 | 34.4 | 14.3 | 44.9 | 7.7 | 10.0 | 212.3 | 27.9 | 31.8 | 2.3 | 13.7 | 20.6 |
| ES-RV-1 | 71.6 | 22.6 | 21.5 | 43.9 | 0.2 | 29.5 | 26.1 | 341.5 | 2.1 | 34.2 | 14.5 | 44.7 | 7.8 | 10.0 | 212.0 | 27.8 | 31.6 | 2.3 | 13.7 | 20.6 |
| ET-C-1 | 72.2 | 22.6 | 20.9 | 44.6 | 0.2 | 29.2 | 26.4 | 341.5 | 2.1 | 33.8 | 14.3 | 44.6 | 7.7 | 10.2 | 211.2 | 28.2 | 31.7 | 2.3 | 13.7 | 20.8 |
| FI-FAT-1-3 | 71.6 | 22.8 | 21.0 | 44.4 | 0.2 | 29.3 | 26.0 | 341.6 | 2.1 | 34.2 | 14.3 | 44.6 | 7.9 | 10.1 | 211.5 | 28.1 | 31.6 | 2.3 | 13.7 | 20.9 |
| FI-FAV-1-1 | 71.7 | 22.7 | 21.0 | 44.6 | 0.2 | 29.2 | 26.1 | 341.6 | 2.1 | 34.0 | 14.3 | 44.6 | 7.8 | 10.2 | 211.2 | 28.7 | 31.5 | 2.3 | 13.8 | 20.7 |
| FI-FSE-3-74 | 71.5 | 22.6 | 21.4 | 44.3 | 0.2 | 29.3 | 26.2 | 341.6 | 2.0 | 34.4 | 14.2 | 44.6 | 7.7 | 10.0 | 211.2 | 28.3 | 31.7 | 2.3 | 13.7 | 20.8 |
| FI-FSP1-1 | 71.9 | 22.6 | 21.1 | 44.3 | 0.2 | 29.2 | 26.0 | 341.6 | 2.1 | 34.0 | 14.4 | 44.6 | 7.9 | 10.2 | 210.9 | 28.5 | 31.5 | 2.3 | 13.8 | 20.8 |
| FI-FUT1-2-1 | 72.0 | 22.7 | 20.8 | 44.7 | 0.2 | 29.1 | 26.2 | 341.6 | 2.2 | 34.1 | 14.0 | 44.6 | 8.0 | 10.2 | 211.2 | 28.2 | 31.6 | 2.3 | 13.7 | 20.8 |
| FI-OER-3-3 | 71.2 | 22.7 | 21.1 | 44.3 | 0.2 | 29.5 | 26.4 | 341.6 | 2.1 | 33.8 | 14.5 | 44.5 | 7.8 | 10.4 | 211.2 | 28.4 | 31.8 | 2.3 | 13.7 | 20.8 |
| FI-SK-17-1 | 71.8 | 22.6 | 20.8 | 44.8 | 0.2 | 29.0 | 26.1 | 341.6 | 2.1 | 34.1 | 14.4 | 44.7 | 7.8 | 10.2 | 211.0 | 28.5 | 31.6 | 2.3 | 13.7 | 20.7 |
| FI-SK-58-2c | 72.0 | 22.6 | 20.8 | 44.8 | 0.2 | 28.9 | 26.3 | 341.6 | 2.1 | 33.9 | 14.4 | 44.7 | 7.9 | 10.1 | 211.1 | 28.5 | 31.5 | 2.3 | 13.7 | 20.8 |
| FI-SKW-2-1 | 72.1 | 22.6 | 20.7 | 45.0 | 0.2 | 28.9 | 26.3 | 341.6 | 2.1 | 34.2 | 14.4 | 44.6 | 7.9 | 10.1 | 210.9 | 28.3 | 31.4 | 2.3 | 13.7 | 20.8 |
| FR-C-1 | 72.3 | 22.8 | 20.7 | 44.7 | 0.3 | 29.0 | 26.2 | 341.6 | 2.2 | 34.0 | 14.6 | 44.5 | 7.7 | 10.0 | 211.7 | 28.1 | 31.5 | 2.3 | 13.8 | 20.4 |
| FR-LR5-1 | 71.4 | 23.0 | 21.5 | 44.1 | 0.3 | 29.3 | 25.9 | 341.5 | 2.2 | 34.3 | 14.5 | 44.6 | 7.7 | 10.1 | 211.8 | 27.9 | 31.6 | 2.3 | 13.8 | 20.7 |
| FR-LR6-1 | 71.8 | 22.8 | 21.6 | 43.8 | 0.3 | 29.5 | 26.2 | 341.3 | 2.1 | 34.1 | 14.4 | 44.6 | 7.8 | 10.1 | 212.0 | 28.0 | 31.2 | 2.3 | 13.5 | 20.7 |
| FR-LR7-1 | 71.8 | 22.8 | 21.7 | 43.9 | 0.3 | 29.4 | 26.0 | 341.4 | 2.0 | 34.1 | 14.4 | 44.7 | 7.7 | 10.0 | 211.9 | 27.9 | 31.4 | 2.3 | 13.6 | 20.6 |
| FR-LR9-1 | 71.6 | 22.8 | 21.4 | 44.0 | 0.3 | 29.3 | 26.0 | 341.5 | 2.2 | 34.1 | 14.5 | 44.7 | 7.7 | 10.1 | 212.3 | 27.7 | 31.5 | 2.3 | 13.8 | 20.6 |
| FR-NPM19-1 | 71.4 | 22.7 | 21.3 | 44.0 | 0.3 | 29.3 | 26.1 | 341.5 | 2.0 | 34.0 | 14.7 | 44.6 | 7.8 | 10.0 | 212.3 | 28.1 | 31.4 | 2.3 | 13.8 | 20.6 |
| FR-NPM21-1 | 71.8 | 22.7 | 21.0 | 44.4 | 0.2 | 29.2 | 26.2 | 341.8 | 2.0 | 34.1 | 14.6 | 44.5 | 7.8 | 10.0 | 211.9 | 28.1 | 31.4 | 2.3 | 13.8 | 20.5 |
| FR-SA-1 | 71.8 | 22.6 | 21.1 | 44.3 | 0.3 | 29.3 | 25.8 | 341.3 | 2.1 | 34.3 | 14.3 | 44.6 | 7.8 | 10.0 | 211.6 | 28.4 | 31.6 | 2.3 | 13.7 | 20.8 |
| FR-TR-1 | 72.0 | 22.6 | 21.1 | 44.2 | 0.3 | 29.2 | 26.2 | 341.6 | 2.2 | 34.4 | 14.4 | 44.6 | 7.8 | 10.0 | 211.6 | 28.2 | 31.3 | 2.3 | 13.7 | 20.4 |
| GB-C1-1 | 71.7 | 22.6 | 21.3 | 44.4 | 0.2 | 29.3 | 26.1 | 341.7 | 2.1 | 34.2 | 14.5 | 44.5 | 8.0 | 10.1 | 212.2 | 28.0 | 31.2 | 2.3 | 13.8 | 20.1 |
| GB-EK1-1 | 72.2 | 22.7 | 20.8 | 44.6 | 0.3 | 29.1 | 26.3 | 341.7 | 2.1 | 33.9 | 14.2 | 44.4 | 7.8 | 10.1 | 211.8 | 28.2 | 31.3 | 2.3 | 13.8 | 20.6 |
| GB-EK1-32 | 71.7 | 22.6 | 20.6 | 44.6 | 0.2 | 29.4 | 26.3 | 341.6 | 2.2 | 34.0 | 14.5 | 44.7 | 7.9 | 10.1 | 212.3 | 27.8 | 31.2 | 2.3 | 13.8 | 20.4 |
| GB-EK2-6 | 72.0 | 22.7 | 20.9 | 44.5 | 0.2 | 29.2 | 26.3 | 341.7 | 2.1 | 33.9 | 14.5 | 44.6 | 7.9 | 10.0 | 212.1 | 27.9 | 31.2 | 2.3 | 13.8 | 20.5 |
| GB-EL75-69 | 71.2 | 22.8 | 21.5 | 44.2 | 0.3 | 29.5 | 26.1 | 341.4 | 2.0 | 34.1 | 14.2 | 44.4 | 7.8 | 10.1 | 211.8 | 28.3 | 31.6 | 2.3 | 13.7 | 20.8 |
| GB-EP-1 | 72.3 | 22.7 | 20.8 | 44.7 | 0.3 | 29.2 | 26.2 | 341.7 | 2.2 | 34.2 | 14.3 | 44.6 | 7.8 | 10.0 | 211.4 | 28.0 | 31.1 | 2.3 | 13.8 | 20.6 |
| GB-FML-1 | 71.9 | 22.7 | 21.1 | 44.4 | 0.2 | 29.3 | 26.2 | 341.8 | 2.2 | 34.0 | 14.3 | 44.6 | 7.8 | 10.2 | 211.9 | 27.8 | 31.0 | 2.3 | 13.8 | 20.8 |
| GB-LK1-1 | 72.0 | 22.5 | 20.9 | 44.6 | 0.2 | 29.2 | 26.3 | 341.8 | 2.1 | 34.0 | 14.5 | 44.6 | 7.8 | 10.1 | 212.0 | 27.8 | 31.6 | 2.3 | 13.8 | 20.2 |
| GB-S17-7 | 72.1 | 22.6 | 21.0 | 44.9 | 0.2 | 28.9 | 25.9 | 341.7 | 2.0 | 34.2 | 14.1 | 44.7 | 7.9 | 10.1 | 211.8 | 28.0 | 31.4 | 2.3 | 13.8 | 20.5 |
| GR-K1-1 | 71.9 | 22.9 | 21.2 | 44.1 | 0.2 | 29.3 | 26.2 | 341.6 | 2.1 | 34.0 | 14.5 | 44.4 | 7.7 | 10.0 | 211.5 | 28.1 | 31.7 | 2.3 | 13.7 | 20.6 |
| HU-AG-03 | 72.2 | 22.7 | 21.1 | 44.4 | 0.3 | 29.2 | 26.2 | 341.6 | 2.1 | 34.0 | 14.2 | 44.6 | 7.7 | 10.0 | 211.5 | 28.0 | 31.6 | 2.3 | 13.8 | 20.6 |
| HU-CR3-01 | 71.8 | 22.8 | 21.6 | 44.0 | 0.2 | 29.4 | 26.3 | 341.6 | 2.1 | 34.1 | 14.3 | 44.6 | 7.8 | 10.1 | 211.7 | 28.2 | 31.2 | 2.3 | 13.7 | 20.7 |
| HU-HO-2 | 72.0 | 22.9 | 21.3 | 44.2 | 0.2 | 29.3 | 26.1 | 341.6 | 2.1 | 34.1 | 14.3 | 44.5 | 7.7 | 10.0 | 211.4 | 28.2 | 31.7 | 2.3 | 13.7 | 20.8 |
| HU-K-6 | 72.5 | 22.8 | 21.2 | 44.3 | 0.2 | 29.3 | 26.0 | 341.6 | 2.1 | 34.1 | 14.4 | 44.6 | 7.8 | 10.0 | 211.9 | 27.8 | 31.3 | 2.3 | 13.7 | 20.5 |
| HU-KEL-01 | 72.3 | 22.7 | 21.3 | 44.4 | 0.3 | 29.1 | 26.3 | 341.6 | 2.1 | 34.0 | 14.3 | 44.5 | 7.8 | 10.0 | 211.8 | 28.0 | 31.4 | 2.3 | 13.7 | 20.4 |
| IE-DUB-5 | 72.3 | 22.5 | 20.9 | 44.5 | 0.2 | 29.1 | 26.0 | 341.8 | 2.0 | 34.1 | 14.2 | 44.7 | 7.8 | 10.0 | 212.1 | 27.8 | 31.8 | 2.3 | 13.8 | 20.3 |
| IE_DUB_1 | 72.2 | 22.5 | 20.9 | 44.5 | 0.2 | 29.1 | 26.0 | 341.8 | 2.0 | 34.1 | 14.2 | 44.7 | 7.8 | 10.0 | 212.3 | 27.8 | 31.7 | 2.3 | 13.8 | 20.3 |
| IL-BM-1 | 71.7 | 22.7 | 21.1 | 44.4 | 0.2 | 29.5 | 26.3 | 341.6 | 2.2 | 34.1 | 14.2 | 44.4 | 7.8 | 10.1 | 211.3 | 28.4 | 31.4 | 2.3 | 13.7 | 21.0 |
| IL-EY-17 | 71.6 | 22.7 | 21.1 | 44.3 | 0.2 | 29.4 | 26.4 | 341.6 | 2.1 | 34.1 | 14.3 | 44.6 | 7.7 | 10.4 | 211.0 | 28.6 | 31.4 | 2.3 | 13.5 | 20.9 |
| IL-G-3 | 71.5 | 22.6 | 21.3 | 44.2 | 0.2 | 29.4 | 26.2 | 341.6 | 2.1 | 34.2 | 14.2 | 44.5 | 7.7 | 10.4 | 211.4 | 28.2 | 31.8 | 2.3 | 13.6 | 20.9 |
| IL-M1-8 | 71.5 | 22.6 | 21.2 | 44.2 | 0.2 | 29.4 | 26.2 | 341.6 | 2.1 | 34.1 | 14.5 | 44.7 | 7.7 | 10.2 | 210.9 | 28.6 | 31.7 | 2.3 | 13.6 | 20.7 |
| IL-PS-2 | 71.8 | 22.7 | 21.0 | 44.8 | 0.2 | 29.1 | 26.1 | 341.5 | 2.1 | 34.1 | 14.0 | 44.4 | 7.7 | 10.1 | 211.3 | 28.4 | 32.1 | 2.3 | 13.6 | 20.9 |
| IL-PS-6 | 71.5 | 22.6 | 21.2 | 44.2 | 0.2 | 29.3 | 26.3 | 341.6 | 2.1 | 34.2 | 14.3 | 44.6 | 7.7 | 10.3 | 211.5 | 28.0 | 31.9 | 2.3 | 13.6 | 20.8 |
| IL-RAM-4 | 71.4 | 22.6 | 21.1 | 44.2 | 0.2 | 29.4 | 26.2 | 341.5 | 2.0 | 34.2 | 14.4 | 44.6 | 7.8 | 10.1 | 211.0 | 28.4 | 31.8 | 2.3 | 13.7 | 20.9 |
| IL-SH-4 | 71.4 | 22.6 | 21.0 | 44.3 | 0.2 | 29.5 | 26.4 | 341.6 | 2.1 | 34.0 | 14.2 | 44.5 | 7.7 | 10.3 | 210.9 | 28.7 | 31.8 | 2.3 | 13.5 | 21.0 |
| IL-TY-10 | 72.5 | 22.6 | 20.8 | 44.8 | 0.2 | 29.0 | 26.3 | 341.6 | 2.1 | 34.1 | 13.9 | 44.6 | 7.6 | 10.2 | 210.9 | 28.6 | 31.7 | 2.3 | 13.7 | 20.8 |
| IL-YERU | 71.4 | 22.7 | 21.3 | 44.1 | 0.2 | 29.6 | 26.4 | 341.6 | 2.1 | 34.1 | 14.2 | 44.5 | 7.7 | 10.1 | 211.2 | 28.2 | 31.8 | 2.3 | 13.7 | 21.0 |
| IR-GG1-1 | 72.0 | 22.8 | 21.1 | 44.3 | 0.2 | 29.2 | 26.2 | 341.6 | 2.1 | 34.1 | 14.4 | 44.4 | 7.7 | 10.0 | 211.4 | 28.4 | 31.9 | 2.3 | 13.7 | 20.6 |
| IR-GG1-102 | 72.4 | 22.6 | 21.1 | 44.5 | 0.2 | 29.2 | 26.3 | 341.4 | 2.0 | 34.2 | 14.1 | 44.4 | 7.7 | 10.1 | 211.6 | 28.2 | 31.5 | 2.3 | 13.6 | 20.7 |
| IT-ISR1-2 | 72.1 | 22.7 | 21.4 | 44.0 | 0.2 | 29.3 | 26.4 | 341.6 | 2.1 | 34.2 | 14.4 | 44.5 | 7.7 | 10.1 | 211.6 | 28.0 | 31.5 | 2.3 | 13.7 | 20.4 |
| IT-MDV-1 | 71.8 | 22.6 | 21.6 | 43.7 | 0.2 | 29.4 | 26.1 | 341.5 | 2.1 | 34.2 | 14.5 | 44.8 | 7.8 | 10.1 | 212.0 | 27.9 | 31.2 | 2.3 | 13.7 | 20.8 |
| IT-PER-2 | 71.8 | 22.7 | 21.0 | 44.8 | 0.2 | 29.0 | 26.1 | 341.5 | 2.0 | 34.3 | 14.0 | 44.5 | 7.8 | 10.1 | 211.5 | 28.3 | 31.8 | 2.3 | 13.8 | 20.8 |
| KG-SK-1 | 72.5 | 22.7 | 21.1 | 44.6 | 0.2 | 29.1 | 26.3 | 341.7 | 2.0 | 34.0 | 14.3 | 44.5 | 7.8 | 9.9 | 212.0 | 27.8 | 31.8 | 2.3 | 13.7 | 20.0 |
| MA-ES-3 | 71.6 | 22.6 | 21.4 | 43.9 | 0.2 | 29.4 | 25.9 | 341.5 | 2.1 | 34.1 | 14.4 | 44.8 | 7.8 | 10.0 | 212.4 | 27.8 | 31.5 | 2.3 | 13.8 | 20.7 |
| MN-DM-1 | 72.8 | 22.7 | 21.2 | 44.6 | 0.2 | 29.0 | 26.4 | 341.6 | 2.1 | 34.0 | 14.5 | 44.5 | 7.8 | 10.2 | 212.2 | 27.2 | 31.4 | 2.3 | 13.7 | 20.1 |
| NL_LA_1 | 71.6 | 22.6 | 21.4 | 44.0 | 0.2 | 29.4 | 26.2 | 341.6 | 2.0 | 34.2 | 14.4 | 44.6 | 7.8 | 10.0 | 212.0 | 28.2 | 31.6 | 2.3 | 13.7 | 20.4 |
| NO-AA-1 | 72.1 | 22.6 | 20.7 | 44.9 | 0.3 | 28.9 | 26.4 | 341.5 | 2.1 | 33.7 | 14.5 | 44.7 | 8.0 | 10.3 | 211.0 | 28.7 | 31.3 | 2.3 | 13.7 | 20.6 |
| NO-F-1 | 71.7 | 22.8 | 21.6 | 43.9 | 0.3 | 29.2 | 26.2 | 341.4 | 2.1 | 34.1 | 14.4 | 44.4 | 7.8 | 10.2 | 211.3 | 28.0 | 31.7 | 2.3 | 13.8 | 20.8 |
| NO-LADE-1 | 71.9 | 22.7 | 21.2 | 44.3 | 0.3 | 29.2 | 26.1 | 341.5 | 2.2 | 34.0 | 14.3 | 44.5 | 7.9 | 10.2 | 211.2 | 28.2 | 31.7 | 2.3 | 13.8 | 20.8 |
| NO-M3-1 | 71.7 | 22.6 | 21.3 | 44.4 | 0.2 | 29.0 | 26.4 | 341.8 | 2.1 | 33.9 | 14.5 | 44.5 | 8.0 | 10.1 | 210.9 | 28.4 | 31.8 | 2.3 | 13.7 | 20.7 |
| NO-MS-1 | 71.7 | 22.6 | 21.3 | 44.4 | 0.2 | 29.0 | 26.4 | 341.8 | 2.1 | 33.9 | 14.5 | 44.5 | 8.0 | 10.1 | 210.9 | 28.4 | 31.8 | 2.3 | 13.7 | 20.7 |
| NO-RO-1 | 72.2 | 22.7 | 20.9 | 44.7 | 0.2 | 29.0 | 26.3 | 341.6 | 2.1 | 34.1 | 14.4 | 44.6 | 8.0 | 10.0 | 210.9 | 28.6 | 31.5 | 2.3 | 13.7 | 20.6 |
| NO-V-7 | 72.4 | 22.8 | 20.9 | 44.6 | 0.2 | 29.1 | 26.1 | 341.5 | 2.1 | 34.2 | 14.4 | 44.6 | 8.0 | 10.1 | 210.8 | 28.7 | 31.3 | 2.3 | 13.7 | 20.5 |
| NO-VADSO-2 | 72.8 | 22.7 | 20.9 | 44.9 | 0.2 | 29.0 | 26.0 | 341.8 | 2.1 | 34.1 | 14.0 | 44.6 | 8.0 | 10.3 | 210.6 | 28.6 | 31.3 | 2.3 | 13.7 | 20.4 |
| NO-VAR1-5 | 72.4 | 22.7 | 21.0 | 44.9 | 0.2 | 28.9 | 26.2 | 341.7 | 2.1 | 34.1 | 14.2 | 44.4 | 8.0 | 10.1 | 210.8 | 28.7 | 31.3 | 2.3 | 13.8 | 20.6 |
| NO-VAR4-3 | 71.9 | 22.7 | 21.0 | 44.9 | 0.2 | 28.9 | 26.2 | 341.8 | 2.0 | 34.1 | 14.2 | 44.4 | 8.0 | 10.3 | 210.9 | 28.4 | 31.6 | 2.3 | 13.7 | 20.6 |
| NO-VAR5-6 | 72.1 | 22.7 | 21.0 | 45.0 | 0.2 | 28.8 | 26.2 | 341.7 | 2.1 | 34.1 | 14.2 | 44.4 | 8.0 | 10.4 | 210.8 | 28.7 | 31.4 | 2.3 | 13.8 | 20.5 |
| NO-VAR6-5 | 72.0 | 22.6 | 21.0 | 44.9 | 0.2 | 28.9 | 26.2 | 341.8 | 2.0 | 34.2 | 14.2 | 44.5 | 8.0 | 10.2 | 210.9 | 28.4 | 31.5 | 2.3 | 13.7 | 20.6 |
| PL-KNP-4 | 72.7 | 22.6 | 20.9 | 44.5 | 0.2 | 29.1 | 26.3 | 341.8 | 2.2 | 33.9 | 14.2 | 44.4 | 7.8 | 10.0 | 211.3 | 28.1 | 31.3 | 2.3 | 13.9 | 20.7 |
| PL-W1-1 | 72.1 | 22.7 | 21.0 | 44.3 | 0.2 | 29.4 | 26.2 | 341.8 | 2.1 | 34.2 | 14.1 | 44.6 | 7.8 | 10.0 | 211.7 | 28.0 | 31.1 | 2.3 | 13.9 | 20.8 |
| PL-W2-1 | 72.4 | 22.8 | 21.0 | 44.5 | 0.2 | 29.1 | 26.2 | 341.7 | 2.2 | 34.4 | 14.2 | 44.5 | 7.8 | 10.1 | 210.9 | 28.3 | 31.3 | 2.3 | 13.7 | 20.5 |
| RO-DS-3 | 71.4 | 22.8 | 21.5 | 44.2 | 0.3 | 29.4 | 26.4 | 341.6 | 2.1 | 34.2 | 14.0 | 44.5 | 7.7 | 10.0 | 211.4 | 28.1 | 31.5 | 2.3 | 13.7 | 21.0 |
| RU-AL1-11 | 72.5 | 22.7 | 21.1 | 44.6 | 0.2 | 28.7 | 26.2 | 341.6 | 2.1 | 33.8 | 14.6 | 44.6 | 7.8 | 10.3 | 211.0 | 28.3 | 31.7 | 2.3 | 13.6 | 20.4 |
| RU-AS2-1 | 72.1 | 22.6 | 21.0 | 44.7 | 0.2 | 29.0 | 26.3 | 341.6 | 2.0 | 34.1 | 13.9 | 44.7 | 7.7 | 10.0 | 211.2 | 28.4 | 31.7 | 2.3 | 13.7 | 20.8 |
| RU-AST1-1 | 71.6 | 22.6 | 21.4 | 44.3 | 0.2 | 29.3 | 26.2 | 341.6 | 2.0 | 34.4 | 14.2 | 44.5 | 7.7 | 10.0 | 211.1 | 28.3 | 31.7 | 2.3 | 13.7 | 20.8 |
| RU-B3-01 | 72.2 | 22.7 | 20.9 | 44.8 | 0.2 | 29.1 | 26.2 | 341.7 | 2.1 | 34.3 | 13.9 | 44.4 | 7.7 | 10.1 | 210.5 | 28.7 | 31.8 | 2.3 | 13.7 | 20.9 |
| RU-B5-1 | 72.7 | 22.6 | 20.9 | 44.7 | 0.2 | 29.0 | 26.3 | 341.5 | 2.0 | 33.9 | 14.1 | 44.4 | 7.7 | 10.0 | 211.5 | 28.4 | 31.2 | 2.3 | 13.8 | 20.8 |
| RU-BAL1-2 | 72.8 | 22.7 | 20.7 | 45.1 | 0.2 | 28.9 | 26.3 | 341.4 | 2.1 | 34.0 | 14.3 | 44.4 | 7.7 | 10.2 | 211.6 | 27.9 | 31.6 | 2.3 | 13.6 | 20.5 |
| RU-BAYA1-1 | 72.4 | 22.6 | 21.0 | 44.9 | 0.2 | 29.0 | 26.2 | 341.5 | 2.1 | 34.1 | 14.5 | 44.3 | 7.8 | 10.3 | 211.3 | 27.9 | 31.8 | 2.3 | 13.6 | 20.4 |
| RU-BN-BB6 | 72.9 | 22.6 | 21.0 | 44.5 | 0.2 | 29.1 | 26.2 | 341.5 | 2.1 | 34.1 | 14.3 | 44.6 | 7.7 | 10.1 | 211.1 | 28.6 | 31.0 | 2.3 | 13.7 | 20.6 |
| RU-BOL1-1 | 72.3 | 22.6 | 20.8 | 44.8 | 0.2 | 29.0 | 26.2 | 341.6 | 2.1 | 34.1 | 14.3 | 44.7 | 7.7 | 10.1 | 211.0 | 28.3 | 31.4 | 2.3 | 13.7 | 20.9 |
| RU-BOR-1 | 72.2 | 22.7 | 21.0 | 44.7 | 0.2 | 29.1 | 26.2 | 341.6 | 2.1 | 34.0 | 14.5 | 44.5 | 7.7 | 10.0 | 211.8 | 28.0 | 31.4 | 2.3 | 13.8 | 20.5 |
| RU-BP-1 | 71.2 | 22.7 | 21.2 | 44.2 | 0.2 | 29.6 | 26.2 | 341.6 | 2.0 | 34.2 | 14.2 | 44.5 | 7.7 | 10.1 | 211.3 | 28.3 | 31.8 | 2.3 | 13.7 | 21.1 |
| RU-BU2-2 | 72.6 | 22.7 | 21.2 | 44.4 | 0.2 | 29.1 | 26.5 | 341.5 | 2.1 | 34.1 | 14.5 | 44.6 | 7.8 | 10.3 | 211.0 | 27.8 | 31.8 | 2.3 | 13.7 | 20.0 |
| RU-C10-06 | 73.0 | 22.7 | 20.8 | 45.0 | 0.2 | 28.8 | 26.3 | 341.7 | 2.1 | 34.0 | 14.3 | 44.5 | 7.7 | 10.2 | 210.6 | 28.5 | 31.2 | 2.3 | 13.7 | 20.5 |
| RU-C12-1 | 72.5 | 22.8 | 21.0 | 44.6 | 0.2 | 29.1 | 26.3 | 341.6 | 2.0 | 34.1 | 14.3 | 44.3 | 7.8 | 10.1 | 211.2 | 28.4 | 31.3 | 2.3 | 13.8 | 20.6 |
| RU-C20-1 | 73.0 | 22.6 | 20.8 | 44.8 | 0.2 | 29.0 | 26.3 | 341.5 | 2.0 | 34.1 | 14.3 | 44.6 | 7.7 | 10.1 | 210.8 | 28.5 | 31.2 | 2.3 | 13.7 | 20.5 |
| RU-FR-LR8-1 | 71.4 | 23.0 | 21.5 | 44.2 | 0.3 | 29.2 | 25.9 | 341.5 | 2.1 | 34.2 | 14.5 | 44.7 | 7.7 | 10.1 | 211.6 | 28.1 | 31.7 | 2.3 | 13.8 | 20.7 |
| RU-HA2-12 | 72.6 | 22.7 | 20.9 | 44.8 | 0.2 | 29.0 | 26.3 | 341.7 | 2.2 | 34.5 | 14.5 | 44.4 | 7.8 | 10.2 | 210.3 | 28.6 | 31.4 | 2.3 | 13.7 | 20.1 |
| RU-HAI-1 | 73.0 | 22.6 | 20.8 | 45.0 | 0.2 | 29.0 | 26.3 | 341.5 | 2.1 | 33.9 | 14.7 | 44.5 | 7.5 | 10.3 | 211.4 | 27.8 | 31.4 | 2.3 | 13.7 | 20.2 |
| RU-IRK1-12 | 72.7 | 22.8 | 20.7 | 45.1 | 0.2 | 28.8 | 26.3 | 341.4 | 2.1 | 34.0 | 14.4 | 44.4 | 7.9 | 10.3 | 211.2 | 27.8 | 31.6 | 2.3 | 13.7 | 20.5 |
| RU-IRK2-14 | 72.6 | 22.8 | 20.9 | 45.1 | 0.2 | 28.8 | 26.4 | 341.5 | 2.1 | 33.9 | 14.2 | 44.4 | 7.6 | 10.2 | 210.8 | 28.4 | 32.1 | 2.3 | 13.6 | 20.4 |
| RU-IRK3-1 | 73.1 | 22.7 | 20.9 | 44.9 | 0.2 | 28.8 | 26.1 | 341.4 | 2.2 | 34.0 | 14.7 | 44.5 | 7.9 | 10.2 | 211.3 | 28.0 | 30.7 | 2.3 | 13.7 | 20.6 |
| RU-IRK4-10 | 72.4 | 22.7 | 20.7 | 45.2 | 0.2 | 28.7 | 26.4 | 341.5 | 2.1 | 34.2 | 14.3 | 44.4 | 7.8 | 10.2 | 210.7 | 28.3 | 32.0 | 2.3 | 13.7 | 20.3 |
| RU-KOR1-1 | 72.3 | 22.6 | 21.1 | 44.5 | 0.2 | 29.0 | 26.2 | 341.6 | 2.1 | 34.2 | 14.2 | 44.6 | 8.0 | 10.1 | 211.2 | 28.3 | 31.3 | 2.3 | 13.8 | 20.7 |
| RU-KUI-2 | 72.4 | 22.7 | 21.2 | 44.3 | 0.2 | 29.2 | 26.2 | 341.7 | 2.0 | 34.1 | 14.4 | 44.3 | 7.7 | 10.1 | 210.9 | 28.6 | 31.5 | 2.3 | 13.8 | 20.6 |
| RU-MA10-3 | 72.5 | 22.7 | 20.7 | 44.7 | 0.2 | 29.2 | 26.2 | 341.8 | 2.1 | 34.2 | 14.1 | 44.6 | 7.6 | 10.0 | 210.7 | 28.3 | 31.7 | 2.3 | 13.7 | 20.8 |
| RU-MA13-3 | 72.1 | 22.7 | 20.8 | 44.5 | 0.2 | 29.2 | 26.3 | 341.6 | 2.1 | 34.1 | 14.0 | 44.4 | 7.7 | 10.1 | 211.1 | 28.4 | 31.9 | 2.3 | 13.8 | 21.0 |
| RU-MA20-10 | 72.1 | 22.6 | 20.9 | 44.6 | 0.2 | 29.0 | 26.2 | 341.7 | 2.0 | 34.4 | 14.2 | 44.7 | 7.7 | 10.1 | 210.7 | 28.5 | 31.9 | 2.3 | 13.8 | 20.6 |
| RU-MA23-8 | 72.8 | 22.7 | 20.8 | 44.8 | 0.2 | 29.0 | 26.3 | 341.6 | 2.0 | 34.2 | 13.8 | 44.6 | 7.7 | 10.1 | 210.4 | 28.6 | 31.7 | 2.3 | 13.7 | 20.9 |
| RU-MA3-5 | 72.6 | 22.8 | 20.8 | 44.8 | 0.2 | 29.0 | 26.3 | 341.5 | 2.1 | 34.0 | 14.2 | 44.4 | 7.7 | 10.0 | 211.1 | 28.5 | 31.5 | 2.3 | 13.7 | 20.7 |
| RU-MA4-3 | 72.8 | 22.6 | 20.9 | 44.9 | 0.2 | 28.9 | 26.2 | 341.5 | 2.0 | 34.0 | 14.3 | 44.5 | 7.6 | 10.1 | 211.0 | 28.1 | 31.6 | 2.3 | 13.7 | 20.9 |
| RU-MA8-01 | 72.3 | 22.7 | 20.6 | 44.8 | 0.2 | 29.0 | 26.3 | 341.6 | 2.1 | 34.1 | 14.0 | 44.3 | 7.8 | 10.0 | 211.4 | 28.4 | 31.6 | 2.3 | 13.8 | 20.9 |
| RU-NOV-3-2 | 71.7 | 22.8 | 21.4 | 44.4 | 0.2 | 29.0 | 26.2 | 341.6 | 2.0 | 34.4 | 14.1 | 44.4 | 8.0 | 10.0 | 211.3 | 28.6 | 31.4 | 2.3 | 13.8 | 20.8 |
| RU-NOV-4-7 | 72.1 | 22.9 | 21.7 | 44.1 | 0.2 | 28.8 | 26.1 | 341.6 | 2.0 | 34.6 | 13.9 | 44.4 | 7.9 | 10.0 | 210.6 | 28.5 | 31.8 | 2.3 | 13.9 | 20.8 |
| RU-NOV1-04 | 71.7 | 22.7 | 21.5 | 44.1 | 0.2 | 29.2 | 26.2 | 341.4 | 2.1 | 34.1 | 14.4 | 44.4 | 8.0 | 10.0 | 211.3 | 28.3 | 31.5 | 2.3 | 13.7 | 20.9 |
| RU-NOV2-01 | 72.2 | 22.7 | 21.3 | 44.5 | 0.2 | 29.0 | 26.3 | 341.4 | 2.1 | 34.2 | 14.2 | 44.4 | 8.0 | 10.1 | 211.0 | 28.4 | 31.6 | 2.3 | 13.7 | 20.7 |
| RU-NOV5-17 | 72.5 | 22.8 | 21.4 | 44.5 | 0.2 | 28.8 | 26.1 | 341.6 | 2.0 | 34.4 | 14.3 | 44.4 | 7.8 | 10.0 | 210.8 | 28.1 | 31.6 | 2.3 | 13.8 | 20.7 |
| RU-P1-01 | 72.1 | 22.8 | 21.0 | 44.3 | 0.2 | 29.5 | 26.2 | 341.6 | 2.1 | 34.0 | 14.3 | 44.5 | 7.7 | 10.2 | 211.4 | 28.1 | 31.4 | 2.3 | 13.8 | 20.6 |
| RU-PIK-2 | 71.6 | 22.6 | 21.2 | 44.3 | 0.2 | 29.4 | 26.2 | 341.5 | 2.0 | 34.1 | 14.4 | 44.5 | 7.7 | 10.4 | 211.5 | 27.8 | 31.6 | 2.3 | 13.7 | 21.1 |
| RU-R2-1 | 72.4 | 22.7 | 21.2 | 44.5 | 0.2 | 28.9 | 26.2 | 341.6 | 2.1 | 34.3 | 14.1 | 44.6 | 8.0 | 10.1 | 210.9 | 28.3 | 31.4 | 2.3 | 13.7 | 20.7 |
| RU-RM1-2 | 72.0 | 22.7 | 20.9 | 44.7 | 0.2 | 29.1 | 26.2 | 341.6 | 2.1 | 34.0 | 14.3 | 44.6 | 7.7 | 10.3 | 210.9 | 28.7 | 31.5 | 2.3 | 13.8 | 20.6 |
| RU-RO1-3 | 72.1 | 22.7 | 20.8 | 44.8 | 0.2 | 29.0 | 26.2 | 341.6 | 2.1 | 34.0 | 14.4 | 44.4 | 7.8 | 10.1 | 211.3 | 28.5 | 31.5 | 2.3 | 13.7 | 20.6 |
| RU-RT1-135 | 72.4 | 22.6 | 20.8 | 44.6 | 0.2 | 29.1 | 26.3 | 341.7 | 2.1 | 34.1 | 14.3 | 44.6 | 7.6 | 10.0 | 211.2 | 28.3 | 31.6 | 2.3 | 13.7 | 20.5 |
| RU-RT21-1 | 72.5 | 22.6 | 20.8 | 44.6 | 0.2 | 29.1 | 26.3 | 341.7 | 2.1 | 34.2 | 14.1 | 44.8 | 7.6 | 10.0 | 211.0 | 28.2 | 31.7 | 2.3 | 13.7 | 20.6 |
| RU-SAM2-1 | 72.0 | 22.8 | 21.1 | 44.5 | 0.2 | 29.1 | 26.2 | 341.7 | 2.0 | 34.1 | 14.2 | 44.4 | 7.7 | 10.0 | 211.4 | 28.2 | 31.7 | 2.3 | 13.8 | 20.6 |
| RU-SAM5-1 | 72.5 | 22.7 | 21.0 | 44.6 | 0.2 | 29.1 | 26.3 | 341.6 | 2.1 | 34.0 | 14.1 | 44.4 | 7.7 | 10.1 | 210.8 | 28.5 | 31.9 | 2.3 | 13.7 | 20.6 |
| RU-SPB-35 | 71.0 | 22.8 | 21.3 | 44.3 | 0.2 | 29.2 | 26.1 | 341.4 | 2.1 | 34.4 | 14.3 | 44.4 | 7.7 | 10.2 | 211.7 | 28.2 | 31.8 | 2.3 | 13.7 | 21.0 |
| RU-SYR1-1 | 72.7 | 22.7 | 20.8 | 44.7 | 0.2 | 29.2 | 26.2 | 341.6 | 2.2 | 34.0 | 14.3 | 44.6 | 7.7 | 10.3 | 210.5 | 28.4 | 31.4 | 2.3 | 13.7 | 20.7 |
| RU-TU1-1 | 72.0 | 22.7 | 21.2 | 44.6 | 0.2 | 29.1 | 26.2 | 341.5 | 2.1 | 34.2 | 14.2 | 44.5 | 7.9 | 10.2 | 211.1 | 28.3 | 31.6 | 2.3 | 13.7 | 20.7 |
| RU-TY1-3 | 72.7 | 22.8 | 21.1 | 44.7 | 0.2 | 28.9 | 26.3 | 341.7 | 2.3 | 34.3 | 14.4 | 44.3 | 7.8 | 10.4 | 210.0 | 28.4 | 32.0 | 2.3 | 13.6 | 20.2 |
| RU-TY2-2 | 73.0 | 22.8 | 21.5 | 44.3 | 0.2 | 29.1 | 26.2 | 341.5 | 2.1 | 34.4 | 14.4 | 44.3 | 7.8 | 10.3 | 210.8 | 28.2 | 31.3 | 2.3 | 13.7 | 20.3 |
| RU-TY4-1 | 72.4 | 22.6 | 21.4 | 44.0 | 0.2 | 29.2 | 26.2 | 341.5 | 2.1 | 34.1 | 14.5 | 44.5 | 7.9 | 10.2 | 213.0 | 26.9 | 31.3 | 2.3 | 13.8 | 20.1 |
| RU-TY5-1 | 71.9 | 22.7 | 21.3 | 44.2 | 0.2 | 29.4 | 26.4 | 341.5 | 2.1 | 34.1 | 14.3 | 44.4 | 7.8 | 10.2 | 211.7 | 28.0 | 31.8 | 2.3 | 13.7 | 20.2 |
| RU-VOL-36 | 72.2 | 22.7 | 21.1 | 44.7 | 0.2 | 28.9 | 26.3 | 341.7 | 2.0 | 34.3 | 14.2 | 44.5 | 7.8 | 10.0 | 210.6 | 28.3 | 31.8 | 2.3 | 13.7 | 20.8 |
| RU-YAK1-6 | 72.7 | 22.7 | 20.6 | 45.3 | 0.2 | 28.7 | 26.3 | 341.4 | 2.2 | 34.2 | 14.5 | 44.3 | 7.7 | 10.2 | 210.8 | 28.3 | 31.9 | 2.3 | 13.7 | 20.4 |
| RU-YAK3-25 | 72.5 | 22.7 | 20.8 | 45.0 | 0.2 | 28.9 | 26.4 | 341.4 | 2.1 | 34.2 | 14.5 | 44.3 | 7.7 | 10.1 | 211.2 | 27.8 | 31.9 | 2.3 | 13.6 | 20.5 |
| RU-ZB1-1 | 73.3 | 22.7 | 21.0 | 44.6 | 0.2 | 29.1 | 26.3 | 341.7 | 2.1 | 34.0 | 14.6 | 44.6 | 7.8 | 10.3 | 210.4 | 28.0 | 31.7 | 2.3 | 13.7 | 19.9 |
| RU-ZB2-22 | 72.0 | 22.7 | 21.1 | 44.3 | 0.2 | 29.2 | 26.3 | 341.6 | 2.1 | 34.2 | 14.6 | 44.6 | 8.0 | 10.3 | 210.8 | 28.1 | 32.0 | 2.3 | 13.7 | 20.2 |
| RU-ZB4-12 | 72.0 | 22.9 | 21.5 | 44.1 | 0.2 | 29.1 | 26.2 | 341.7 | 2.1 | 34.1 | 14.5 | 44.6 | 7.8 | 10.3 | 211.4 | 27.9 | 31.7 | 2.3 | 13.7 | 20.4 |
| RU-ZB5-1 | 72.2 | 22.7 | 21.4 | 44.2 | 0.2 | 29.2 | 26.2 | 341.6 | 2.2 | 34.0 | 14.7 | 44.5 | 7.8 | 10.3 | 211.0 | 28.0 | 31.8 | 2.3 | 13.7 | 20.3 |
| RU-ZB6-03 | 72.6 | 22.8 | 21.3 | 44.5 | 0.2 | 28.9 | 26.3 | 341.6 | 2.1 | 34.2 | 14.6 | 44.5 | 7.8 | 10.2 | 210.9 | 28.0 | 31.7 | 2.3 | 13.7 | 20.2 |
| RU_IRK1_10 | 72.5 | 22.7 | 20.7 | 45.2 | 0.2 | 28.7 | 26.4 | 341.5 | 2.1 | 34.2 | 14.3 | 44.4 | 7.8 | 10.2 | 210.7 | 28.3 | 32.0 | 2.3 | 13.7 | 20.3 |
| RU_IRK5_72 | 72.7 | 22.7 | 21.0 | 45.1 | 0.2 | 28.7 | 26.3 | 341.6 | 2.1 | 34.1 | 14.5 | 44.3 | 7.8 | 10.2 | 210.4 | 28.3 | 32.0 | 2.3 | 13.6 | 20.2 |
| RU_IRK9_1 | 72.8 | 22.7 | 21.4 | 44.2 | 0.2 | 29.2 | 26.5 | 341.6 | 2.2 | 34.0 | 14.4 | 44.3 | 7.8 | 10.3 | 211.8 | 27.4 | 31.2 | 2.3 | 13.6 | 20.2 |
| RU_KA2_6 | 72.4 | 22.8 | 20.8 | 44.7 | 0.2 | 29.1 | 26.2 | 341.6 | 2.0 | 34.1 | 14.1 | 44.6 | 7.6 | 10.0 | 211.1 | 28.4 | 31.7 | 2.3 | 13.7 | 20.8 |
| RU_KA3_2 | 72.5 | 22.6 | 20.8 | 44.6 | 0.2 | 29.2 | 26.3 | 341.5 | 2.1 | 34.2 | 14.3 | 44.6 | 7.6 | 10.0 | 211.1 | 28.3 | 31.7 | 2.3 | 13.7 | 20.6 |
| RU_KAI_105 | 72.6 | 22.7 | 21.0 | 44.5 | 0.2 | 29.2 | 26.3 | 341.6 | 2.1 | 33.9 | 14.3 | 44.4 | 7.7 | 10.1 | 211.1 | 28.2 | 31.5 | 2.3 | 13.7 | 20.7 |
| RU_MA21_2 | 72.5 | 22.6 | 20.9 | 44.7 | 0.2 | 28.9 | 26.3 | 341.6 | 2.1 | 34.1 | 14.0 | 44.5 | 7.7 | 10.1 | 211.2 | 28.3 | 31.6 | 2.3 | 13.7 | 20.8 |
| SE-BY-6 | 72.0 | 22.5 | 21.6 | 43.8 | 0.3 | 29.5 | 26.2 | 341.5 | 2.0 | 34.2 | 14.5 | 44.7 | 7.7 | 9.9 | 211.5 | 28.2 | 31.4 | 2.3 | 13.9 | 20.4 |
| SE-G1-9c | 70.8 | 22.5 | 21.4 | 44.3 | 0.2 | 29.4 | 26.3 | 341.7 | 2.1 | 34.0 | 14.6 | 44.5 | 7.8 | 10.2 | 211.7 | 28.2 | 31.7 | 2.3 | 13.7 | 20.7 |
| SE-G2-8c | 71.0 | 22.7 | 21.4 | 44.4 | 0.2 | 29.2 | 26.2 | 341.6 | 2.1 | 34.2 | 14.5 | 44.4 | 7.8 | 10.2 | 211.3 | 28.3 | 31.5 | 2.3 | 13.7 | 20.8 |
| SE-G4-20 | 71.0 | 22.8 | 21.5 | 44.4 | 0.2 | 29.3 | 26.2 | 341.6 | 2.1 | 34.2 | 14.6 | 44.4 | 7.8 | 10.1 | 211.2 | 28.3 | 31.6 | 2.3 | 13.7 | 20.8 |
| SE-GN1-3d3 | 71.1 | 22.6 | 21.2 | 44.5 | 0.2 | 29.3 | 26.0 | 341.6 | 2.1 | 34.0 | 14.5 | 44.5 | 7.8 | 10.2 | 212.0 | 28.1 | 31.5 | 2.3 | 13.7 | 20.8 |
| SE-GN2-3A10 | 71.0 | 22.6 | 21.2 | 44.5 | 0.2 | 29.3 | 26.1 | 341.6 | 2.1 | 34.0 | 14.5 | 44.5 | 7.8 | 10.2 | 211.8 | 28.1 | 31.6 | 2.3 | 13.7 | 20.9 |
| SE-H1-4 | 72.5 | 22.7 | 20.8 | 44.9 | 0.2 | 29.0 | 26.3 | 341.6 | 2.1 | 34.0 | 14.4 | 44.5 | 7.8 | 10.3 | 210.8 | 28.5 | 31.3 | 2.3 | 13.8 | 20.5 |
| TN-RA-2 | 72.1 | 22.4 | 21.2 | 44.0 | 0.2 | 29.4 | 26.0 | 341.4 | 2.2 | 34.3 | 14.7 | 44.7 | 7.8 | 10.0 | 212.3 | 28.0 | 31.1 | 2.3 | 13.6 | 20.5 |
| TR-EG-1 | 72.7 | 22.6 | 20.9 | 44.7 | 0.2 | 29.2 | 26.2 | 341.5 | 2.1 | 33.9 | 14.3 | 44.5 | 7.7 | 10.1 | 211.4 | 28.2 | 31.5 | 2.3 | 13.7 | 20.5 |
| UA-KRI-7 | 72.4 | 22.5 | 20.8 | 44.8 | 0.2 | 29.1 | 26.4 | 341.6 | 2.0 | 34.1 | 13.9 | 44.4 | 7.7 | 10.1 | 211.4 | 28.2 | 31.8 | 2.3 | 13.8 | 20.8 |
| US-D-1 | 73.4 | 22.9 | 20.8 | 45.0 | 0.2 | 28.8 | 26.5 | 341.2 | 2.2 | 33.8 | 14.8 | 44.1 | 8.0 | 10.3 | 209.5 | 28.8 | 31.6 | 2.3 | 13.7 | 20.1 |
| US-FAR1-1 | 73.3 | 22.9 | 20.9 | 45.0 | 0.2 | 28.8 | 26.5 | 341.2 | 2.2 | 33.8 | 14.5 | 44.1 | 8.0 | 10.4 | 210.2 | 28.7 | 31.3 | 2.3 | 13.7 | 20.2 |
| US-SP15-1 | 73.0 | 22.9 | 20.7 | 45.3 | 0.2 | 28.8 | 26.6 | 341.2 | 2.3 | 34.0 | 14.6 | 44.2 | 8.0 | 10.4 | 209.2 | 28.8 | 31.8 | 2.3 | 13.7 | 20.2 |
| US-SP163-1 | 73.0 | 22.9 | 20.7 | 45.3 | 0.2 | 28.8 | 26.6 | 341.2 | 2.3 | 34.0 | 14.6 | 44.2 | 8.0 | 10.4 | 209.2 | 28.9 | 31.8 | 2.3 | 13.7 | 20.2 |
| US-SP2-1 | 73.0 | 22.9 | 20.7 | 45.3 | 0.2 | 28.8 | 26.6 | 341.2 | 2.3 | 34.0 | 14.6 | 44.2 | 8.0 | 10.4 | 209.2 | 28.8 | 31.8 | 2.3 | 13.7 | 20.2 |
| US-SP22-1-1 | 73.0 | 22.9 | 20.7 | 45.3 | 0.2 | 28.8 | 26.6 | 341.2 | 2.3 | 34.0 | 14.6 | 44.2 | 8.0 | 10.4 | 209.2 | 28.9 | 31.8 | 2.3 | 13.7 | 20.2 |
| US-SP6-10 | 73.0 | 22.9 | 20.7 | 45.3 | 0.2 | 28.8 | 26.6 | 341.2 | 2.3 | 34.0 | 14.6 | 44.2 | 8.0 | 10.4 | 209.2 | 28.8 | 31.8 | 2.3 | 13.7 | 20.2 |
| US-SP7-1 | 73.0 | 22.9 | 20.7 | 45.3 | 0.2 | 28.8 | 26.6 | 341.2 | 2.3 | 34.0 | 14.6 | 44.2 | 8.0 | 10.4 | 209.2 | 28.8 | 31.8 | 2.3 | 13.7 | 20.2 |
| US-SU-1 | 73.6 | 22.9 | 20.9 | 45.0 | 0.2 | 28.8 | 26.5 | 341.2 | 2.2 | 33.8 | 14.6 | 44.1 | 8.0 | 10.4 | 210.2 | 28.5 | 31.1 | 2.3 | 13.7 | 20.2 |

### **Table S4.** Mean amino acid content (counts per 1000 residues) in collagen genes averaged among the D. magna genotypes used in this study. Mean, coefficient of variation (CV) and standard deviation (STD) are shown.

| **Amino acid** | **Mean** | **CV** | **STD** |
| --- | --- | --- | --- |
| Alanine | 72.21 | 0.86 | 0.62 |
| Arginine | 22.69 | 0.56 | 0.13 |
| Asparagine | 21.07 | 1.48 | 0.31 |
| Aspartic acid | 44.53 | 0.91 | 0.41 |
| Cysteine | 0.18 | 26.45 | 0.05 |
| Glutamic acid | 29.12 | 0.84 | 0.25 |
| Glutamine | 26.24 | 0.65 | 0.17 |
| Glycine | 341.56 | 0.04 | 0.15 |
| Histidine | 2.12 | 3.85 | 0.08 |
| Isoleucine | 34.07 | 0.52 | 0.18 |
| Leucine | 14.35 | 1.50 | 0.22 |
| Lysine | 44.50 | 0.38 | 0.17 |
| Methionine | 7.79 | 1.67 | 0.13 |
| Phenylalanine | 10.15 | 1.39 | 0.14 |
| Proline | 211.26 | 0.35 | 0.74 |
| Serine | 28.19 | 1.37 | 0.39 |
| Threonine | 31.54 | 0.92 | 0.29 |
| Tryptophan | 2.31 | 1.45 | 0.03 |
| Tyrosine | 13.72 | 0.52 | 0.07 |
| Valine | 20.56 | 1.37 | 0.28 |

### **Table S5.** Description of proline and serine amino acids content across the 50 collagen genes of D. magna, and its correlation with annual mean temperature. Summary statistics showing the sample size (N), mean (Mean), standard deviation (STD), maximum (Max) and minimum (Min) values. For the entire dataset (All), a multiple regression model was performed using annual mean temperature (Temp.) and phylogenetic lineage as independent variables. Additionally, a linear regression model was conducted for each lineage independently: Western Eurasia (WA), East Asia (EA) and North America (NA). The degrees of freedom (Df), the F-value, p-value and percentage of explained variance (Exp. Var; in %) are shown for each regression model. Significant p-values of a threshold at 0.05 are in bold

| **A. Proline** | | | | | | | | | | | |
| --- | --- | --- | --- | --- | --- | --- | --- | --- | --- | --- | --- |
| **Data** | **N** | **Mean** | **STD** | **Max** | **Min** | **Regression model** | | | | | |
|  |  |  |  |  |  | **Variables** | **Df** | **F-value** | ***p*-value** | **slope** | **Exp. Var.** |
| All | 215 | 214.98 | 0.66 | 216.58 | 213.17 | Temp. | 1 | 141.09 | **<2.2x10^-16^** | 0.08 | 21.96 |
|  |  |  |  |  |  | Lineage | 2 | 137.75 | **<2.2x10^-16^** | - | 42.88 |
|  |  |  |  |  |  | Interaction | 2 | 8.45 | **2.9x10^-4^** | - | 2.63 |
| WA | 161 | 215.13 | 0.42 | 216.28 | 214.27 | Temp. | 1 | 67.32 | **7.4x10^-14^** | 0.04 | 29.75 |
| EA | 35 | 215.08 | 0.69 | 216.58 | 214.20 | Temp. | 1 | 22.19 | **4.32x10^-5^** | 0.08 | 40.20 |
| NA | 19 | 213.46 | 0.23 | 214.10 | 213.17 | Temp. | 1 | 1.72 | 0.21 | 9.1x10^-3^ | 9.19 |

| **B. Serine** | | | | | | | | | | | |
| --- | --- | --- | --- | --- | --- | --- | --- | --- | --- | --- | --- |
| **Data** | **N** | **Mean** | **STD** | **Max** | **Min** | **Regression model** | | | | | |
|  |  |  |  |  |  | **Variables** | **Df** | **F-value** | ***p*-value** | **slope** | **Exp. Var.** |
| All | 215 | 29.83 | 0.34 | 30.80 | 28.69 | Temp. | 1 | 35.82 | **9.3x10^-9^** | -0.04 | 7.85 |
|  |  |  |  |  |  | Lineage | 2 | 101.73 | **<2.2x10^-16^** | - | 44.61 |
|  |  |  |  |  |  | Interaction | 2 | 3.89 | **0.02** | - | 1.70 |
| WA | 161 | 29.80 | 0.23 | 30.29 | 29.13 | Temp. | 1 | 28.71 | **2.9x10^-7^** | -0.02 | 15.29 |
| EA | 35 | 29.61 | 0.42 | 30.25 | 28.69 | Temp. | 1 | 11.58 | **1.8x10^-3^** | -0.04 | 25.98 |
| NA | 19 | 30.51 | 0.14 | 30.80 | 30.19 | Temp. | 1 | 11.56 | **3.4x10^-3^** | -0.01 | 40.47 |

### **Table S6.** Phylogenetic generalized least squares (PGLS) model to test the correlation of proline (A) and serine (B) content with the annual mean temperature by correcting for the phylogenetic signal. Significant correlations are highlighted in bold (i.e., p-value ≤ 0.05 on the slope). The degrees of freedom (Df), the F-value and p-value and are shown for each regression model. Significant p-values of a threshold at 0.05 are in bold.

| A. Model: Proline ~ Annual Mean Temperature * Lineage | | | | |
| --- | --- | --- | --- | --- |
| Independent variables | **Df** | **Mean Squares** | **F-value** | ***p-*Value** |
| Annual Mean Temperature | 1 | 26.12 | 7.68 | **6.1x10^-3^** |
| Lineage | 2 | 15.92 | 4.68 | **0.01** |
| Interaction | 2 | 15.25 | 4.48 | **0.01** |

| B. Model: Serine ~ Annual Mean Temperature * Lineage | | | | |
| --- | --- | --- | --- | --- |
| Independent variables | **Df** | **Mean Squares** | **F-value** | ***p-*Value** |
| Annual Mean Temperature | 1 | 3.76 | 2.92 | 0.09 |
| Lineage | 2 | 8.22 | 6.37 | **2.1 x10^-3^** |
| Interaction | 2 | 2.89 | 2.24 | 0.11 |

### **Table S7.** πN/πS ratio estimates for the Western Eurasia lineage, excluding the genotypes originated from rockpools (N= 130). Mean values are reported together with standard deviation. Wilcoxon rank tests show that πN/πS ratio of BUSCO genes is significantly smaller than those of collagen (W=23287, p-value=8.9x10^-9^) and immune genes (W=175198, p-value<2.2x10^-16^). Immune and collagen genes show a statistically similar πN/πS ratio (W=5561, p-value=0.96).

| **Gene set** | **πN/πS ratio** |
| --- | --- |
| Collagen genes | 0.27±0.18 |
| Immune genes | 0.29±0.28 |
| BUSCO genes | 0.14±0.21 |

### **Table S8.** Description of the immune and BUSCO gene sets used in this study. The coordinates based on the D. magna reference genome are displayed with the chromosome (Chr), contig number, and start and end position (Pos).

| **Gene Id** | **Chr** | **Contig** | **Start Pos** | **End Pos** | **Gene set** |
| --- | --- | --- | --- | --- | --- |
| maker-000016F-augustus-gene-4.85-mRNA-1 | 1 | 000016F | 395150 | 395829 | Immune |
| maker-000016F-augustus-gene-7.48-mRNA-1 | 1 | 000016F | 687816 | 693338 | Immune |
| maker-000016F-augustus-gene-8.70-mRNA-1 | 1 | 000016F | 777221 | 778965 | Immune |
| genemark-000016F-processed-gene-8.99-mRNA-1 | 1 | 000016F | 879818 | 884027 | Immune |
| augustus_masked-000016F-processed-gene-9.67-mRNA-1 | 1 | 000016F | 899346 | 901910 | Immune |
| maker-000016F-augustus-gene-10.83-mRNA-1 | 1 | 000016F | 1056461 | 1058269 | Immune |
| maker-000016F-augustus-gene-13.129-mRNA-4 | 1 | 000016F | 1352267 | 1365480 | Immune |
| genemark-000016F-processed-gene-21.82-mRNA-1 | 1 | 000016F | 2138876 | 2142032 | Immune |
| maker-000016F-augustus-gene-24.47-mRNA-1 | 1 | 000016F | 2477008 | 2479513 | Immune |
| maker-000016F-snap-gene-25.97-mRNA-1 | 1 | 000016F | 2495785 | 2496419 | Immune |
| maker-000000F-augustus-gene-6.147-mRNA-1 | 2 | 000000F | 592955 | 594543 | Immune |
| maker-000000F-snap-gene-6.42-mRNA-1 | 2 | 000000F | 594627 | 597855 | Immune |
| maker-000000F-augustus-gene-6.140-mRNA-1 | 2 | 000000F | 604368 | 605423 | Immune |
| augustus_masked-000000F-processed-gene-6.95-mRNA-1 | 2 | 000000F | 656795 | 657835 | Immune |
| genemark-000000F-processed-gene-7.36-mRNA-1 | 2 | 000000F | 747828 | 749156 | Immune |
| maker-000000F-augustus-gene-9.86-mRNA-1 | 2 | 000000F | 883871 | 889555 | Immune |
| maker-000000F-augustus-gene-9.87-mRNA-1 | 2 | 000000F | 892092 | 896512 | Immune |
| maker-000000F-augustus-gene-9.88-mRNA-1 | 2 | 000000F | 897413 | 903170 | Immune |
| maker-000000F-augustus-gene-10.30-mRNA-1 | 2 | 000000F | 1019076 | 1020838 | Immune |
| maker-000000F-augustus-gene-19.33-mRNA-1 | 2 | 000000F | 1918528 | 1924209 | Immune |
| maker-000000F-augustus-gene-20.50-mRNA-1 | 2 | 000000F | 2050088 | 2053244 | Immune |
| maker-000000F-augustus-gene-23.90-mRNA-1 | 2 | 000000F | 2369072 | 2379488 | Immune |
| maker-000000F-augustus-gene-32.58-mRNA-1 | 2 | 000000F | 3247090 | 3252278 | Immune |
| maker-000000F-augustus-gene-33.18-mRNA-1 | 2 | 000000F | 3357927 | 3363809 | Immune |
| maker-000000F-snap-gene-33.32-mRNA-1 | 2 | 000000F | 3391464 | 3393629 | Immune |
| maker-000000F-snap-gene-35.48-mRNA-1 | 2 | 000000F | 3500685 | 3515203 | Immune |
| maker-000000F-augustus-gene-41.15-mRNA-1 | 2 | 000000F | 4153450 | 4155990 | Immune |
| maker-000000F-snap-gene-44.47-mRNA-1 | 2 | 000000F | 4425102 | 4437276 | Immune |
| maker-000000F-snap-gene-44.47-mRNA-2 | 2 | 000000F | 4425102 | 4437276 | Immune |
| maker-000000F-snap-gene-44.47-mRNA-3 | 2 | 000000F | 4425472 | 4437276 | Immune |
| maker-000000F-augustus-gene-44.82-mRNA-1 | 2 | 000000F | 4447087 | 4449048 | Immune |
| maker-000000F-augustus-gene-46.51-mRNA-1 | 2 | 000000F | 4592356 | 4597524 | Immune |
| genemark-000000F-processed-gene-46.26-mRNA-1 | 2 | 000000F | 4622871 | 4630257 | Immune |
| maker-000000F-snap-gene-50.15-mRNA-1 | 2 | 000000F | 5068561 | 5074805 | Immune |
| maker-000000F-augustus-gene-56.66-mRNA-1 | 2 | 000000F | 5599354 | 5602058 | Immune |
| maker-000000F-augustus-gene-57.54-mRNA-1 | 2 | 000000F | 5693459 | 5694311 | Immune |
| augustus_masked-000000F-processed-gene-57.64-mRNA-1 | 2 | 000000F | 5694908 | 5697656 | Immune |
| augustus_masked-000000F-processed-gene-57.65-mRNA-1 | 2 | 000000F | 5698209 | 5699212 | Immune |
| maker-000000F-augustus-gene-57.45-mRNA-1 | 2 | 000000F | 5703442 | 5704852 | Immune |
| genemark-000000F-processed-gene-57.6-mRNA-1 | 2 | 000000F | 5705823 | 5706536 | Immune |
| maker-000000F-augustus-gene-57.47-mRNA-1 | 2 | 000000F | 5707347 | 5708289 | Immune |
| maker-000000F-augustus-gene-57.50-mRNA-1 | 2 | 000000F | 5751490 | 5753195 | Immune |
| maker-000000F-augustus-gene-57.51-mRNA-1 | 2 | 000000F | 5754899 | 5756133 | Immune |
| maker-000000F-augustus-gene-57.52-mRNA-1 | 2 | 000000F | 5756953 | 5758400 | Immune |
| maker-000000F-snap-gene-67.66-mRNA-1 | 2 | 000000F | 6724496 | 6728238 | Immune |
| maker-000004F-snap-gene-1.45-mRNA-1 | 2 | 000004F | 174795 | 185171 | Immune |
| maker-000004F-augustus-gene-4.5-mRNA-1 | 2 | 000004F | 406819 | 415389 | Immune |
| maker-000004F-snap-gene-14.34-mRNA-1 | 2 | 000004F | 1440771 | 1449270 | Immune |
| maker-000004F-augustus-gene-14.19-mRNA-1 | 2 | 000004F | 1449143 | 1451191 | Immune |
| maker-000004F-augustus-gene-16.82-mRNA-1 | 2 | 000004F | 1604496 | 1614802 | Immune |
| augustus_masked-000004F-processed-gene-16.24-mRNA-1 | 2 | 000004F | 1642417 | 1654990 | Immune |
| maker-000004F-snap-gene-19.36-mRNA-1 | 2 | 000004F | 1872929 | 1889478 | Immune |
| maker-000004F-augustus-gene-22.57-mRNA-1 | 2 | 000004F | 2231376 | 2234393 | Immune |
| genemark-000004F-processed-gene-25.11-mRNA-1 | 2 | 000004F | 2504453 | 2509797 | Immune |
| genemark-000004F-processed-gene-25.13-mRNA-1 | 2 | 000004F | 2514690 | 2519537 | Immune |
| maker-000004F-augustus-gene-27.24-mRNA-1 | 2 | 000004F | 2699274 | 2705725 | Immune |
| maker-000004F-augustus-gene-27.27-mRNA-1 | 2 | 000004F | 2726198 | 2727409 | Immune |
| maker-000004F-augustus-gene-27.31-mRNA-1 | 2 | 000004F | 2737422 | 2739436 | Immune |
| augustus_masked-000004F-processed-gene-37.30-mRNA-1 | 2 | 000004F | 3688545 | 3694412 | Immune |
| genemark-000004F-processed-gene-37.78-mRNA-1 | 2 | 000004F | 3782865 | 3788303 | Immune |
| maker-000002F-snap-gene-1.65-mRNA-1 | 3 | 000002F | 117228 | 134541 | Immune |
| maker-000002F-augustus-gene-3.34-mRNA-1 | 3 | 000002F | 295104 | 296534 | Immune |
| augustus_masked-000002F-processed-gene-4.69-mRNA-1 | 3 | 000002F | 431237 | 435310 | Immune |
| maker-000002F-augustus-gene-9.62-mRNA-1 | 3 | 000002F | 927454 | 941177 | Immune |
| augustus_masked-000002F-processed-gene-17.3-mRNA-1 | 3 | 000002F | 1734075 | 1735068 | Immune |
| maker-000002F-augustus-gene-24.52-mRNA-1 | 3 | 000002F | 2457406 | 2466194 | Immune |
| maker-000002F-augustus-gene-40.51-mRNA-1 | 3 | 000002F | 4021127 | 4022382 | Immune |
| maker-000009F-augustus-gene-11.3-mRNA-1 | 3 | 000009F | 1166001 | 1171799 | Immune |
| maker-000009F-augustus-gene-11.4-mRNA-1 | 3 | 000009F | 1171857 | 1175052 | Immune |
| genemark-000009F-processed-gene-17.63-mRNA-1 | 3 | 000009F | 1745440 | 1752301 | Immune |
| augustus_masked-000011F-processed-gene-0.66-mRNA-1 | 4 | 000011F | 76360 | 82506 | Immune |
| maker-000011F-snap-gene-0.21-mRNA-1 | 4 | 000011F | 7012 | 10491 | Immune |
| genemark-000011F-processed-gene-2.50-mRNA-1 | 4 | 000011F | 266694 | 272096 | Immune |
| genemark-000011F-processed-gene-2.51-mRNA-1 | 4 | 000011F | 273527 | 274950 | Immune |
| maker-000011F-augustus-gene-5.93-mRNA-1 | 4 | 000011F | 552209 | 559903 | Immune |
| maker-000011F-augustus-gene-6.28-mRNA-1 | 4 | 000011F | 649093 | 650962 | Immune |
| maker-000011F-snap-gene-7.125-mRNA-1 | 4 | 000011F | 731482 | 733129 | Immune |
| maker-000011F-snap-gene-7.116-mRNA-1 | 4 | 000011F | 751791 | 757106 | Immune |
| maker-000011F-snap-gene-7.116-mRNA-2 | 4 | 000011F | 753132 | 757106 | Immune |
| augustus_masked-000011F-processed-gene-8.58-mRNA-1 | 4 | 000011F | 866191 | 868974 | Immune |
| maker-000011F-augustus-gene-14.28-mRNA-2 | 4 | 000011F | 1465519 | 1478828 | Immune |
| maker-000011F-augustus-gene-14.38-mRNA-1 | 4 | 000011F | 1470380 | 1471938 | Immune |
| maker-000011F-augustus-gene-16.30-mRNA-1 | 4 | 000011F | 1677583 | 1680767 | Immune |
| maker-000011F-augustus-gene-19.79-mRNA-1 | 4 | 000011F | 1963256 | 1966250 | Immune |
| maker-000011F-augustus-gene-19.79-mRNA-3 | 4 | 000011F | 1963256 | 1966293 | Immune |
| maker-000011F-snap-gene-21.42-mRNA-1 | 4 | 000011F | 2138669 | 2139714 | Immune |
| genemark-000011F-processed-gene-21.35-mRNA-1 | 4 | 000011F | 2140257 | 2142146 | Immune |
| maker-000011F-augustus-gene-22.102-mRNA-1 | 4 | 000011F | 2225276 | 2227525 | Immune |
| maker-000011F-augustus-gene-22.97-mRNA-1 | 4 | 000011F | 2260336 | 2262148 | Immune |
| augustus_masked-000011F-processed-gene-22.90-mRNA-1 | 4 | 000011F | 2263084 | 2270489 | Immune |
| snap_masked-000011F-processed-gene-22.25-mRNA-1 | 4 | 000011F | 2276641 | 2277951 | Immune |
| maker-000011F-snap-gene-23.57-mRNA-1 | 4 | 000011F | 2293574 | 2294994 | Immune |
| maker-000011F-augustus-gene-23.92-mRNA-1 | 4 | 000011F | 2330095 | 2331989 | Immune |
| maker-000011F-augustus-gene-23.93-mRNA-1 | 4 | 000011F | 2338421 | 2339365 | Immune |
| maker-000011F-augustus-gene-23.94-mRNA-1 | 4 | 000011F | 2340267 | 2342739 | Immune |
| maker-000011F-snap-gene-23.73-mRNA-1 | 4 | 000011F | 2343207 | 2344704 | Immune |
| augustus_masked-000011F-processed-gene-23.23-mRNA-1 | 4 | 000011F | 2351137 | 2352118 | Immune |
| augustus_masked-000011F-processed-gene-23.10-mRNA-1 | 4 | 000011F | 2354215 | 2355144 | Immune |
| genemark-000011F-processed-gene-23.53-mRNA-1 | 4 | 000011F | 2355579 | 2356842 | Immune |
| maker-000011F-augustus-gene-23.98-mRNA-1 | 4 | 000011F | 2357556 | 2359103 | Immune |
| maker-000011F-snap-gene-24.30-mRNA-1 | 4 | 000011F | 2382761 | 2396586 | Immune |
| augustus_masked-000011F-processed-gene-24.42-mRNA-1 | 4 | 000011F | 2464139 | 2470650 | Immune |
| augustus_masked-000011F-processed-gene-27.51-mRNA-1 | 4 | 000011F | 2726703 | 2731947 | Immune |
| maker-000011F-augustus-gene-27.134-mRNA-1 | 4 | 000011F | 2747744 | 2750430 | Immune |
| maker-000011F-augustus-gene-27.135-mRNA-1 | 4 | 000011F | 2750928 | 2755888 | Immune |
| maker-000011F-augustus-gene-27.127-mRNA-1 | 4 | 000011F | 2791648 | 2793477 | Immune |
| maker-000011F-augustus-gene-28.175-mRNA-1 | 4 | 000011F | 2799370 | 2802674 | Immune |
| augustus_masked-000011F-processed-gene-28.77-mRNA-1 | 4 | 000011F | 2807906 | 2824942 | Immune |
| maker-000011F-augustus-gene-28.180-mRNA-1 | 4 | 000011F | 2825757 | 2827658 | Immune |
| maker-000011F-augustus-gene-28.157-mRNA-1 | 4 | 000011F | 2874403 | 2877729 | Immune |
| augustus_masked-000011F-processed-gene-28.85-mRNA-1 | 4 | 000011F | 2877757 | 2885909 | Immune |
| genemark-000011F-processed-gene-28.40-mRNA-1 | 4 | 000011F | 2886130 | 2889195 | Immune |
| maker-000011F-augustus-gene-28.164-mRNA-1 | 4 | 000011F | 2926188 | 2927047 | Immune |
| maker-000011F-augustus-gene-28.196-mRNA-1 | 4 | 000011F | 2968764 | 2971724 | Immune |
| maker-000023F-augustus-gene-0.39-mRNA-1 | 4 | 000023F | 39044 | 40564 | Immune |
| maker-000023F-snap-gene-0.68-mRNA-1 | 4 | 000023F | 41487 | 45862 | Immune |
| maker-000023F-snap-gene-0.70-mRNA-1 | 4 | 000023F | 65987 | 71700 | Immune |
| maker-000023F-augustus-gene-1.49-mRNA-1 | 4 | 000023F | 90316 | 106882 | Immune |
| genemark-000023F-processed-gene-1.37-mRNA-1 | 4 | 000023F | 165851 | 173791 | Immune |
| maker-000023F-augustus-gene-5.12-mRNA-1 | 4 | 000023F | 533761 | 544194 | Immune |
| maker-000023F-augustus-gene-9.101-mRNA-1 | 4 | 000023F | 964209 | 966519 | Immune |
| maker-000023F-snap-gene-9.38-mRNA-1 | 4 | 000023F | 968756 | 969347 | Immune |
| maker-000026F-augustus-gene-0.83-mRNA-1 | 4 | 000026F | 33041 | 39382 | Immune |
| augustus_masked-000026F-processed-gene-0.45-mRNA-1 | 4 | 000026F | 39920 | 45204 | Immune |
| maker-000026F-augustus-gene-0.91-mRNA-1 | 4 | 000026F | 45479 | 46971 | Immune |
| maker-000026F-snap-gene-0.37-mRNA-1 | 4 | 000026F | 54872 | 59400 | Immune |
| maker-000026F-augustus-gene-0.96-mRNA-1 | 4 | 000026F | 73444 | 82759 | Immune |
| maker-000026F-augustus-gene-3.63-mRNA-1 | 4 | 000026F | 262747 | 275803 | Immune |
| maker-000026F-augustus-gene-7.89-mRNA-1 | 4 | 000026F | 715655 | 721787 | Immune |
| maker-000028F-snap-gene-0.79-mRNA-1 | 4 | 000028F | 88037 | 89004 | Immune |
| genemark-000028F-processed-gene-2.42-mRNA-1 | 4 | 000028F | 201656 | 204334 | Immune |
| maker-000028F-augustus-gene-2.72-mRNA-1 | 4 | 000028F | 213373 | 223087 | Immune |
| genemark-000028F-processed-gene-2.44-mRNA-1 | 4 | 000028F | 214099 | 215223 | Immune |
| maker-000028F-snap-gene-2.104-mRNA-1 | 4 | 000028F | 239879 | 243727 | Immune |
| maker-000028F-augustus-gene-2.94-mRNA-1 | 4 | 000028F | 276436 | 280105 | Immune |
| maker-000028F-snap-gene-3.117-mRNA-1 | 4 | 000028F | 284792 | 288007 | Immune |
| snap_masked-000028F-processed-gene-3.3-mRNA-1 | 4 | 000028F | 297474 | 302657 | Immune |
| maker-000028F-snap-gene-3.109-mRNA-1 | 4 | 000028F | 304402 | 310273 | Immune |
| maker-000028F-snap-gene-3.126-mRNA-1 | 4 | 000028F | 342530 | 349674 | Immune |
| genemark-000028F-processed-gene-3.44-mRNA-1 | 4 | 000028F | 377341 | 379989 | Immune |
| maker-000028F-augustus-gene-4.61-mRNA-1 | 4 | 000028F | 484714 | 486190 | Immune |
| genemark-000028F-processed-gene-5.1-mRNA-1 | 4 | 000028F | 501014 | 505639 | Immune |
| maker-000028F-augustus-gene-6.43-mRNA-1 | 4 | 000028F | 665263 | 672009 | Immune |
| maker-000028F-augustus-gene-11.83-mRNA-1 | 4 | 000028F | 1129041 | 1130127 | Immune |
| maker-000028F-augustus-gene-13.126-mRNA-1 | 4 | 000028F | 1380392 | 1386515 | Immune |
| maker-000028F-augustus-gene-14.130-mRNA-1 | 4 | 000028F | 1400208 | 1406329 | Immune |
| snap_masked-000028F-processed-gene-14.174-mRNA-1 | 4 | 000028F | 1408110 | 1413839 | Immune |
| maker-000028F-snap-gene-14.25-mRNA-1 | 4 | 000028F | 1419234 | 1426554 | Immune |
| genemark-000028F-processed-gene-14.67-mRNA-1 | 4 | 000028F | 1439482 | 1443738 | Immune |
| maker-000001F-augustus-gene-5.103-mRNA-1 | 5 | 000001F | 590572 | 591510 | Immune |
| maker-000001F-augustus-gene-13.40-mRNA-1 | 5 | 000001F | 1289074 | 1291397 | Immune |
| genemark-000001F-processed-gene-13.19-mRNA-1 | 5 | 000001F | 1332544 | 1334189 | Immune |
| maker-000001F-augustus-gene-20.38-mRNA-1 | 5 | 000001F | 2059751 | 2063780 | Immune |
| maker-000001F-augustus-gene-24.25-mRNA-1 | 5 | 000001F | 2409448 | 2422310 | Immune |
| augustus_masked-000001F-processed-gene-26.20-mRNA-1 | 5 | 000001F | 2620504 | 2622583 | Immune |
| maker-000001F-snap-gene-26.50-mRNA-1 | 5 | 000001F | 2625940 | 2630694 | Immune |
| maker-000001F-snap-gene-29.79-mRNA-1 | 5 | 000001F | 2960339 | 2981179 | Immune |
| genemark-000001F-processed-gene-33.71-mRNA-1 | 5 | 000001F | 3376285 | 3377523 | Immune |
| maker-000001F-snap-gene-35.20-mRNA-1 | 5 | 000001F | 3509961 | 3515325 | Immune |
| maker-000001F-augustus-gene-35.71-mRNA-1 | 5 | 000001F | 3554903 | 3556005 | Immune |
| maker-000001F-snap-gene-37.37-mRNA-1 | 5 | 000001F | 3731617 | 3737398 | Immune |
| maker-000001F-augustus-gene-47.67-mRNA-1 | 5 | 000001F | 4778595 | 4782425 | Immune |
| maker-000001F-augustus-gene-48.53-mRNA-1 | 5 | 000001F | 4805198 | 4809306 | Immune |
| maker-000001F-augustus-gene-48.69-mRNA-1 | 5 | 000001F | 4882752 | 4885356 | Immune |
| maker-000074F-augustus-gene-2.76-mRNA-1 | 5 | 000074F | 230815 | 257085 | Immune |
| maker-000108F-augustus-gene-0.66-mRNA-1 | 5 | 000108F | 69916 | 70876 | Immune |
| maker-000108F-augustus-gene-0.59-mRNA-1 | 5 | 000108F | 113392 | 125151 | Immune |
| maker-000012F-augustus-gene-20.59-mRNA-1 | 6 | 000012F | 2030881 | 2033408 | Immune |
| augustus_masked-000012F-processed-gene-21.62-mRNA-1 | 6 | 000012F | 2155676 | 2162064 | Immune |
| maker-000012F-augustus-gene-21.79-mRNA-1 | 6 | 000012F | 2177476 | 2187561 | Immune |
| genemark-000012F-processed-gene-22.38-mRNA-1 | 6 | 000012F | 2185762 | 2187143 | Immune |
| maker-000012F-snap-gene-25.49-mRNA-1 | 6 | 000012F | 2492085 | 2499156 | Immune |
| maker-000012F-snap-gene-25.51-mRNA-1 | 6 | 000012F | 2507383 | 2509937 | Immune |
| maker-000021F-augustus-gene-4.118-mRNA-1 | 6 | 000021F | 448583 | 450077 | Immune |
| maker-000021F-augustus-gene-15.17-mRNA-1 | 6 | 000021F | 1497388 | 1503377 | Immune |
| maker-000021F-augustus-gene-20.128-mRNA-1 | 6 | 000021F | 2074679 | 2091233 | Immune |
| maker-000052F-snap-gene-0.80-mRNA-1 | 6 | 000052F | 13418 | 23361 | Immune |
| maker-000058F-augustus-gene-2.75-mRNA-1 | 6 | 000058F | 206728 | 212090 | Immune |
| maker-000058F-augustus-gene-2.77-mRNA-1 | 6 | 000058F | 217789 | 226223 | Immune |
| augustus_masked-000058F-processed-gene-3.70-mRNA-1 | 6 | 000058F | 346568 | 347227 | Immune |
| maker-000073F-augustus-gene-0.82-mRNA-1 | 6 | 000073F | 66482 | 69167 | Immune |
| maker-000073F-augustus-gene-1.71-mRNA-1 | 6 | 000073F | 101405 | 102551 | Immune |
| maker-000073F-snap-gene-1.7-mRNA-1 | 6 | 000073F | 124835 | 126475 | Immune |
| maker-000082F-augustus-gene-1.94-mRNA-1 | 6 | 000082F | 170689 | 171667 | Immune |
| maker-000003F-augustus-gene-11.39-mRNA-1 | 7 | 000003F | 1162274 | 1166478 | Immune |
| maker-000003F-augustus-gene-15.97-mRNA-1 | 7 | 000003F | 1589607 | 1591721 | Immune |
| maker-000003F-augustus-gene-21.63-mRNA-1 | 7 | 000003F | 2147844 | 2156020 | Immune |
| genemark-000003F-processed-gene-24.12-mRNA-1 | 7 | 000003F | 2416487 | 2435833 | Immune |
| genemark-000003F-processed-gene-27.86-mRNA-1 | 7 | 000003F | 2748297 | 2753637 | Immune |
| maker-000003F-augustus-gene-30.58-mRNA-1 | 7 | 000003F | 3039376 | 3047981 | Immune |
| genemark-000003F-processed-gene-31.6-mRNA-1 | 7 | 000003F | 3153319 | 3158796 | Immune |
| maker-000003F-augustus-gene-33.49-mRNA-1 | 7 | 000003F | 3330833 | 3344534 | Immune |
| maker-000003F-augustus-gene-33.58-mRNA-1 | 7 | 000003F | 3358222 | 3360624 | Immune |
| maker-000003F-snap-gene-35.76-mRNA-1 | 7 | 000003F | 3500399 | 3507515 | Immune |
| genemark-000003F-processed-gene-36.49-mRNA-1 | 7 | 000003F | 3587049 | 3590031 | Immune |
| maker-000003F-augustus-gene-36.86-mRNA-1 | 7 | 000003F | 3621088 | 3622796 | Immune |
| maker-000003F-snap-gene-38.67-mRNA-1 | 7 | 000003F | 3819182 | 3825102 | Immune |
| maker-000003F-snap-gene-38.69-mRNA-1 | 7 | 000003F | 3836815 | 3846942 | Immune |
| maker-000003F-augustus-gene-40.30-mRNA-1 | 7 | 000003F | 4003948 | 4005793 | Immune |
| maker-000003F-snap-gene-40.74-mRNA-1 | 7 | 000003F | 4006516 | 4026331 | Immune |
| maker-000003F-augustus-gene-40.22-mRNA-1 | 7 | 000003F | 4048649 | 4062633 | Immune |
| genemark-000003F-processed-gene-42.55-mRNA-1 | 7 | 000003F | 4216155 | 4216652 | Immune |
| augustus_masked-000018F-processed-gene-2.21-mRNA-1 | 7 | 000018F | 233618 | 238770 | Immune |
| maker-000018F-augustus-gene-3.53-mRNA-1 | 7 | 000018F | 297421 | 310076 | Immune |
| augustus_masked-000018F-processed-gene-10.79-mRNA-1 | 7 | 000018F | 1009897 | 1012555 | Immune |
| augustus_masked-000018F-processed-gene-10.68-mRNA-1 | 7 | 000018F | 1013389 | 1016614 | Immune |
| augustus_masked-000018F-processed-gene-10.69-mRNA-1 | 7 | 000018F | 1017172 | 1020481 | Immune |
| maker-000018F-augustus-gene-10.50-mRNA-1 | 7 | 000018F | 1020578 | 1022156 | Immune |
| augustus_masked-000018F-processed-gene-10.71-mRNA-1 | 7 | 000018F | 1023100 | 1024492 | Immune |
| augustus_masked-000018F-processed-gene-12.31-mRNA-1 | 7 | 000018F | 1283529 | 1284322 | Immune |
| maker-000018F-augustus-gene-13.65-mRNA-1 | 7 | 000018F | 1284666 | 1285856 | Immune |
| maker-000018F-augustus-gene-13.66-mRNA-1 | 7 | 000018F | 1286214 | 1287131 | Immune |
| augustus_masked-000018F-processed-gene-13.28-mRNA-1 | 7 | 000018F | 1311141 | 1314050 | Immune |
| maker-000018F-augustus-gene-14.99-mRNA-1 | 7 | 000018F | 1434586 | 1436024 | Immune |
| maker-000018F-augustus-gene-14.100-mRNA-1 | 7 | 000018F | 1437028 | 1438704 | Immune |
| maker-000018F-snap-gene-15.31-mRNA-1 | 7 | 000018F | 1506215 | 1507722 | Immune |
| maker-000018F-augustus-gene-16.117-mRNA-1 | 7 | 000018F | 1620389 | 1625587 | Immune |
| maker-000018F-augustus-gene-16.117-mRNA-2 | 7 | 000018F | 1620542 | 1625587 | Immune |
| maker-000018F-augustus-gene-16.118-mRNA-1 | 7 | 000018F | 1631064 | 1632124 | Immune |
| genemark-000018F-processed-gene-16.61-mRNA-1 | 7 | 000018F | 1632887 | 1634322 | Immune |
| snap_masked-000018F-processed-gene-16.12-mRNA-1 | 7 | 000018F | 1646457 | 1647683 | Immune |
| maker-000018F-snap-gene-16.153-mRNA-1 | 7 | 000018F | 1647843 | 1648729 | Immune |
| maker-000018F-augustus-gene-16.102-mRNA-1 | 7 | 000018F | 1649368 | 1650340 | Immune |
| maker-000018F-augustus-gene-16.124-mRNA-1 | 7 | 000018F | 1660104 | 1661170 | Immune |
| augustus_masked-000018F-processed-gene-16.95-mRNA-1 | 7 | 000018F | 1661549 | 1665824 | Immune |
| augustus_masked-000018F-processed-gene-16.79-mRNA-1 | 7 | 000018F | 1667781 | 1670434 | Immune |
| genemark-000018F-processed-gene-16.50-mRNA-1 | 7 | 000018F | 1670589 | 1674954 | Immune |
| maker-000018F-augustus-gene-17.59-mRNA-1 | 7 | 000018F | 1793419 | 1795761 | Immune |
| maker-000018F-augustus-gene-18.10-mRNA-1 | 7 | 000018F | 1799876 | 1801441 | Immune |
| maker-000018F-augustus-gene-18.11-mRNA-1 | 7 | 000018F | 1803421 | 1805089 | Immune |
| augustus_masked-000018F-processed-gene-18.30-mRNA-1 | 7 | 000018F | 1812352 | 1817411 | Immune |
| maker-000018F-snap-gene-18.70-mRNA-1 | 7 | 000018F | 1867491 | 1873893 | Immune |
| maker-000018F-snap-gene-19.97-mRNA-1 | 7 | 000018F | 1903143 | 1907298 | Immune |
| augustus_masked-000018F-processed-gene-19.120-mRNA-1 | 7 | 000018F | 1910248 | 1911822 | Immune |
| maker-000018F-snap-gene-19.86-mRNA-1 | 7 | 000018F | 1913566 | 1921528 | Immune |
| maker-000018F-augustus-gene-19.48-mRNA-1 | 7 | 000018F | 1914070 | 1922405 | Immune |
| maker-000018F-snap-gene-19.90-mRNA-1 | 7 | 000018F | 1931429 | 1936768 | Immune |
| maker-000018F-augustus-gene-20.127-mRNA-1 | 7 | 000018F | 2072969 | 2074947 | Immune |
| snap_masked-000018F-processed-gene-21.6-mRNA-1 | 7 | 000018F | 2129436 | 2130641 | Immune |
| genemark-000018F-processed-gene-21.63-mRNA-1 | 7 | 000018F | 2146305 | 2148608 | Immune |
| maker-000018F-augustus-gene-21.121-mRNA-1 | 7 | 000018F | 2153463 | 2154330 | Immune |
| maker-000018F-augustus-gene-21.122-mRNA-1 | 7 | 000018F | 2160480 | 2162692 | Immune |
| maker-000018F-augustus-gene-21.123-mRNA-1 | 7 | 000018F | 2163391 | 2164538 | Immune |
| maker-000018F-augustus-gene-22.75-mRNA-1 | 7 | 000018F | 2243854 | 2244831 | Immune |
| genemark-000018F-processed-gene-23.56-mRNA-1 | 7 | 000018F | 2302373 | 2304423 | Immune |
| genemark-000018F-processed-gene-23.70-mRNA-1 | 7 | 000018F | 2365125 | 2372926 | Immune |
| augustus_masked-000018F-processed-gene-24.179-mRNA-1 | 7 | 000018F | 2427104 | 2428997 | Immune |
| maker-000018F-snap-gene-24.84-mRNA-1 | 7 | 000018F | 2447559 | 2461187 | Immune |
| maker-000018F-augustus-gene-24.234-mRNA-1 | 7 | 000018F | 2534802 | 2545559 | Immune |
| maker-000038F-augustus-gene-2.133-mRNA-1 | 7 | 000038F | 235532 | 238485 | Immune |
| augustus_masked-000038F-processed-gene-2.75-mRNA-1 | 7 | 000038F | 238647 | 245968 | Immune |
| maker-000038F-augustus-gene-3.45-mRNA-1 | 7 | 000038F | 332304 | 333359 | Immune |
| snap_masked-000038F-processed-gene-3.121-mRNA-1 | 7 | 000038F | 381649 | 382839 | Immune |
| genemark-000038F-processed-gene-3.13-mRNA-1 | 7 | 000038F | 384859 | 389600 | Immune |
| maker-000038F-augustus-gene-4.136-mRNA-1 | 7 | 000038F | 394027 | 395845 | Immune |
| maker-000038F-snap-gene-4.117-mRNA-1 | 7 | 000038F | 419632 | 424478 | Immune |
| maker-000038F-augustus-gene-4.127-mRNA-1 | 7 | 000038F | 423590 | 426734 | Immune |
| snap_masked-000038F-processed-gene-4.28-mRNA-1 | 7 | 000038F | 464192 | 465469 | Immune |
| maker-000038F-augustus-gene-4.143-mRNA-1 | 7 | 000038F | 471977 | 474418 | Immune |
| augustus_masked-000038F-processed-gene-4.95-mRNA-1 | 7 | 000038F | 474559 | 484144 | Immune |
| maker-000038F-augustus-gene-5.44-mRNA-1 | 7 | 000038F | 495613 | 496259 | Immune |
| snap_masked-000038F-processed-gene-5.17-mRNA-1 | 7 | 000038F | 569838 | 570974 | Immune |
| augustus_masked-000038F-processed-gene-8.98-mRNA-1 | 7 | 000038F | 867776 | 876473 | Immune |
| genemark-000014F-processed-gene-15.11-mRNA-1 | 9 | 000014F | 1492885 | 1500335 | Immune |
| maker-000014F-augustus-gene-18.31-mRNA-1 | 9 | 000014F | 1813417 | 1824673 | Immune |
| maker-000014F-augustus-gene-19.26-mRNA-1 | 9 | 000014F | 1968243 | 1978198 | Immune |
| maker-000014F-augustus-gene-20.57-mRNA-1 | 9 | 000014F | 2035576 | 2041634 | Immune |
| maker-000014F-augustus-gene-20.59-mRNA-1 | 9 | 000014F | 2076803 | 2078861 | Immune |
| augustus_masked-000014F-processed-gene-21.8-mRNA-1 | 9 | 000014F | 2132131 | 2138061 | Immune |
| maker-000014F-augustus-gene-23.5-mRNA-1 | 9 | 000014F | 2296928 | 2298409 | Immune |
| maker-000014F-augustus-gene-25.57-mRNA-1 | 9 | 000014F | 2528135 | 2532405 | Immune |
| maker-000081F-augustus-gene-1.86-mRNA-1 | 9 | 000081F | 195285 | 208194 | Immune |
| augustus_masked-000081F-processed-gene-1.22-mRNA-1 | 9 | 000081F | 208779 | 215521 | Immune |
| maker-000006F-augustus-gene-9.89-mRNA-1 | 10 | 000006F | 892327 | 911345 | Immune |
| maker-000006F-augustus-gene-13.42-mRNA-1 | 10 | 000006F | 1314285 | 1321114 | Immune |
| maker-000006F-augustus-gene-26.63-mRNA-1 | 10 | 000006F | 2666245 | 2667621 | Immune |
| augustus_masked-000006F-processed-gene-26.23-mRNA-1 | 10 | 000006F | 2667543 | 2680139 | Immune |
| maker-000006F-augustus-gene-27.28-mRNA-1 | 10 | 000006F | 2760659 | 2763367 | Immune |
| maker-000006F-augustus-gene-30.145-mRNA-1 | 10 | 000006F | 2961458 | 2962799 | Immune |
| augustus_masked-000006F-processed-gene-30.176-mRNA-1 | 10 | 000006F | 2963351 | 2963701 | Immune |
| snap_masked-000006F-processed-gene-30.23-mRNA-1 | 10 | 000006F | 3022666 | 3023450 | Immune |
| maker-000006F-augustus-gene-30.135-mRNA-1 | 10 | 000006F | 3037874 | 3039558 | Immune |
| maker-000006F-augustus-gene-30.165-mRNA-1 | 10 | 000006F | 3058141 | 3059235 | Immune |
| genemark-000006F-processed-gene-33.52-mRNA-1 | 10 | 000006F | 3360449 | 3373410 | Immune |
| maker-000013F-snap-gene-7.63-mRNA-1 | 10 | 000013F | 724438 | 728371 | Immune |
| maker-000013F-snap-gene-7.68-mRNA-1 | 10 | 000013F | 752031 | 755971 | Immune |
| maker-000013F-snap-gene-9.78-mRNA-1 | 10 | 000013F | 935799 | 939641 | Immune |
| augustus_masked-000013F-processed-gene-9.72-mRNA-1 | 10 | 000013F | 961576 | 973039 | Immune |
| maker-000013F-augustus-gene-10.73-mRNA-1 | 10 | 000013F | 1001537 | 1003055 | Immune |
| maker-000013F-snap-gene-10.65-mRNA-1 | 10 | 000013F | 1077341 | 1080582 | Immune |
| maker-000013F-augustus-gene-10.80-mRNA-1 | 10 | 000013F | 1085192 | 1085748 | Immune |
| augustus_masked-000013F-processed-gene-11.84-mRNA-1 | 10 | 000013F | 1104513 | 1105806 | Immune |
| augustus_masked-000013F-processed-gene-11.85-mRNA-1 | 10 | 000013F | 1117203 | 1118550 | Immune |
| maker-000013F-augustus-gene-11.108-mRNA-1 | 10 | 000013F | 1126660 | 1129520 | Immune |
| maker-000013F-snap-gene-11.45-mRNA-1 | 10 | 000013F | 1129787 | 1135054 | Immune |
| maker-000013F-augustus-gene-16.6-mRNA-1 | 10 | 000013F | 1632463 | 1634740 | Immune |
| genemark-000013F-processed-gene-16.101-mRNA-1 | 10 | 000013F | 1634899 | 1639832 | Immune |
| maker-000013F-augustus-gene-26.3-mRNA-1 | 10 | 000013F | 2659036 | 2660750 | Immune |
| maker-000008F-snap-gene-0.54-mRNA-1 | 1 | 000008F | 3245 | 4903 | BUSCO |
| maker-000008F-augustus-gene-2.83-mRNA-1 | 1 | 000008F | 259080 | 260224 | BUSCO |
| maker-000008F-augustus-gene-4.47-mRNA-2 | 1 | 000008F | 484534 | 486306 | BUSCO |
| genemark-000008F-processed-gene-5.52-mRNA-1 | 1 | 000008F | 493748 | 496235 | BUSCO |
| maker-000008F-augustus-gene-5.21-mRNA-1 | 1 | 000008F | 495750 | 496868 | BUSCO |
| snap_masked-000008F-processed-gene-7.3-mRNA-1 | 1 | 000008F | 783612 | 787685 | BUSCO |
| maker-000008F-snap-gene-10.42-mRNA-1 | 1 | 000008F | 1078030 | 1079583 | BUSCO |
| maker-000008F-augustus-gene-12.40-mRNA-1 | 1 | 000008F | 1219435 | 1221668 | BUSCO |
| maker-000008F-augustus-gene-14.40-mRNA-1 | 1 | 000008F | 1404559 | 1409194 | BUSCO |
| maker-000008F-augustus-gene-14.36-mRNA-1 | 1 | 000008F | 1465516 | 1466815 | BUSCO |
| genemark-000008F-processed-gene-15.42-mRNA-1 | 1 | 000008F | 1556581 | 1558986 | BUSCO |
| augustus_masked-000008F-processed-gene-15.29-mRNA-1 | 1 | 000008F | 1559210 | 1576206 | BUSCO |
| maker-000008F-augustus-gene-16.61-mRNA-3 | 1 | 000008F | 1607285 | 1611081 | BUSCO |
| maker-000008F-augustus-gene-16.74-mRNA-1 | 1 | 000008F | 1643132 | 1643577 | BUSCO |
| maker-000008F-snap-gene-16.51-mRNA-3 | 1 | 000008F | 1650558 | 1652370 | BUSCO |
| maker-000008F-augustus-gene-18.20-mRNA-2 | 1 | 000008F | 1842494 | 1845632 | BUSCO |
| maker-000008F-augustus-gene-19.32-mRNA-1 | 1 | 000008F | 1907007 | 1909222 | BUSCO |
| maker-000008F-augustus-gene-19.34-mRNA-1 | 1 | 000008F | 1916685 | 1917786 | BUSCO |
| maker-000008F-augustus-gene-23.36-mRNA-1 | 1 | 000008F | 2319786 | 2323416 | BUSCO |
| genemark-000008F-processed-gene-29.91-mRNA-1 | 1 | 000008F | 2973432 | 2975781 | BUSCO |
| genemark-000008F-processed-gene-29.93-mRNA-1 | 1 | 000008F | 2990409 | 2994624 | BUSCO |
| augustus_masked-000010F-processed-gene-1.72-mRNA-1 | 1 | 000010F | 178143 | 191996 | BUSCO |
| maker-000010F-augustus-gene-4.51-mRNA-3 | 1 | 000010F | 403801 | 408418 | BUSCO |
| maker-000010F-augustus-gene-4.46-mRNA-1 | 1 | 000010F | 433321 | 436231 | BUSCO |
| genemark-000010F-processed-gene-9.70-mRNA-1 | 1 | 000010F | 911268 | 922396 | BUSCO |
| maker-000010F-augustus-gene-11.59-mRNA-2 | 1 | 000010F | 1091830 | 1094556 | BUSCO |
| augustus_masked-000010F-processed-gene-11.29-mRNA-1 | 1 | 000010F | 1156025 | 1166517 | BUSCO |
| genemark-000010F-processed-gene-12.38-mRNA-1 | 1 | 000010F | 1203759 | 1208880 | BUSCO |
| maker-000010F-augustus-gene-12.78-mRNA-1 | 1 | 000010F | 1212239 | 1214053 | BUSCO |
| genemark-000010F-processed-gene-12.40-mRNA-1 | 1 | 000010F | 1214964 | 1221081 | BUSCO |
| maker-000010F-snap-gene-12.113-mRNA-1 | 1 | 000010F | 1286577 | 1289705 | BUSCO |
| augustus_masked-000010F-processed-gene-13.70-mRNA-1 | 1 | 000010F | 1345363 | 1348170 | BUSCO |
| augustus_masked-000010F-processed-gene-13.59-mRNA-1 | 1 | 000010F | 1345490 | 1346299 | BUSCO |
| maker-000010F-augustus-gene-13.109-mRNA-1 | 1 | 000010F | 1363373 | 1366712 | BUSCO |
| maker-000010F-augustus-gene-16.83-mRNA-2 | 1 | 000010F | 1654059 | 1657336 | BUSCO |
| maker-000010F-augustus-gene-17.86-mRNA-2 | 1 | 000010F | 1754957 | 1756947 | BUSCO |
| genemark-000010F-processed-gene-18.14-mRNA-1 | 1 | 000010F | 1826747 | 1828495 | BUSCO |
| maker-000010F-augustus-gene-18.45-mRNA-3 | 1 | 000010F | 1837915 | 1843247 | BUSCO |
| augustus_masked-000010F-processed-gene-20.0-mRNA-1 | 1 | 000010F | 2002701 | 2016593 | BUSCO |
| maker-000010F-augustus-gene-20.64-mRNA-1 | 1 | 000010F | 2017063 | 2017776 | BUSCO |
| maker-000010F-augustus-gene-20.69-mRNA-1 | 1 | 000010F | 2018040 | 2020162 | BUSCO |
| augustus_masked-000010F-processed-gene-20.16-mRNA-1 | 1 | 000010F | 2068843 | 2072145 | BUSCO |
| maker-000010F-augustus-gene-21.58-mRNA-1 | 1 | 000010F | 2142637 | 2144257 | BUSCO |
| maker-000010F-augustus-gene-22.66-mRNA-1 | 1 | 000010F | 2249622 | 2253306 | BUSCO |
| maker-000010F-augustus-gene-23.74-mRNA-1 | 1 | 000010F | 2300128 | 2301444 | BUSCO |
| maker-000010F-augustus-gene-23.83-mRNA-1 | 1 | 000010F | 2303805 | 2305211 | BUSCO |
| maker-000010F-augustus-gene-23.75-mRNA-1 | 1 | 000010F | 2305362 | 2306227 | BUSCO |
| maker-000010F-augustus-gene-23.77-mRNA-1 | 1 | 000010F | 2327382 | 2330527 | BUSCO |
| maker-000010F-augustus-gene-24.64-mRNA-1 | 1 | 000010F | 2399679 | 2401295 | BUSCO |
| maker-000010F-augustus-gene-24.65-mRNA-1 | 1 | 000010F | 2427316 | 2428719 | BUSCO |
| maker-000010F-augustus-gene-26.54-mRNA-3 | 1 | 000010F | 2645083 | 2650243 | BUSCO |
| maker-000010F-augustus-gene-29.46-mRNA-1 | 1 | 000010F | 2922187 | 2923458 | BUSCO |
| augustus_masked-000016F-processed-gene-0.40-mRNA-1 | 1 | 000016F | 45884 | 52606 | BUSCO |
| maker-000016F-augustus-gene-1.36-mRNA-1 | 1 | 000016F | 93881 | 99401 | BUSCO |
| maker-000016F-snap-gene-1.56-mRNA-3 | 1 | 000016F | 127719 | 129978 | BUSCO |
| maker-000016F-augustus-gene-1.46-mRNA-1 | 1 | 000016F | 129685 | 130756 | BUSCO |
| maker-000016F-augustus-gene-1.50-mRNA-1 | 1 | 000016F | 179951 | 184657 | BUSCO |
| maker-000016F-augustus-gene-3.10-mRNA-1 | 1 | 000016F | 355274 | 356336 | BUSCO |
| maker-000016F-augustus-gene-3.3-mRNA-4 | 1 | 000016F | 356183 | 359952 | BUSCO |
| maker-000016F-augustus-gene-3.6-mRNA-1 | 1 | 000016F | 389181 | 391438 | BUSCO |
| maker-000016F-augustus-gene-4.87-mRNA-1 | 1 | 000016F | 411820 | 413084 | BUSCO |
| maker-000016F-augustus-gene-4.78-mRNA-1 | 1 | 000016F | 414567 | 416552 | BUSCO |
| maker-000016F-augustus-gene-6.56-mRNA-1 | 1 | 000016F | 601456 | 606783 | BUSCO |
| maker-000016F-augustus-gene-8.70-mRNA-1 | 1 | 000016F | 777221 | 778965 | BUSCO |
| snap_masked-000016F-processed-gene-8.144-mRNA-1 | 1 | 000016F | 825939 | 835503 | BUSCO |
| maker-000016F-augustus-gene-8.83-mRNA-1 | 1 | 000016F | 888411 | 891160 | BUSCO |
| genemark-000016F-processed-gene-11.32-mRNA-1 | 1 | 000016F | 1119149 | 1122883 | BUSCO |
| maker-000016F-snap-gene-12.28-mRNA-1 | 1 | 000016F | 1188175 | 1192008 | BUSCO |
| maker-000016F-augustus-gene-13.124-mRNA-1 | 1 | 000016F | 1322649 | 1323214 | BUSCO |
| maker-000016F-augustus-gene-17.15-mRNA-1 | 1 | 000016F | 1707214 | 1710911 | BUSCO |
| maker-000016F-snap-gene-17.73-mRNA-2 | 1 | 000016F | 1748723 | 1754105 | BUSCO |
| genemark-000016F-processed-gene-18.45-mRNA-1 | 1 | 000016F | 1790691 | 1792847 | BUSCO |
| maker-000016F-augustus-gene-18.103-mRNA-2 | 1 | 000016F | 1849792 | 1851551 | BUSCO |
| maker-000016F-augustus-gene-25.93-mRNA-1 | 1 | 000016F | 2597784 | 2600013 | BUSCO |
| maker-000022F-snap-gene-1.27-mRNA-1 | 1 | 000022F | 121283 | 124249 | BUSCO |
| maker-000022F-augustus-gene-2.38-mRNA-1 | 1 | 000022F | 266255 | 267642 | BUSCO |
| maker-000022F-augustus-gene-3.19-mRNA-1 | 1 | 000022F | 366117 | 368522 | BUSCO |
| maker-000022F-augustus-gene-5.69-mRNA-2 | 1 | 000022F | 563451 | 566472 | BUSCO |
| maker-000022F-augustus-gene-6.47-mRNA-2 | 1 | 000022F | 613302 | 615669 | BUSCO |
| genemark-000022F-processed-gene-6.4-mRNA-1 | 1 | 000022F | 626325 | 629396 | BUSCO |
| maker-000022F-augustus-gene-6.52-mRNA-1 | 1 | 000022F | 640203 | 646737 | BUSCO |
| maker-000022F-augustus-gene-6.43-mRNA-1 | 1 | 000022F | 670342 | 671639 | BUSCO |
| augustus_masked-000022F-processed-gene-7.51-mRNA-1 | 1 | 000022F | 741232 | 750544 | BUSCO |
| maker-000022F-augustus-gene-7.83-mRNA-1 | 1 | 000022F | 750586 | 756070 | BUSCO |
| maker-000022F-snap-gene-9.42-mRNA-1 | 1 | 000022F | 908086 | 909943 | BUSCO |
| augustus_masked-000022F-processed-gene-9.27-mRNA-1 | 1 | 000022F | 929155 | 945292 | BUSCO |
| augustus_masked-000022F-processed-gene-11.81-mRNA-1 | 1 | 000022F | 1087971 | 1099675 | BUSCO |
| maker-000022F-augustus-gene-11.69-mRNA-1 | 1 | 000022F | 1124137 | 1127401 | BUSCO |
| maker-000022F-augustus-gene-11.70-mRNA-1 | 1 | 000022F | 1135271 | 1140336 | BUSCO |
| maker-000022F-augustus-gene-12.52-mRNA-1 | 1 | 000022F | 1253453 | 1259402 | BUSCO |
| maker-000022F-augustus-gene-13.106-mRNA-2 | 1 | 000022F | 1295234 | 1297132 | BUSCO |
| maker-000022F-augustus-gene-15.15-mRNA-1 | 1 | 000022F | 1552937 | 1557171 | BUSCO |
| maker-000022F-augustus-gene-16.24-mRNA-1 | 1 | 000022F | 1636860 | 1640634 | BUSCO |
| maker-000022F-augustus-gene-17.75-mRNA-1 | 1 | 000022F | 1676421 | 1683644 | BUSCO |
| genemark-000022F-processed-gene-17.90-mRNA-1 | 1 | 000022F | 1683950 | 1686083 | BUSCO |
| augustus_masked-000022F-processed-gene-16.18-mRNA-1 | 1 | 000022F | 1690738 | 1695344 | BUSCO |
| genemark-000022F-processed-gene-17.94-mRNA-1 | 1 | 000022F | 1695403 | 1696383 | BUSCO |
| maker-000022F-augustus-gene-17.67-mRNA-2 | 1 | 000022F | 1723295 | 1725914 | BUSCO |
| maker-000022F-augustus-gene-17.70-mRNA-1 | 1 | 000022F | 1754766 | 1757678 | BUSCO |
| maker-000022F-augustus-gene-18.18-mRNA-1 | 1 | 000022F | 1808320 | 1810739 | BUSCO |
| augustus_masked-000022F-processed-gene-19.66-mRNA-1 | 1 | 000022F | 1956090 | 1969056 | BUSCO |
| maker-000022F-snap-gene-19.44-mRNA-1 | 1 | 000022F | 1956122 | 1957565 | BUSCO |
| augustus_masked-000022F-processed-gene-19.64-mRNA-1 | 1 | 000022F | 1984131 | 1986757 | BUSCO |
| maker-000022F-augustus-gene-20.89-mRNA-1 | 1 | 000022F | 2018757 | 2021433 | BUSCO |
| maker-000022F-augustus-gene-20.91-mRNA-1 | 1 | 000022F | 2037727 | 2041865 | BUSCO |
| maker-000032F-augustus-gene-6.65-mRNA-1 | 1 | 000032F | 652949 | 659138 | BUSCO |
| maker-000032F-augustus-gene-7.59-mRNA-1 | 1 | 000032F | 720567 | 724349 | BUSCO |
| genemark-000037F-processed-gene-1.36-mRNA-1 | 1 | 000037F | 110884 | 113368 | BUSCO |
| maker-000040F-augustus-gene-6.4-mRNA-2 | 1 | 000040F | 650367 | 653959 | BUSCO |
| maker-000040F-snap-gene-7.50-mRNA-1 | 1 | 000040F | 728776 | 733365 | BUSCO |
| augustus_masked-000047F-processed-gene-1.7-mRNA-1 | 1 | 000047F | 161142 | 164925 | BUSCO |
| maker-000047F-augustus-gene-4.57-mRNA-1 | 1 | 000047F | 475991 | 479098 | BUSCO |
| maker-000047F-snap-gene-5.27-mRNA-1 | 1 | 000047F | 566636 | 569451 | BUSCO |
| augustus_masked-000000F-processed-gene-1.6-mRNA-1 | 2 | 000000F | 156423 | 158149 | BUSCO |
| snap_masked-000000F-processed-gene-1.90-mRNA-1 | 2 | 000000F | 161525 | 162520 | BUSCO |
| maker-000000F-augustus-gene-2.20-mRNA-1 | 2 | 000000F | 244580 | 253269 | BUSCO |
| maker-000000F-augustus-gene-3.40-mRNA-1 | 2 | 000000F | 284927 | 288178 | BUSCO |
| maker-000000F-augustus-gene-5.32-mRNA-1 | 2 | 000000F | 503973 | 505721 | BUSCO |
| maker-000000F-augustus-gene-6.142-mRNA-1 | 2 | 000000F | 611134 | 615247 | BUSCO |
| maker-000000F-augustus-gene-7.11-mRNA-1 | 2 | 000000F | 725151 | 726603 | BUSCO |
| augustus_masked-000000F-processed-gene-8.29-mRNA-1 | 2 | 000000F | 828694 | 830718 | BUSCO |
| augustus_masked-000000F-processed-gene-9.38-mRNA-1 | 2 | 000000F | 935760 | 937628 | BUSCO |
| augustus_masked-000000F-processed-gene-11.33-mRNA-1 | 2 | 000000F | 1081026 | 1082106 | BUSCO |
| augustus_masked-000000F-processed-gene-12.27-mRNA-1 | 2 | 000000F | 1217601 | 1219359 | BUSCO |
| maker-000000F-augustus-gene-12.40-mRNA-1 | 2 | 000000F | 1263822 | 1264585 | BUSCO |
| maker-000000F-augustus-gene-18.41-mRNA-1 | 2 | 000000F | 1817493 | 1821516 | BUSCO |
| maker-000000F-snap-gene-18.26-mRNA-1 | 2 | 000000F | 1848406 | 1861565 | BUSCO |
| genemark-000000F-processed-gene-20.9-mRNA-1 | 2 | 000000F | 2021031 | 2024305 | BUSCO |
| maker-000000F-snap-gene-23.42-mRNA-3 | 2 | 000000F | 2306512 | 2309535 | BUSCO |
| maker-000000F-augustus-gene-23.91-mRNA-2 | 2 | 000000F | 2386732 | 2388069 | BUSCO |
| maker-000000F-augustus-gene-25.40-mRNA-1 | 2 | 000000F | 2585603 | 2587100 | BUSCO |
| maker-000000F-augustus-gene-26.12-mRNA-1 | 2 | 000000F | 2635960 | 2639119 | BUSCO |
| maker-000000F-augustus-gene-28.32-mRNA-2 | 2 | 000000F | 2843361 | 2847802 | BUSCO |
| augustus_masked-000000F-processed-gene-31.19-mRNA-1 | 2 | 000000F | 3113441 | 3126481 | BUSCO |
| augustus_masked-000000F-processed-gene-33.1-mRNA-1 | 2 | 000000F | 3386444 | 3391513 | BUSCO |
| augustus_masked-000000F-processed-gene-35.23-mRNA-1 | 2 | 000000F | 3516635 | 3518315 | BUSCO |
| maker-000000F-augustus-gene-36.72-mRNA-1 | 2 | 000000F | 3646641 | 3647513 | BUSCO |
| maker-000000F-augustus-gene-36.63-mRNA-1 | 2 | 000000F | 3658504 | 3662625 | BUSCO |
| maker-000000F-augustus-gene-37.29-mRNA-3 | 2 | 000000F | 3703954 | 3708738 | BUSCO |
| maker-000000F-augustus-gene-37.25-mRNA-1 | 2 | 000000F | 3708924 | 3710772 | BUSCO |
| maker-000000F-snap-gene-40.25-mRNA-1 | 2 | 000000F | 4022611 | 4029544 | BUSCO |
| augustus_masked-000000F-processed-gene-40.9-mRNA-1 | 2 | 000000F | 4044983 | 4050601 | BUSCO |
| maker-000000F-augustus-gene-41.12-mRNA-1 | 2 | 000000F | 4109843 | 4112081 | BUSCO |
| maker-000000F-augustus-gene-41.22-mRNA-1 | 2 | 000000F | 4110782 | 4111529 | BUSCO |
| genemark-000000F-processed-gene-42.36-mRNA-1 | 2 | 000000F | 4220355 | 4222962 | BUSCO |
| maker-000000F-augustus-gene-43.92-mRNA-1 | 2 | 000000F | 4383942 | 4387312 | BUSCO |
| augustus_masked-000000F-processed-gene-44.74-mRNA-1 | 2 | 000000F | 4485801 | 4491215 | BUSCO |
| maker-000000F-augustus-gene-46.57-mRNA-2 | 2 | 000000F | 4597449 | 4599199 | BUSCO |
| maker-000000F-augustus-gene-46.54-mRNA-1 | 2 | 000000F | 4649061 | 4650771 | BUSCO |
| maker-000000F-augustus-gene-47.65-mRNA-1 | 2 | 000000F | 4729763 | 4733849 | BUSCO |
| maker-000000F-augustus-gene-47.60-mRNA-1 | 2 | 000000F | 4745791 | 4753678 | BUSCO |
| maker-000000F-augustus-gene-48.89-mRNA-1 | 2 | 000000F | 4837377 | 4839825 | BUSCO |
| genemark-000000F-processed-gene-48.66-mRNA-1 | 2 | 000000F | 4852694 | 4854020 | BUSCO |
| maker-000000F-augustus-gene-48.79-mRNA-1 | 2 | 000000F | 4854017 | 4855872 | BUSCO |
| augustus_masked-000000F-processed-gene-49.19-mRNA-1 | 2 | 000000F | 4888821 | 4895895 | BUSCO |
| maker-000000F-augustus-gene-49.63-mRNA-1 | 2 | 000000F | 4932209 | 4936985 | BUSCO |
| genemark-000000F-processed-gene-51.40-mRNA-1 | 2 | 000000F | 5170594 | 5178589 | BUSCO |
| maker-000000F-augustus-gene-52.49-mRNA-1 | 2 | 000000F | 5242958 | 5250626 | BUSCO |
| maker-000000F-augustus-gene-55.71-mRNA-1 | 2 | 000000F | 5507279 | 5509285 | BUSCO |
| maker-000000F-augustus-gene-56.68-mRNA-1 | 2 | 000000F | 5622882 | 5623654 | BUSCO |
| genemark-000000F-processed-gene-56.18-mRNA-1 | 2 | 000000F | 5658726 | 5663806 | BUSCO |
| maker-000000F-augustus-gene-58.77-mRNA-1 | 2 | 000000F | 5791217 | 5793910 | BUSCO |
| augustus_masked-000000F-processed-gene-59.76-mRNA-1 | 2 | 000000F | 5893360 | 5898928 | BUSCO |
| maker-000000F-augustus-gene-59.6-mRNA-2 | 2 | 000000F | 5963575 | 5965024 | BUSCO |
| maker-000000F-augustus-gene-59.21-mRNA-1 | 2 | 000000F | 5965009 | 5966660 | BUSCO |
| genemark-000000F-processed-gene-62.7-mRNA-1 | 2 | 000000F | 6237157 | 6250934 | BUSCO |
| snap_masked-000000F-processed-gene-65.13-mRNA-1 | 2 | 000000F | 6545094 | 6546610 | BUSCO |
| maker-000000F-augustus-gene-66.48-mRNA-1 | 2 | 000000F | 6634531 | 6635428 | BUSCO |
| maker-000000F-augustus-gene-66.42-mRNA-1 | 2 | 000000F | 6635684 | 6637194 | BUSCO |
| maker-000000F-snap-gene-68.6-mRNA-1 | 2 | 000000F | 6820240 | 6824789 | BUSCO |
| maker-000000F-augustus-gene-68.87-mRNA-1 | 2 | 000000F | 6870669 | 6873063 | BUSCO |
| maker-000000F-snap-gene-71.70-mRNA-1 | 2 | 000000F | 7115261 | 7121168 | BUSCO |
| snap_masked-000000F-processed-gene-71.10-mRNA-1 | 2 | 000000F | 7139357 | 7144980 | BUSCO |
| maker-000000F-augustus-gene-72.68-mRNA-1 | 2 | 000000F | 7227407 | 7229091 | BUSCO |
| maker-000000F-augustus-gene-74.42-mRNA-1 | 2 | 000000F | 7392590 | 7400098 | BUSCO |
| augustus_masked-000000F-processed-gene-76.29-mRNA-1 | 2 | 000000F | 7615779 | 7619720 | BUSCO |
| genemark-000000F-processed-gene-78.14-mRNA-1 | 2 | 000000F | 7866912 | 7878370 | BUSCO |
| maker-000000F-augustus-gene-79.28-mRNA-1 | 2 | 000000F | 7915042 | 7917819 | BUSCO |
| maker-000000F-augustus-gene-79.30-mRNA-1 | 2 | 000000F | 7922979 | 7924509 | BUSCO |
| maker-000000F-augustus-gene-80.34-mRNA-3 | 2 | 000000F | 8063120 | 8066726 | BUSCO |
| maker-000000F-augustus-gene-81.46-mRNA-2 | 2 | 000000F | 8113803 | 8119007 | BUSCO |
| augustus_masked-000000F-processed-gene-85.67-mRNA-1 | 2 | 000000F | 8493966 | 8496359 | BUSCO |
| maker-000004F-snap-gene-0.84-mRNA-1 | 2 | 000004F | 92762 | 95284 | BUSCO |
| maker-000004F-snap-gene-2.82-mRNA-4 | 2 | 000004F | 195427 | 198907 | BUSCO |
| maker-000004F-snap-gene-2.82-mRNA-6 | 2 | 000004F | 200232 | 202237 | BUSCO |
| genemark-000004F-processed-gene-3.30-mRNA-1 | 2 | 000004F | 384737 | 390721 | BUSCO |
| maker-000004F-augustus-gene-5.68-mRNA-1 | 2 | 000004F | 545949 | 548856 | BUSCO |
| maker-000004F-augustus-gene-6.7-mRNA-1 | 2 | 000004F | 635892 | 636889 | BUSCO |
| maker-000004F-augustus-gene-7.83-mRNA-1 | 2 | 000004F | 741523 | 749396 | BUSCO |
| maker-000004F-augustus-gene-7.74-mRNA-1 | 2 | 000004F | 770144 | 773025 | BUSCO |
| genemark-000004F-processed-gene-8.53-mRNA-1 | 2 | 000004F | 805974 | 816374 | BUSCO |
| maker-000004F-augustus-gene-8.45-mRNA-1 | 2 | 000004F | 814838 | 816384 | BUSCO |
| genemark-000004F-processed-gene-15.50-mRNA-1 | 2 | 000004F | 1518376 | 1518921 | BUSCO |
| maker-000004F-augustus-gene-17.98-mRNA-1 | 2 | 000004F | 1703499 | 1705765 | BUSCO |
| maker-000004F-augustus-gene-18.49-mRNA-1 | 2 | 000004F | 1840644 | 1844045 | BUSCO |
| maker-000004F-augustus-gene-19.66-mRNA-2 | 2 | 000004F | 1924022 | 1926486 | BUSCO |
| maker-000004F-augustus-gene-19.67-mRNA-1 | 2 | 000004F | 1928231 | 1931365 | BUSCO |
| maker-000004F-augustus-gene-19.69-mRNA-1 | 2 | 000004F | 1945044 | 1947480 | BUSCO |
| maker-000004F-augustus-gene-19.70-mRNA-1 | 2 | 000004F | 1957008 | 1958290 | BUSCO |
| maker-000004F-augustus-gene-21.39-mRNA-1 | 2 | 000004F | 2176815 | 2184503 | BUSCO |
| maker-000004F-augustus-gene-23.101-mRNA-2 | 2 | 000004F | 2384968 | 2386376 | BUSCO |
| maker-000004F-augustus-gene-25.70-mRNA-1 | 2 | 000004F | 2496336 | 2501370 | BUSCO |
| maker-000004F-augustus-gene-25.61-mRNA-1 | 2 | 000004F | 2508086 | 2509794 | BUSCO |
| maker-000004F-augustus-gene-25.66-mRNA-3 | 2 | 000004F | 2552756 | 2560446 | BUSCO |
| maker-000004F-snap-gene-25.29-mRNA-2 | 2 | 000004F | 2561240 | 2570484 | BUSCO |
| maker-000004F-augustus-gene-26.100-mRNA-2 | 2 | 000004F | 2622864 | 2625240 | BUSCO |
| maker-000004F-snap-gene-29.67-mRNA-1 | 2 | 000004F | 2918797 | 2919631 | BUSCO |
| maker-000004F-augustus-gene-29.42-mRNA-1 | 2 | 000004F | 2933311 | 2934953 | BUSCO |
| maker-000004F-augustus-gene-31.47-mRNA-1 | 2 | 000004F | 3102982 | 3104817 | BUSCO |
| snap_masked-000004F-processed-gene-31.66-mRNA-1 | 2 | 000004F | 3104869 | 3105519 | BUSCO |
| maker-000004F-augustus-gene-31.48-mRNA-1 | 2 | 000004F | 3111694 | 3114604 | BUSCO |
| maker-000004F-snap-gene-35.43-mRNA-1 | 2 | 000004F | 3527511 | 3539189 | BUSCO |
| augustus_masked-000004F-processed-gene-35.18-mRNA-1 | 2 | 000004F | 3576039 | 3577088 | BUSCO |
| augustus_masked-000004F-processed-gene-36.23-mRNA-1 | 2 | 000004F | 3610408 | 3616314 | BUSCO |
| maker-000004F-snap-gene-37.118-mRNA-3 | 2 | 000004F | 3699129 | 3708794 | BUSCO |
| maker-000004F-augustus-gene-37.91-mRNA-1 | 2 | 000004F | 3747856 | 3749363 | BUSCO |
| maker-000004F-augustus-gene-38.33-mRNA-1 | 2 | 000004F | 3866254 | 3868137 | BUSCO |
| maker-000004F-augustus-gene-41.55-mRNA-2 | 2 | 000004F | 4183380 | 4184718 | BUSCO |
| maker-000007F-augustus-gene-9.80-mRNA-1 | 2 | 000007F | 955547 | 959573 | BUSCO |
| maker-000007F-augustus-gene-19.81-mRNA-1 | 2 | 000007F | 1955373 | 1957515 | BUSCO |
| snap_masked-000007F-processed-gene-20.69-mRNA-1 | 2 | 000007F | 2030168 | 2035024 | BUSCO |
| maker-000007F-augustus-gene-30.47-mRNA-2 | 2 | 000007F | 3002495 | 3007058 | BUSCO |
| maker-000036F-augustus-gene-0.17-mRNA-1 | 2 | 000036F | 55379 | 58394 | BUSCO |
| genemark-000036F-processed-gene-1.43-mRNA-1 | 2 | 000036F | 98338 | 99748 | BUSCO |
| maker-000036F-snap-gene-7.81-mRNA-3 | 2 | 000036F | 759628 | 763589 | BUSCO |
| maker-000036F-augustus-gene-8.100-mRNA-1 | 2 | 000036F | 845272 | 847487 | BUSCO |
| maker-000036F-snap-gene-8.109-mRNA-1 | 2 | 000036F | 850882 | 852688 | BUSCO |
| maker-000036F-augustus-gene-8.101-mRNA-1 | 2 | 000036F | 865132 | 866448 | BUSCO |
| maker-000036F-augustus-gene-9.42-mRNA-1 | 2 | 000036F | 968272 | 969852 | BUSCO |
| genemark-000036F-processed-gene-9.76-mRNA-1 | 2 | 000036F | 969794 | 979673 | BUSCO |
| maker-000045F-augustus-gene-3.45-mRNA-4 | 2 | 000045F | 292390 | 296621 | BUSCO |
| maker-000045F-snap-gene-4.48-mRNA-1 | 2 | 000045F | 387791 | 396877 | BUSCO |
| maker-000045F-augustus-gene-5.34-mRNA-1 | 2 | 000045F | 528109 | 532401 | BUSCO |
| augustus_masked-000002F-processed-gene-1.21-mRNA-1 | 3 | 000002F | 145759 | 149872 | BUSCO |
| genemark-000002F-processed-gene-3.25-mRNA-1 | 3 | 000002F | 306835 | 310560 | BUSCO |
| maker-000002F-augustus-gene-4.84-mRNA-1 | 3 | 000002F | 437486 | 440236 | BUSCO |
| maker-000002F-augustus-gene-7.17-mRNA-1 | 3 | 000002F | 785446 | 786359 | BUSCO |
| maker-000002F-augustus-gene-8.21-mRNA-1 | 3 | 000002F | 830542 | 842458 | BUSCO |
| maker-000002F-augustus-gene-9.60-mRNA-1 | 3 | 000002F | 894013 | 901754 | BUSCO |
| maker-000002F-augustus-gene-9.61-mRNA-2 | 3 | 000002F | 921099 | 927053 | BUSCO |
| genemark-000002F-processed-gene-10.2-mRNA-1 | 3 | 000002F | 1009536 | 1010341 | BUSCO |
| maker-000002F-augustus-gene-11.65-mRNA-1 | 3 | 000002F | 1076526 | 1079514 | BUSCO |
| maker-000002F-augustus-gene-11.75-mRNA-1 | 3 | 000002F | 1182527 | 1186973 | BUSCO |
| maker-000002F-snap-gene-14.35-mRNA-2 | 3 | 000002F | 1485314 | 1487723 | BUSCO |
| genemark-000002F-processed-gene-15.22-mRNA-1 | 3 | 000002F | 1569677 | 1570554 | BUSCO |
| genemark-000002F-processed-gene-18.9-mRNA-1 | 3 | 000002F | 1769660 | 1778685 | BUSCO |
| maker-000002F-augustus-gene-21.38-mRNA-1 | 3 | 000002F | 2111770 | 2112869 | BUSCO |
| maker-000002F-augustus-gene-23.102-mRNA-1 | 3 | 000002F | 2317171 | 2319859 | BUSCO |
| augustus_masked-000002F-processed-gene-24.12-mRNA-1 | 3 | 000002F | 2478150 | 2479798 | BUSCO |
| maker-000002F-augustus-gene-25.33-mRNA-3 | 3 | 000002F | 2511133 | 2514669 | BUSCO |
| maker-000002F-snap-gene-25.88-mRNA-1 | 3 | 000002F | 2518311 | 2522940 | BUSCO |
| maker-000002F-augustus-gene-29.41-mRNA-1 | 3 | 000002F | 2889326 | 2897326 | BUSCO |
| maker-000002F-augustus-gene-31.109-mRNA-3 | 3 | 000002F | 3114847 | 3118170 | BUSCO |
| maker-000002F-augustus-gene-31.96-mRNA-1 | 3 | 000002F | 3118695 | 3122367 | BUSCO |
| maker-000002F-augustus-gene-31.111-mRNA-1 | 3 | 000002F | 3132782 | 3133945 | BUSCO |
| maker-000002F-snap-gene-33.84-mRNA-1 | 3 | 000002F | 3361616 | 3363477 | BUSCO |
| maker-000002F-augustus-gene-34.33-mRNA-2 | 3 | 000002F | 3454704 | 3460881 | BUSCO |
| maker-000002F-snap-gene-36.23-mRNA-1 | 3 | 000002F | 3627690 | 3629542 | BUSCO |
| maker-000002F-augustus-gene-36.63-mRNA-1 | 3 | 000002F | 3630978 | 3640156 | BUSCO |
| maker-000009F-augustus-gene-0.120-mRNA-1 | 3 | 000009F | 69978 | 71420 | BUSCO |
| augustus_masked-000009F-processed-gene-4.4-mRNA-1 | 3 | 000009F | 380337 | 399071 | BUSCO |
| maker-000009F-augustus-gene-4.91-mRNA-1 | 3 | 000009F | 466952 | 468629 | BUSCO |
| maker-000009F-snap-gene-5.0-mRNA-1 | 3 | 000009F | 478227 | 481328 | BUSCO |
| maker-000009F-augustus-gene-10.31-mRNA-1 | 3 | 000009F | 1023203 | 1024938 | BUSCO |
| augustus_masked-000009F-processed-gene-10.65-mRNA-1 | 3 | 000009F | 1069706 | 1073905 | BUSCO |
| genemark-000009F-processed-gene-11.49-mRNA-1 | 3 | 000009F | 1162764 | 1163210 | BUSCO |
| maker-000009F-augustus-gene-11.11-mRNA-1 | 3 | 000009F | 1163314 | 1164645 | BUSCO |
| maker-000009F-augustus-gene-14.52-mRNA-1 | 3 | 000009F | 1440776 | 1442862 | BUSCO |
| maker-000009F-augustus-gene-16.36-mRNA-1 | 3 | 000009F | 1592931 | 1594709 | BUSCO |
| maker-000009F-snap-gene-17.88-mRNA-1 | 3 | 000009F | 1735856 | 1745434 | BUSCO |
| genemark-000009F-processed-gene-17.63-mRNA-1 | 3 | 000009F | 1745440 | 1752301 | BUSCO |
| maker-000009F-augustus-gene-17.125-mRNA-1 | 3 | 000009F | 1770634 | 1774257 | BUSCO |
| maker-000009F-snap-gene-17.96-mRNA-2 | 3 | 000009F | 1779087 | 1784828 | BUSCO |
| maker-000009F-augustus-gene-19.111-mRNA-1 | 3 | 000009F | 1968251 | 1969838 | BUSCO |
| maker-000009F-augustus-gene-20.62-mRNA-1 | 3 | 000009F | 2027258 | 2029029 | BUSCO |
| maker-000009F-augustus-gene-21.88-mRNA-1 | 3 | 000009F | 2096783 | 2099278 | BUSCO |
| augustus_masked-000009F-processed-gene-21.75-mRNA-1 | 3 | 000009F | 2142325 | 2153813 | BUSCO |
| maker-000009F-augustus-gene-21.103-mRNA-2 | 3 | 000009F | 2170961 | 2172771 | BUSCO |
| maker-000009F-augustus-gene-21.96-mRNA-1 | 3 | 000009F | 2184182 | 2188836 | BUSCO |
| maker-000009F-snap-gene-23.43-mRNA-1 | 3 | 000009F | 2367296 | 2374141 | BUSCO |
| genemark-000009F-processed-gene-23.13-mRNA-1 | 3 | 000009F | 2375122 | 2377524 | BUSCO |
| maker-000009F-augustus-gene-23.59-mRNA-1 | 3 | 000009F | 2377640 | 2383106 | BUSCO |
| maker-000009F-augustus-gene-24.62-mRNA-2 | 3 | 000009F | 2480878 | 2484009 | BUSCO |
| maker-000009F-snap-gene-25.42-mRNA-2 | 3 | 000009F | 2505100 | 2511658 | BUSCO |
| maker-000009F-augustus-gene-25.73-mRNA-2 | 3 | 000009F | 2526179 | 2529535 | BUSCO |
| snap_masked-000009F-processed-gene-29.2-mRNA-1 | 3 | 000009F | 2886829 | 2895057 | BUSCO |
| maker-000009F-augustus-gene-29.65-mRNA-2 | 3 | 000009F | 2897744 | 2898551 | BUSCO |
| maker-000017F-augustus-gene-8.20-mRNA-2 | 3 | 000017F | 879381 | 883295 | BUSCO |
| maker-000017F-augustus-gene-11.88-mRNA-1 | 3 | 000017F | 1111865 | 1114758 | BUSCO |
| maker-000017F-augustus-gene-11.90-mRNA-4 | 3 | 000017F | 1139037 | 1140875 | BUSCO |
| maker-000017F-augustus-gene-16.100-mRNA-1 | 3 | 000017F | 1669187 | 1673369 | BUSCO |
| maker-000017F-augustus-gene-17.138-mRNA-1 | 3 | 000017F | 1752314 | 1754140 | BUSCO |
| maker-000017F-augustus-gene-19.61-mRNA-1 | 3 | 000017F | 1919046 | 1924183 | BUSCO |
| maker-000017F-augustus-gene-22.35-mRNA-1 | 3 | 000017F | 2226643 | 2228041 | BUSCO |
| maker-000024F-augustus-gene-0.97-mRNA-1 | 3 | 000024F | 89127 | 90480 | BUSCO |
| maker-000024F-augustus-gene-4.84-mRNA-1 | 3 | 000024F | 459989 | 461993 | BUSCO |
| maker-000024F-augustus-gene-4.92-mRNA-3 | 3 | 000024F | 463288 | 471151 | BUSCO |
| maker-000024F-augustus-gene-14.120-mRNA-2 | 3 | 000024F | 1470582 | 1472320 | BUSCO |
| augustus_masked-000024F-processed-gene-14.64-mRNA-1 | 3 | 000024F | 1492626 | 1495696 | BUSCO |
| maker-000024F-augustus-gene-16.48-mRNA-2 | 3 | 000024F | 1624373 | 1626842 | BUSCO |
| maker-000024F-augustus-gene-16.47-mRNA-5 | 3 | 000024F | 1686395 | 1688238 | BUSCO |
| maker-000024F-augustus-gene-18.51-mRNA-1 | 3 | 000024F | 1797794 | 1800538 | BUSCO |
| maker-000024F-augustus-gene-18.53-mRNA-1 | 3 | 000024F | 1818290 | 1819204 | BUSCO |
| maker-000024F-augustus-gene-20.54-mRNA-1 | 3 | 000024F | 1988293 | 1989593 | BUSCO |
| maker-000024F-augustus-gene-19.40-mRNA-1 | 3 | 000024F | 1989479 | 1990714 | BUSCO |
| maker-000024F-augustus-gene-20.55-mRNA-2 | 3 | 000024F | 1992065 | 1996435 | BUSCO |
| maker-000044F-augustus-gene-2.59-mRNA-1 | 3 | 000044F | 223803 | 231959 | BUSCO |
| maker-000044F-augustus-gene-5.8-mRNA-1 | 3 | 000044F | 499568 | 502747 | BUSCO |
| maker-000046F-augustus-gene-1.56-mRNA-1 | 3 | 000046F | 97339 | 99771 | BUSCO |
| maker-000046F-augustus-gene-1.49-mRNA-1 | 3 | 000046F | 99970 | 103105 | BUSCO |
| genemark-000046F-processed-gene-3.21-mRNA-1 | 3 | 000046F | 311868 | 315080 | BUSCO |
| maker-000046F-augustus-gene-3.34-mRNA-2 | 3 | 000046F | 380769 | 382952 | BUSCO |
| maker-000046F-augustus-gene-4.86-mRNA-2 | 3 | 000046F | 463121 | 464785 | BUSCO |
| maker-000046F-augustus-gene-5.13-mRNA-1 | 3 | 000046F | 497948 | 500637 | BUSCO |
| maker-000046F-augustus-gene-6.142-mRNA-1 | 3 | 000046F | 718801 | 719877 | BUSCO |
| augustus_masked-000046F-processed-gene-6.126-mRNA-1 | 3 | 000046F | 749161 | 751205 | BUSCO |
| maker-000062F-augustus-gene-1.44-mRNA-1 | 3 | 000062F | 124597 | 126015 | BUSCO |
| maker-000062F-augustus-gene-1.57-mRNA-2 | 3 | 000062F | 185487 | 186906 | BUSCO |
| augustus_masked-000062F-processed-gene-2.19-mRNA-1 | 3 | 000062F | 234622 | 237228 | BUSCO |
| maker-000062F-augustus-gene-3.80-mRNA-1 | 3 | 000062F | 397965 | 402332 | BUSCO |
| augustus_masked-000068F-processed-gene-0.4-mRNA-1 | 3 | 000068F | 41102 | 54864 | BUSCO |
| maker-000068F-augustus-gene-0.36-mRNA-1 | 3 | 000068F | 70349 | 83495 | BUSCO |
| maker-000115F-augustus-gene-0.68-mRNA-1 | 3 | 000115F | 51025 | 52232 | BUSCO |
| maker-000115F-augustus-gene-0.77-mRNA-1 | 3 | 000115F | 53892 | 58487 | BUSCO |
| snap_masked-000011F-processed-gene-0.89-mRNA-1 | 4 | 000011F | 30623 | 31168 | BUSCO |
| maker-000011F-augustus-gene-1.112-mRNA-1 | 4 | 000011F | 115296 | 120174 | BUSCO |
| maker-000011F-augustus-gene-1.116-mRNA-2 | 4 | 000011F | 141777 | 144281 | BUSCO |
| maker-000011F-augustus-gene-1.101-mRNA-3 | 4 | 000011F | 147372 | 149074 | BUSCO |
| maker-000011F-augustus-gene-1.101-mRNA-1 | 4 | 000011F | 147375 | 152661 | BUSCO |
| snap_masked-000011F-processed-gene-4.56-mRNA-1 | 4 | 000011F | 419778 | 421979 | BUSCO |
| genemark-000011F-processed-gene-4.24-mRNA-1 | 4 | 000011F | 471198 | 474297 | BUSCO |
| genemark-000011F-processed-gene-6.15-mRNA-1 | 4 | 000011F | 647064 | 649145 | BUSCO |
| maker-000011F-snap-gene-7.111-mRNA-1 | 4 | 000011F | 710946 | 712522 | BUSCO |
| maker-000011F-augustus-gene-10.53-mRNA-1 | 4 | 000011F | 1017672 | 1019684 | BUSCO |
| maker-000011F-augustus-gene-12.77-mRNA-1 | 4 | 000011F | 1290371 | 1291170 | BUSCO |
| maker-000011F-snap-gene-13.65-mRNA-1 | 4 | 000011F | 1321746 | 1323274 | BUSCO |
| maker-000011F-augustus-gene-14.27-mRNA-1 | 4 | 000011F | 1448852 | 1450565 | BUSCO |
| maker-000011F-augustus-gene-15.83-mRNA-1 | 4 | 000011F | 1502640 | 1505250 | BUSCO |
| snap_masked-000011F-processed-gene-15.3-mRNA-1 | 4 | 000011F | 1506259 | 1509526 | BUSCO |
| maker-000011F-augustus-gene-15.86-mRNA-1 | 4 | 000011F | 1521782 | 1523238 | BUSCO |
| maker-000011F-augustus-gene-16.27-mRNA-2 | 4 | 000011F | 1658832 | 1660834 | BUSCO |
| augustus_masked-000011F-processed-gene-17.37-mRNA-1 | 4 | 000011F | 1748412 | 1749495 | BUSCO |
| maker-000011F-augustus-gene-18.0-mRNA-2 | 4 | 000011F | 1752601 | 1754597 | BUSCO |
| genemark-000011F-processed-gene-18.58-mRNA-1 | 4 | 000011F | 1874058 | 1876628 | BUSCO |
| genemark-000011F-processed-gene-22.30-mRNA-1 | 4 | 000011F | 2190255 | 2202678 | BUSCO |
| maker-000011F-augustus-gene-22.101-mRNA-1 | 4 | 000011F | 2220178 | 2221468 | BUSCO |
| genemark-000011F-processed-gene-22.32-mRNA-1 | 4 | 000011F | 2220225 | 2235233 | BUSCO |
| maker-000011F-augustus-gene-24.53-mRNA-2 | 4 | 000011F | 2445705 | 2451222 | BUSCO |
| maker-000011F-augustus-gene-25.85-mRNA-2 | 4 | 000011F | 2500027 | 2502559 | BUSCO |
| maker-000011F-augustus-gene-28.156-mRNA-1 | 4 | 000011F | 2868012 | 2870259 | BUSCO |
| maker-000011F-augustus-gene-28.191-mRNA-2 | 4 | 000011F | 2905507 | 2908275 | BUSCO |
| maker-000011F-augustus-gene-28.194-mRNA-1 | 4 | 000011F | 2937442 | 2939911 | BUSCO |
| genemark-000023F-processed-gene-1.28-mRNA-1 | 4 | 000023F | 174030 | 185483 | BUSCO |
| maker-000023F-augustus-gene-2.12-mRNA-3 | 4 | 000023F | 283968 | 289121 | BUSCO |
| maker-000023F-augustus-gene-2.22-mRNA-1 | 4 | 000023F | 289250 | 291456 | BUSCO |
| maker-000023F-augustus-gene-4.99-mRNA-1 | 4 | 000023F | 473200 | 475768 | BUSCO |
| maker-000023F-augustus-gene-7.125-mRNA-1 | 4 | 000023F | 704085 | 705447 | BUSCO |
| maker-000023F-augustus-gene-10.74-mRNA-1 | 4 | 000023F | 1003107 | 1004053 | BUSCO |
| maker-000023F-augustus-gene-10.88-mRNA-1 | 4 | 000023F | 1034360 | 1036951 | BUSCO |
| maker-000023F-augustus-gene-10.76-mRNA-1 | 4 | 000023F | 1036783 | 1038644 | BUSCO |
| maker-000023F-augustus-gene-10.92-mRNA-1 | 4 | 000023F | 1087365 | 1091043 | BUSCO |
| maker-000023F-snap-gene-12.19-mRNA-2 | 4 | 000023F | 1194639 | 1199620 | BUSCO |
| genemark-000023F-processed-gene-13.2-mRNA-1 | 4 | 000023F | 1286577 | 1292364 | BUSCO |
| maker-000023F-augustus-gene-13.86-mRNA-1 | 4 | 000023F | 1315947 | 1320270 | BUSCO |
| genemark-000023F-processed-gene-13.25-mRNA-1 | 4 | 000023F | 1351906 | 1353679 | BUSCO |
| maker-000023F-snap-gene-15.86-mRNA-1 | 4 | 000023F | 1508198 | 1511383 | BUSCO |
| augustus_masked-000023F-processed-gene-15.48-mRNA-1 | 4 | 000023F | 1535467 | 1537568 | BUSCO |
| maker-000023F-augustus-gene-15.139-mRNA-1 | 4 | 000023F | 1540220 | 1543447 | BUSCO |
| maker-000023F-augustus-gene-18.62-mRNA-3 | 4 | 000023F | 1818291 | 1819667 | BUSCO |
| augustus_masked-000026F-processed-gene-0.42-mRNA-1 | 4 | 000026F | 17734 | 21144 | BUSCO |
| maker-000026F-augustus-gene-1.85-mRNA-3 | 4 | 000026F | 140110 | 143212 | BUSCO |
| maker-000026F-augustus-gene-2.81-mRNA-1 | 4 | 000026F | 213632 | 216057 | BUSCO |
| augustus_masked-000026F-processed-gene-4.80-mRNA-1 | 4 | 000026F | 415160 | 430931 | BUSCO |
| genemark-000026F-processed-gene-7.40-mRNA-1 | 4 | 000026F | 759784 | 764335 | BUSCO |
| maker-000026F-augustus-gene-13.14-mRNA-1 | 4 | 000026F | 1283765 | 1286429 | BUSCO |
| maker-000026F-augustus-gene-13.17-mRNA-1 | 4 | 000026F | 1293325 | 1295223 | BUSCO |
| maker-000026F-augustus-gene-13.11-mRNA-1 | 4 | 000026F | 1376277 | 1378729 | BUSCO |
| genemark-000026F-processed-gene-14.28-mRNA-1 | 4 | 000026F | 1438165 | 1441739 | BUSCO |
| maker-000026F-augustus-gene-15.47-mRNA-1 | 4 | 000026F | 1530319 | 1532445 | BUSCO |
| maker-000028F-augustus-gene-0.111-mRNA-1 | 4 | 000028F | 65981 | 68604 | BUSCO |
| maker-000028F-augustus-gene-5.34-mRNA-1 | 4 | 000028F | 513571 | 516268 | BUSCO |
| maker-000028F-augustus-gene-7.39-mRNA-1 | 4 | 000028F | 725507 | 727352 | BUSCO |
| augustus_masked-000028F-processed-gene-8.61-mRNA-1 | 4 | 000028F | 805669 | 809036 | BUSCO |
| maker-000028F-augustus-gene-8.85-mRNA-1 | 4 | 000028F | 809104 | 810393 | BUSCO |
| maker-000028F-augustus-gene-8.87-mRNA-2 | 4 | 000028F | 820520 | 827298 | BUSCO |
| maker-000028F-augustus-gene-12.103-mRNA-2 | 4 | 000028F | 1187155 | 1193541 | BUSCO |
| augustus_masked-000028F-processed-gene-13.42-mRNA-1 | 4 | 000028F | 1302325 | 1313013 | BUSCO |
| maker-000028F-snap-gene-14.8-mRNA-2 | 4 | 000028F | 1430132 | 1435129 | BUSCO |
| maker-000051F-augustus-gene-5.107-mRNA-1 | 4 | 000051F | 561087 | 565142 | BUSCO |
| maker-000060F-augustus-gene-2.70-mRNA-2 | 4 | 000060F | 272622 | 274200 | BUSCO |
| maker-000060F-augustus-gene-3.17-mRNA-2 | 4 | 000060F | 332670 | 335141 | BUSCO |
| maker-000064F-augustus-gene-0.43-mRNA-1 | 4 | 000064F | 29990 | 31455 | BUSCO |
| maker-000064F-augustus-gene-2.106-mRNA-2 | 4 | 000064F | 246227 | 249735 | BUSCO |
| maker-000065F-augustus-gene-2.51-mRNA-1 | 4 | 000065F | 186648 | 189069 | BUSCO |
| maker-000069F-augustus-gene-2.70-mRNA-1 | 4 | 000069F | 270548 | 279250 | BUSCO |
| maker-000069F-augustus-gene-3.39-mRNA-1 | 4 | 000069F | 320561 | 322981 | BUSCO |
| maker-000087F-augustus-gene-0.65-mRNA-1 | 4 | 000087F | 77990 | 81682 | BUSCO |
| maker-000087F-augustus-gene-1.72-mRNA-1 | 4 | 000087F | 133557 | 136455 | BUSCO |
| maker-000092F-augustus-gene-0.17-mRNA-1 | 4 | 000092F | 84918 | 86561 | BUSCO |
| maker-000225F-augustus-gene-0.20-mRNA-1 | 4 | 000225F | 22230 | 24099 | BUSCO |
| maker-000001F-augustus-gene-1.68-mRNA-1 | 5 | 000001F | 136739 | 138897 | BUSCO |
| maker-000001F-snap-gene-2.18-mRNA-1 | 5 | 000001F | 241047 | 242248 | BUSCO |
| maker-000001F-augustus-gene-2.80-mRNA-1 | 5 | 000001F | 282991 | 291013 | BUSCO |
| maker-000001F-snap-gene-3.67-mRNA-1 | 5 | 000001F | 320245 | 325129 | BUSCO |
| maker-000001F-augustus-gene-3.63-mRNA-1 | 5 | 000001F | 335152 | 337058 | BUSCO |
| maker-000001F-augustus-gene-4.33-mRNA-1 | 5 | 000001F | 439765 | 440714 | BUSCO |
| genemark-000001F-processed-gene-6.80-mRNA-1 | 5 | 000001F | 691381 | 694222 | BUSCO |
| maker-000001F-augustus-gene-8.27-mRNA-1 | 5 | 000001F | 819047 | 820886 | BUSCO |
| augustus_masked-000001F-processed-gene-10.23-mRNA-1 | 5 | 000001F | 1042193 | 1043511 | BUSCO |
| genemark-000001F-processed-gene-10.49-mRNA-1 | 5 | 000001F | 1043164 | 1046225 | BUSCO |
| maker-000001F-augustus-gene-11.14-mRNA-1 | 5 | 000001F | 1170339 | 1174209 | BUSCO |
| maker-000001F-augustus-gene-12.71-mRNA-3 | 5 | 000001F | 1262047 | 1275395 | BUSCO |
| maker-000001F-augustus-gene-12.63-mRNA-1 | 5 | 000001F | 1272083 | 1273684 | BUSCO |
| maker-000001F-augustus-gene-15.54-mRNA-2 | 5 | 000001F | 1498764 | 1500838 | BUSCO |
| maker-000001F-augustus-gene-15.58-mRNA-1 | 5 | 000001F | 1522289 | 1523109 | BUSCO |
| maker-000001F-augustus-gene-16.81-mRNA-1 | 5 | 000001F | 1594713 | 1600121 | BUSCO |
| maker-000001F-augustus-gene-16.86-mRNA-1 | 5 | 000001F | 1634348 | 1635615 | BUSCO |
| maker-000001F-snap-gene-16.65-mRNA-1 | 5 | 000001F | 1637681 | 1639584 | BUSCO |
| maker-000001F-augustus-gene-21.15-mRNA-1 | 5 | 000001F | 2107858 | 2115248 | BUSCO |
| augustus_masked-000001F-processed-gene-22.25-mRNA-1 | 5 | 000001F | 2254465 | 2255881 | BUSCO |
| maker-000001F-augustus-gene-24.26-mRNA-2 | 5 | 000001F | 2432279 | 2437870 | BUSCO |
| maker-000001F-augustus-gene-25.57-mRNA-2 | 5 | 000001F | 2568500 | 2571701 | BUSCO |
| genemark-000001F-processed-gene-25.39-mRNA-1 | 5 | 000001F | 2575514 | 2577976 | BUSCO |
| maker-000001F-augustus-gene-29.94-mRNA-1 | 5 | 000001F | 2935432 | 2937469 | BUSCO |
| augustus_masked-000001F-processed-gene-30.52-mRNA-1 | 5 | 000001F | 3010078 | 3012186 | BUSCO |
| maker-000001F-augustus-gene-30.7-mRNA-2 | 5 | 000001F | 3048215 | 3051441 | BUSCO |
| maker-000001F-augustus-gene-32.28-mRNA-1 | 5 | 000001F | 3225876 | 3228775 | BUSCO |
| maker-000001F-augustus-gene-33.43-mRNA-2 | 5 | 000001F | 3319977 | 3326131 | BUSCO |
| maker-000001F-snap-gene-34.21-mRNA-1 | 5 | 000001F | 3442598 | 3443860 | BUSCO |
| maker-000001F-augustus-gene-35.70-mRNA-1 | 5 | 000001F | 3546883 | 3548251 | BUSCO |
| maker-000001F-augustus-gene-36.77-mRNA-2 | 5 | 000001F | 3665430 | 3671402 | BUSCO |
| maker-000001F-augustus-gene-37.61-mRNA-2 | 5 | 000001F | 3701190 | 3703780 | BUSCO |
| snap_masked-000001F-processed-gene-37.73-mRNA-1 | 5 | 000001F | 3722876 | 3731433 | BUSCO |
| genemark-000001F-processed-gene-37.20-mRNA-1 | 5 | 000001F | 3750722 | 3756340 | BUSCO |
| maker-000001F-augustus-gene-37.56-mRNA-1 | 5 | 000001F | 3768656 | 3769550 | BUSCO |
| maker-000001F-snap-gene-38.14-mRNA-4 | 5 | 000001F | 3822100 | 3823866 | BUSCO |
| maker-000001F-augustus-gene-38.77-mRNA-2 | 5 | 000001F | 3859382 | 3861380 | BUSCO |
| maker-000001F-augustus-gene-39.76-mRNA-4 | 5 | 000001F | 3944865 | 3946974 | BUSCO |
| maker-000001F-augustus-gene-40.89-mRNA-1 | 5 | 000001F | 4034683 | 4037034 | BUSCO |
| maker-000001F-augustus-gene-40.91-mRNA-3 | 5 | 000001F | 4088112 | 4091045 | BUSCO |
| maker-000001F-augustus-gene-41.67-mRNA-1 | 5 | 000001F | 4107503 | 4109221 | BUSCO |
| maker-000001F-snap-gene-41.1-mRNA-1 | 5 | 000001F | 4109168 | 4110788 | BUSCO |
| maker-000001F-snap-gene-41.4-mRNA-5 | 5 | 000001F | 4124257 | 4126747 | BUSCO |
| maker-000001F-augustus-gene-42.64-mRNA-1 | 5 | 000001F | 4205116 | 4206620 | BUSCO |
| maker-000001F-augustus-gene-44.32-mRNA-1 | 5 | 000001F | 4409322 | 4410908 | BUSCO |
| augustus_masked-000001F-processed-gene-44.21-mRNA-1 | 5 | 000001F | 4411027 | 4413792 | BUSCO |
| maker-000001F-augustus-gene-48.63-mRNA-1 | 5 | 000001F | 4819173 | 4819938 | BUSCO |
| maker-000001F-augustus-gene-49.45-mRNA-2 | 5 | 000001F | 4910172 | 4918223 | BUSCO |
| maker-000001F-augustus-gene-49.51-mRNA-3 | 5 | 000001F | 4958820 | 4963753 | BUSCO |
| maker-000001F-augustus-gene-51.29-mRNA-1 | 5 | 000001F | 5101616 | 5108630 | BUSCO |
| maker-000001F-augustus-gene-53.3-mRNA-1 | 5 | 000001F | 5311821 | 5312987 | BUSCO |
| maker-000001F-augustus-gene-57.48-mRNA-1 | 5 | 000001F | 5745336 | 5746761 | BUSCO |
| maker-000001F-augustus-gene-59.29-mRNA-1 | 5 | 000001F | 5903524 | 5905356 | BUSCO |
| maker-000001F-augustus-gene-59.26-mRNA-1 | 5 | 000001F | 5952371 | 5952979 | BUSCO |
| maker-000001F-augustus-gene-59.33-mRNA-1 | 5 | 000001F | 5953154 | 5954596 | BUSCO |
| maker-000001F-augustus-gene-61.84-mRNA-1 | 5 | 000001F | 6189892 | 6191377 | BUSCO |
| maker-000001F-augustus-gene-64.34-mRNA-1 | 5 | 000001F | 6437031 | 6440320 | BUSCO |
| maker-000025F-snap-gene-14.45-mRNA-1 | 5 | 000025F | 1420595 | 1421477 | BUSCO |
| maker-000027F-augustus-gene-14.85-mRNA-1 | 5 | 000027F | 1469314 | 1472989 | BUSCO |
| augustus_masked-000049F-processed-gene-5.2-mRNA-1 | 5 | 000049F | 618297 | 621915 | BUSCO |
| maker-000049F-augustus-gene-5.163-mRNA-1 | 5 | 000049F | 620914 | 622137 | BUSCO |
| genemark-000049F-processed-gene-5.98-mRNA-1 | 5 | 000049F | 638008 | 649468 | BUSCO |
| maker-000061F-augustus-gene-2.19-mRNA-2 | 5 | 000061F | 198217 | 200813 | BUSCO |
| maker-000061F-augustus-gene-3.79-mRNA-1 | 5 | 000061F | 391616 | 393556 | BUSCO |
| maker-000067F-augustus-gene-1.108-mRNA-1 | 5 | 000067F | 162111 | 163764 | BUSCO |
| genemark-000067F-processed-gene-2.127-mRNA-1 | 5 | 000067F | 219391 | 222104 | BUSCO |
| maker-000067F-augustus-gene-3.127-mRNA-1 | 5 | 000067F | 363786 | 366101 | BUSCO |
| maker-000074F-augustus-gene-2.87-mRNA-1 | 5 | 000074F | 201922 | 203588 | BUSCO |
| maker-000077F-augustus-gene-0.27-mRNA-2 | 5 | 000077F | 34794 | 36071 | BUSCO |
| maker-000077F-augustus-gene-0.29-mRNA-1 | 5 | 000077F | 54469 | 57521 | BUSCO |
| snap_masked-000080F-processed-gene-1.46-mRNA-1 | 5 | 000080F | 231100 | 232573 | BUSCO |
| augustus_masked-000108F-processed-gene-0.95-mRNA-1 | 5 | 000108F | 111290 | 113217 | BUSCO |
| snap_masked-000012F-processed-gene-10.67-mRNA-1 | 6 | 000012F | 1004242 | 1006152 | BUSCO |
| augustus_masked-000012F-processed-gene-10.36-mRNA-1 | 6 | 000012F | 1033474 | 1034232 | BUSCO |
| maker-000012F-snap-gene-11.68-mRNA-2 | 6 | 000012F | 1099590 | 1109467 | BUSCO |
| maker-000012F-augustus-gene-12.37-mRNA-4 | 6 | 000012F | 1231879 | 1235241 | BUSCO |
| maker-000012F-augustus-gene-13.7-mRNA-1 | 6 | 000012F | 1354656 | 1355812 | BUSCO |
| maker-000012F-augustus-gene-13.29-mRNA-1 | 6 | 000012F | 1382758 | 1385480 | BUSCO |
| maker-000012F-snap-gene-13.96-mRNA-1 | 6 | 000012F | 1391731 | 1393433 | BUSCO |
| maker-000012F-augustus-gene-15.29-mRNA-1 | 6 | 000012F | 1562525 | 1567074 | BUSCO |
| maker-000012F-augustus-gene-15.35-mRNA-1 | 6 | 000012F | 1567217 | 1569956 | BUSCO |
| maker-000012F-augustus-gene-16.143-mRNA-1 | 6 | 000012F | 1592933 | 1596345 | BUSCO |
| maker-000012F-augustus-gene-16.132-mRNA-1 | 6 | 000012F | 1596209 | 1597550 | BUSCO |
| maker-000012F-augustus-gene-16.145-mRNA-1 | 6 | 000012F | 1603342 | 1604654 | BUSCO |
| maker-000012F-augustus-gene-18.71-mRNA-2 | 6 | 000012F | 1804064 | 1807180 | BUSCO |
| augustus_masked-000012F-processed-gene-20.69-mRNA-1 | 6 | 000012F | 1977742 | 1994695 | BUSCO |
| maker-000012F-augustus-gene-20.65-mRNA-1 | 6 | 000012F | 2025377 | 2026836 | BUSCO |
| maker-000012F-augustus-gene-20.59-mRNA-1 | 6 | 000012F | 2030881 | 2033408 | BUSCO |
| maker-000012F-augustus-gene-21.77-mRNA-1 | 6 | 000012F | 2151341 | 2153998 | BUSCO |
| maker-000012F-augustus-gene-22.66-mRNA-1 | 6 | 000012F | 2251176 | 2261105 | BUSCO |
| maker-000012F-augustus-gene-22.75-mRNA-1 | 6 | 000012F | 2285529 | 2288072 | BUSCO |
| maker-000012F-augustus-gene-22.68-mRNA-1 | 6 | 000012F | 2290467 | 2291772 | BUSCO |
| maker-000012F-augustus-gene-23.48-mRNA-1 | 6 | 000012F | 2291725 | 2294324 | BUSCO |
| genemark-000012F-processed-gene-24.7-mRNA-1 | 6 | 000012F | 2477408 | 2479396 | BUSCO |
| maker-000012F-augustus-gene-25.87-mRNA-1 | 6 | 000012F | 2543300 | 2545692 | BUSCO |
| genemark-000012F-processed-gene-26.58-mRNA-1 | 6 | 000012F | 2674373 | 2675660 | BUSCO |
| maker-000012F-augustus-gene-27.85-mRNA-1 | 6 | 000012F | 2743501 | 2746153 | BUSCO |
| maker-000019F-augustus-gene-3.65-mRNA-2 | 6 | 000019F | 343356 | 348069 | BUSCO |
| maker-000019F-augustus-gene-6.10-mRNA-1 | 6 | 000019F | 659027 | 661786 | BUSCO |
| maker-000019F-augustus-gene-7.71-mRNA-2 | 6 | 000019F | 699864 | 701778 | BUSCO |
| augustus_masked-000019F-processed-gene-13.90-mRNA-1 | 6 | 000019F | 1371169 | 1378630 | BUSCO |
| maker-000019F-augustus-gene-15.61-mRNA-1 | 6 | 000019F | 1512869 | 1516315 | BUSCO |
| maker-000019F-augustus-gene-15.62-mRNA-1 | 6 | 000019F | 1529069 | 1532190 | BUSCO |
| augustus_masked-000019F-processed-gene-20.30-mRNA-1 | 6 | 000019F | 2009714 | 2016863 | BUSCO |
| maker-000019F-snap-gene-21.50-mRNA-3 | 6 | 000019F | 2116186 | 2117254 | BUSCO |
| maker-000019F-augustus-gene-21.39-mRNA-1 | 6 | 000019F | 2130744 | 2133830 | BUSCO |
| maker-000019F-augustus-gene-23.68-mRNA-1 | 6 | 000019F | 2338656 | 2340557 | BUSCO |
| maker-000019F-augustus-gene-23.81-mRNA-1 | 6 | 000019F | 2394487 | 2395591 | BUSCO |
| maker-000019F-augustus-gene-24.98-mRNA-1 | 6 | 000019F | 2402238 | 2403876 | BUSCO |
| maker-000021F-augustus-gene-1.74-mRNA-1 | 6 | 000021F | 106124 | 110343 | BUSCO |
| maker-000021F-augustus-gene-4.125-mRNA-4 | 6 | 000021F | 405491 | 415904 | BUSCO |
| maker-000021F-augustus-gene-5.50-mRNA-1 | 6 | 000021F | 512841 | 515880 | BUSCO |
| maker-000021F-augustus-gene-6.67-mRNA-1 | 6 | 000021F | 593482 | 599836 | BUSCO |
| augustus_masked-000021F-processed-gene-6.53-mRNA-1 | 6 | 000021F | 615633 | 615842 | BUSCO |
| maker-000021F-augustus-gene-7.41-mRNA-3 | 6 | 000021F | 711354 | 713295 | BUSCO |
| genemark-000021F-processed-gene-7.23-mRNA-1 | 6 | 000021F | 772825 | 783932 | BUSCO |
| maker-000021F-augustus-gene-8.54-mRNA-1 | 6 | 000021F | 783361 | 785549 | BUSCO |
| maker-000021F-augustus-gene-7.40-mRNA-1 | 6 | 000021F | 790458 | 793138 | BUSCO |
| genemark-000021F-processed-gene-9.29-mRNA-1 | 6 | 000021F | 986741 | 990574 | BUSCO |
| maker-000021F-augustus-gene-10.31-mRNA-1 | 6 | 000021F | 995611 | 996986 | BUSCO |
| maker-000021F-augustus-gene-11.9-mRNA-1 | 6 | 000021F | 1110772 | 1116652 | BUSCO |
| genemark-000021F-processed-gene-11.46-mRNA-1 | 6 | 000021F | 1144721 | 1147888 | BUSCO |
| maker-000021F-snap-gene-11.58-mRNA-3 | 6 | 000021F | 1186131 | 1188658 | BUSCO |
| maker-000021F-augustus-gene-12.78-mRNA-1 | 6 | 000021F | 1196259 | 1198067 | BUSCO |
| maker-000021F-augustus-gene-13.41-mRNA-1 | 6 | 000021F | 1292599 | 1295430 | BUSCO |
| maker-000021F-augustus-gene-16.12-mRNA-1 | 6 | 000021F | 1644439 | 1646712 | BUSCO |
| maker-000021F-augustus-gene-17.87-mRNA-3 | 6 | 000021F | 1746071 | 1750992 | BUSCO |
| maker-000021F-augustus-gene-18.25-mRNA-1 | 6 | 000021F | 1796471 | 1797939 | BUSCO |
| maker-000021F-augustus-gene-19.61-mRNA-1 | 6 | 000021F | 1871858 | 1873548 | BUSCO |
| maker-000021F-augustus-gene-19.51-mRNA-1 | 6 | 000021F | 1874384 | 1876737 | BUSCO |
| genemark-000021F-processed-gene-19.81-mRNA-1 | 6 | 000021F | 1876815 | 1878812 | BUSCO |
| maker-000021F-augustus-gene-20.134-mRNA-1 | 6 | 000021F | 1979184 | 1980729 | BUSCO |
| genemark-000021F-processed-gene-20.86-mRNA-1 | 6 | 000021F | 1980921 | 1983660 | BUSCO |
| maker-000052F-augustus-gene-0.116-mRNA-1 | 6 | 000052F | 55499 | 57658 | BUSCO |
| maker-000052F-augustus-gene-3.47-mRNA-1 | 6 | 000052F | 362690 | 364485 | BUSCO |
| maker-000052F-augustus-gene-3.39-mRNA-6 | 6 | 000052F | 365507 | 367786 | BUSCO |
| maker-000052F-snap-gene-5.61-mRNA-1 | 6 | 000052F | 578138 | 581600 | BUSCO |
| maker-000052F-augustus-gene-5.44-mRNA-1 | 6 | 000052F | 629732 | 631314 | BUSCO |
| augustus_masked-000058F-processed-gene-3.63-mRNA-1 | 6 | 000058F | 292331 | 294030 | BUSCO |
| maker-000073F-augustus-gene-1.73-mRNA-1 | 6 | 000073F | 118542 | 120390 | BUSCO |
| maker-000076F-augustus-gene-0.76-mRNA-1 | 6 | 000076F | 57333 | 58090 | BUSCO |
| augustus_masked-000076F-processed-gene-1.22-mRNA-1 | 6 | 000076F | 169187 | 175318 | BUSCO |
| maker-000076F-augustus-gene-2.116-mRNA-1 | 6 | 000076F | 193928 | 197353 | BUSCO |
| maker-000076F-augustus-gene-2.103-mRNA-1 | 6 | 000076F | 197350 | 198594 | BUSCO |
| maker-000076F-augustus-gene-2.124-mRNA-7 | 6 | 000076F | 275019 | 277021 | BUSCO |
| genemark-000082F-processed-gene-0.23-mRNA-1 | 6 | 000082F | 42132 | 43658 | BUSCO |
| augustus_masked-000082F-processed-gene-1.53-mRNA-1 | 6 | 000082F | 228912 | 229899 | BUSCO |
| maker-000098F-augustus-gene-0.100-mRNA-1 | 6 | 000098F | 18214 | 19082 | BUSCO |
| maker-000098F-augustus-gene-0.102-mRNA-1 | 6 | 000098F | 29902 | 31477 | BUSCO |
| maker-000098F-augustus-gene-0.116-mRNA-1 | 6 | 000098F | 31042 | 32983 | BUSCO |
| maker-000098F-augustus-gene-0.109-mRNA-1 | 6 | 000098F | 97782 | 100813 | BUSCO |
| maker-000098F-augustus-gene-0.123-mRNA-1 | 6 | 000098F | 116133 | 117900 | BUSCO |
| genemark-000201F-processed-gene-0.21-mRNA-1 | 6 | 000201F | 32616 | 34989 | BUSCO |
| maker-000003F-snap-gene-2.89-mRNA-1 | 7 | 000003F | 285896 | 292701 | BUSCO |
| maker-000003F-augustus-gene-4.101-mRNA-2 | 7 | 000003F | 484613 | 488291 | BUSCO |
| maker-000003F-augustus-gene-9.40-mRNA-4 | 7 | 000003F | 927991 | 934509 | BUSCO |
| maker-000003F-augustus-gene-12.95-mRNA-1 | 7 | 000003F | 1267755 | 1268416 | BUSCO |
| maker-000003F-augustus-gene-14.79-mRNA-1 | 7 | 000003F | 1402988 | 1403550 | BUSCO |
| maker-000003F-augustus-gene-15.96-mRNA-1 | 7 | 000003F | 1578485 | 1581826 | BUSCO |
| maker-000003F-augustus-gene-17.51-mRNA-1 | 7 | 000003F | 1719695 | 1723073 | BUSCO |
| maker-000003F-augustus-gene-17.63-mRNA-2 | 7 | 000003F | 1722502 | 1724399 | BUSCO |
| maker-000003F-augustus-gene-17.69-mRNA-1 | 7 | 000003F | 1770900 | 1771439 | BUSCO |
| maker-000003F-augustus-gene-17.57-mRNA-3 | 7 | 000003F | 1792500 | 1794110 | BUSCO |
| maker-000003F-augustus-gene-18.49-mRNA-1 | 7 | 000003F | 1827520 | 1828497 | BUSCO |
| maker-000003F-augustus-gene-22.53-mRNA-1 | 7 | 000003F | 2257813 | 2264908 | BUSCO |
| maker-000003F-augustus-gene-23.93-mRNA-1 | 7 | 000003F | 2321556 | 2323821 | BUSCO |
| maker-000003F-augustus-gene-26.56-mRNA-2 | 7 | 000003F | 2657769 | 2661159 | BUSCO |
| maker-000003F-augustus-gene-27.96-mRNA-1 | 7 | 000003F | 2753809 | 2756177 | BUSCO |
| augustus_masked-000003F-processed-gene-28.35-mRNA-1 | 7 | 000003F | 2877628 | 2883869 | BUSCO |
| maker-000003F-augustus-gene-30.59-mRNA-1 | 7 | 000003F | 3048369 | 3051240 | BUSCO |
| maker-000003F-augustus-gene-31.38-mRNA-2 | 7 | 000003F | 3098170 | 3100742 | BUSCO |
| maker-000003F-augustus-gene-33.56-mRNA-1 | 7 | 000003F | 3342263 | 3344342 | BUSCO |
| maker-000003F-augustus-gene-34.42-mRNA-5 | 7 | 000003F | 3484813 | 3490335 | BUSCO |
| maker-000003F-augustus-gene-36.71-mRNA-1 | 7 | 000003F | 3603157 | 3606573 | BUSCO |
| maker-000003F-snap-gene-36.22-mRNA-1 | 7 | 000003F | 3655073 | 3655912 | BUSCO |
| maker-000003F-augustus-gene-37.36-mRNA-1 | 7 | 000003F | 3737817 | 3742486 | BUSCO |
| genemark-000003F-processed-gene-38.5-mRNA-1 | 7 | 000003F | 3792066 | 3801433 | BUSCO |
| genemark-000003F-processed-gene-39.32-mRNA-1 | 7 | 000003F | 3954438 | 3967707 | BUSCO |
| maker-000003F-augustus-gene-40.23-mRNA-1 | 7 | 000003F | 4062833 | 4063695 | BUSCO |
| augustus_masked-000003F-processed-gene-42.176-mRNA-1 | 7 | 000003F | 4199153 | 4214946 | BUSCO |
| augustus_masked-000003F-processed-gene-42.191-mRNA-1 | 7 | 000003F | 4345867 | 4349354 | BUSCO |
| maker-000018F-augustus-gene-3.62-mRNA-1 | 7 | 000018F | 318442 | 320291 | BUSCO |
| augustus_masked-000018F-processed-gene-3.97-mRNA-1 | 7 | 000018F | 334790 | 335308 | BUSCO |
| maker-000018F-augustus-gene-3.57-mRNA-1 | 7 | 000018F | 374630 | 375535 | BUSCO |
| maker-000018F-augustus-gene-4.27-mRNA-4 | 7 | 000018F | 415667 | 421134 | BUSCO |
| maker-000018F-augustus-gene-4.29-mRNA-1 | 7 | 000018F | 434770 | 440891 | BUSCO |
| maker-000018F-augustus-gene-5.80-mRNA-1 | 7 | 000018F | 534462 | 535287 | BUSCO |
| maker-000018F-augustus-gene-6.33-mRNA-1 | 7 | 000018F | 627344 | 628745 | BUSCO |
| maker-000018F-augustus-gene-10.60-mRNA-2 | 7 | 000018F | 1005554 | 1007490 | BUSCO |
| maker-000018F-augustus-gene-12.62-mRNA-3 | 7 | 000018F | 1269260 | 1274029 | BUSCO |
| augustus_masked-000018F-processed-gene-13.22-mRNA-1 | 7 | 000018F | 1296095 | 1307525 | BUSCO |
| maker-000018F-augustus-gene-15.44-mRNA-1 | 7 | 000018F | 1508961 | 1511883 | BUSCO |
| augustus_masked-000018F-processed-gene-15.27-mRNA-1 | 7 | 000018F | 1521387 | 1524477 | BUSCO |
| maker-000018F-augustus-gene-16.98-mRNA-1 | 7 | 000018F | 1605567 | 1607013 | BUSCO |
| maker-000018F-augustus-gene-18.1-mRNA-2 | 7 | 000018F | 1822936 | 1826960 | BUSCO |
| augustus_masked-000018F-processed-gene-18.36-mRNA-1 | 7 | 000018F | 1843559 | 1854742 | BUSCO |
| maker-000018F-snap-gene-18.67-mRNA-1 | 7 | 000018F | 1845045 | 1846292 | BUSCO |
| maker-000018F-augustus-gene-24.222-mRNA-1 | 7 | 000018F | 2466510 | 2467506 | BUSCO |
| maker-000018F-augustus-gene-24.204-mRNA-1 | 7 | 000018F | 2489647 | 2501476 | BUSCO |
| maker-000020F-augustus-gene-4.29-mRNA-1 | 7 | 000020F | 430824 | 433045 | BUSCO |
| maker-000020F-augustus-gene-7.87-mRNA-1 | 7 | 000020F | 708616 | 713022 | BUSCO |
| maker-000020F-augustus-gene-13.105-mRNA-1 | 7 | 000020F | 1348595 | 1350773 | BUSCO |
| maker-000020F-snap-gene-15.51-mRNA-1 | 7 | 000020F | 1503486 | 1509159 | BUSCO |
| augustus_masked-000020F-processed-gene-18.28-mRNA-1 | 7 | 000020F | 1890237 | 1894029 | BUSCO |
| maker-000020F-augustus-gene-19.86-mRNA-1 | 7 | 000020F | 1906610 | 1907401 | BUSCO |
| snap_masked-000020F-processed-gene-21.118-mRNA-1 | 7 | 000020F | 2145101 | 2145649 | BUSCO |
| maker-000038F-snap-gene-0.4-mRNA-1 | 7 | 000038F | 86672 | 90486 | BUSCO |
| snap_masked-000038F-processed-gene-1.66-mRNA-1 | 7 | 000038F | 114417 | 116363 | BUSCO |
| maker-000038F-augustus-gene-2.117-mRNA-1 | 7 | 000038F | 193046 | 194565 | BUSCO |
| maker-000038F-augustus-gene-2.120-mRNA-2 | 7 | 000038F | 231922 | 235212 | BUSCO |
| maker-000038F-augustus-gene-2.124-mRNA-1 | 7 | 000038F | 252625 | 257964 | BUSCO |
| maker-000038F-augustus-gene-3.40-mRNA-1 | 7 | 000038F | 287365 | 290836 | BUSCO |
| genemark-000038F-processed-gene-6.1-mRNA-1 | 7 | 000038F | 579430 | 598746 | BUSCO |
| augustus_masked-000038F-processed-gene-6.30-mRNA-1 | 7 | 000038F | 594716 | 601260 | BUSCO |
| augustus_masked-000038F-processed-gene-7.29-mRNA-1 | 7 | 000038F | 702403 | 703085 | BUSCO |
| maker-000038F-augustus-gene-7.91-mRNA-2 | 7 | 000038F | 750003 | 755717 | BUSCO |
| maker-000038F-augustus-gene-7.83-mRNA-1 | 7 | 000038F | 754555 | 766159 | BUSCO |
| maker-000038F-snap-gene-7.77-mRNA-5 | 7 | 000038F | 770341 | 773651 | BUSCO |
| maker-000038F-augustus-gene-8.151-mRNA-1 | 7 | 000038F | 882136 | 882748 | BUSCO |
| maker-000038F-augustus-gene-8.135-mRNA-1 | 7 | 000038F | 936009 | 937741 | BUSCO |
| augustus_masked-000054F-processed-gene-2.73-mRNA-1 | 7 | 000054F | 182773 | 197392 | BUSCO |
| maker-000005F-snap-gene-9.47-mRNA-1 | 8 | 000005F | 962832 | 964778 | BUSCO |
| maker-000005F-augustus-gene-13.73-mRNA-1 | 8 | 000005F | 1298330 | 1304119 | BUSCO |
| maker-000005F-snap-gene-16.30-mRNA-1 | 8 | 000005F | 1652623 | 1654519 | BUSCO |
| maker-000005F-augustus-gene-17.31-mRNA-4 | 8 | 000005F | 1737441 | 1741504 | BUSCO |
| genemark-000005F-processed-gene-18.31-mRNA-1 | 8 | 000005F | 1839280 | 1852521 | BUSCO |
| maker-000005F-snap-gene-22.39-mRNA-1 | 8 | 000005F | 2197538 | 2198659 | BUSCO |
| maker-000005F-augustus-gene-23.84-mRNA-1 | 8 | 000005F | 2296925 | 2299059 | BUSCO |
| maker-000005F-augustus-gene-23.86-mRNA-1 | 8 | 000005F | 2346790 | 2348803 | BUSCO |
| maker-000005F-augustus-gene-23.82-mRNA-4 | 8 | 000005F | 2349719 | 2354854 | BUSCO |
| maker-000005F-augustus-gene-25.65-mRNA-1 | 8 | 000005F | 2504552 | 2507402 | BUSCO |
| maker-000005F-augustus-gene-28.52-mRNA-2 | 8 | 000005F | 2811645 | 2820974 | BUSCO |
| maker-000005F-augustus-gene-29.123-mRNA-1 | 8 | 000005F | 2964088 | 2967240 | BUSCO |
| maker-000005F-augustus-gene-31.22-mRNA-1 | 8 | 000005F | 3113270 | 3115855 | BUSCO |
| maker-000005F-augustus-gene-33.83-mRNA-1 | 8 | 000005F | 3324086 | 3328080 | BUSCO |
| genemark-000005F-processed-gene-33.68-mRNA-1 | 8 | 000005F | 3328901 | 3332028 | BUSCO |
| genemark-000005F-processed-gene-34.25-mRNA-1 | 8 | 000005F | 3394727 | 3395883 | BUSCO |
| maker-000005F-augustus-gene-34.85-mRNA-1 | 8 | 000005F | 3453242 | 3476464 | BUSCO |
| maker-000005F-augustus-gene-35.48-mRNA-3 | 8 | 000005F | 3561983 | 3567506 | BUSCO |
| maker-000005F-augustus-gene-37.53-mRNA-1 | 8 | 000005F | 3729126 | 3730423 | BUSCO |
| maker-000015F-augustus-gene-2.41-mRNA-4 | 8 | 000015F | 252538 | 255611 | BUSCO |
| maker-000015F-augustus-gene-3.80-mRNA-1 | 8 | 000015F | 367655 | 369825 | BUSCO |
| maker-000015F-snap-gene-3.20-mRNA-1 | 8 | 000015F | 370021 | 372380 | BUSCO |
| genemark-000015F-processed-gene-5.17-mRNA-1 | 8 | 000015F | 562900 | 572548 | BUSCO |
| maker-000015F-augustus-gene-9.47-mRNA-1 | 8 | 000015F | 985831 | 987548 | BUSCO |
| maker-000015F-augustus-gene-10.126-mRNA-1 | 8 | 000015F | 1036502 | 1038387 | BUSCO |
| maker-000015F-augustus-gene-11.16-mRNA-1 | 8 | 000015F | 1091801 | 1095650 | BUSCO |
| augustus_masked-000015F-processed-gene-11.0-mRNA-1 | 8 | 000015F | 1095466 | 1100703 | BUSCO |
| maker-000015F-augustus-gene-11.21-mRNA-1 | 8 | 000015F | 1149791 | 1153494 | BUSCO |
| maker-000015F-augustus-gene-12.39-mRNA-1 | 8 | 000015F | 1254642 | 1257152 | BUSCO |
| genemark-000015F-processed-gene-13.24-mRNA-1 | 8 | 000015F | 1351696 | 1357055 | BUSCO |
| maker-000015F-augustus-gene-15.51-mRNA-1 | 8 | 000015F | 1535284 | 1537596 | BUSCO |
| augustus_masked-000015F-processed-gene-16.1-mRNA-1 | 8 | 000015F | 1647810 | 1652574 | BUSCO |
| maker-000015F-augustus-gene-16.58-mRNA-1 | 8 | 000015F | 1652689 | 1656611 | BUSCO |
| maker-000015F-augustus-gene-16.52-mRNA-3 | 8 | 000015F | 1670645 | 1677791 | BUSCO |
| maker-000015F-augustus-gene-17.11-mRNA-1 | 8 | 000015F | 1733808 | 1735464 | BUSCO |
| maker-000015F-augustus-gene-17.2-mRNA-1 | 8 | 000015F | 1735300 | 1736956 | BUSCO |
| maker-000015F-snap-gene-17.60-mRNA-5 | 8 | 000015F | 1764270 | 1771393 | BUSCO |
| maker-000015F-augustus-gene-18.5-mRNA-1 | 8 | 000015F | 1871046 | 1872457 | BUSCO |
| maker-000015F-snap-gene-19.56-mRNA-3 | 8 | 000015F | 1875026 | 1877409 | BUSCO |
| maker-000015F-augustus-gene-20.22-mRNA-1 | 8 | 000015F | 2055015 | 2062074 | BUSCO |
| maker-000015F-augustus-gene-20.23-mRNA-1 | 8 | 000015F | 2062315 | 2063915 | BUSCO |
| snap_masked-000015F-processed-gene-21.44-mRNA-1 | 8 | 000015F | 2147428 | 2148668 | BUSCO |
| genemark-000015F-processed-gene-22.46-mRNA-1 | 8 | 000015F | 2271668 | 2273063 | BUSCO |
| maker-000015F-augustus-gene-22.106-mRNA-4 | 8 | 000015F | 2274754 | 2277062 | BUSCO |
| maker-000015F-augustus-gene-24.65-mRNA-1 | 8 | 000015F | 2476622 | 2479263 | BUSCO |
| maker-000015F-augustus-gene-25.77-mRNA-1 | 8 | 000015F | 2502451 | 2504250 | BUSCO |
| maker-000015F-snap-gene-25.55-mRNA-3 | 8 | 000015F | 2586613 | 2593069 | BUSCO |
| maker-000015F-augustus-gene-25.70-mRNA-1 | 8 | 000015F | 2589010 | 2589756 | BUSCO |
| maker-000030F-augustus-gene-3.87-mRNA-1 | 8 | 000030F | 352899 | 356142 | BUSCO |
| maker-000030F-augustus-gene-13.82-mRNA-1 | 8 | 000030F | 1317323 | 1318587 | BUSCO |
| maker-000031F-augustus-gene-1.74-mRNA-1 | 8 | 000031F | 124420 | 128425 | BUSCO |
| maker-000031F-augustus-gene-3.21-mRNA-1 | 8 | 000031F | 354044 | 355100 | BUSCO |
| maker-000031F-augustus-gene-6.72-mRNA-2 | 8 | 000031F | 652586 | 655011 | BUSCO |
| maker-000031F-augustus-gene-8.92-mRNA-1 | 8 | 000031F | 803727 | 806185 | BUSCO |
| maker-000031F-augustus-gene-8.109-mRNA-1 | 8 | 000031F | 874614 | 875823 | BUSCO |
| maker-000031F-augustus-gene-9.84-mRNA-1 | 8 | 000031F | 927736 | 931454 | BUSCO |
| augustus_masked-000050F-processed-gene-0.44-mRNA-1 | 8 | 000050F | 29321 | 41562 | BUSCO |
| maker-000050F-augustus-gene-1.47-mRNA-3 | 8 | 000050F | 83613 | 88061 | BUSCO |
| snap_masked-000050F-processed-gene-1.69-mRNA-1 | 8 | 000050F | 116605 | 117009 | BUSCO |
| genemark-000050F-processed-gene-3.60-mRNA-1 | 8 | 000050F | 311675 | 318549 | BUSCO |
| augustus_masked-000050F-processed-gene-5.41-mRNA-1 | 8 | 000050F | 467130 | 469946 | BUSCO |
| maker-000050F-augustus-gene-5.155-mRNA-1 | 8 | 000050F | 611386 | 611886 | BUSCO |
| maker-000050F-augustus-gene-5.156-mRNA-2 | 8 | 000050F | 616000 | 618602 | BUSCO |
| maker-000050F-augustus-gene-5.137-mRNA-1 | 8 | 000050F | 654255 | 663748 | BUSCO |
| genemark-000063F-processed-gene-2.66-mRNA-1 | 8 | 000063F | 252068 | 255681 | BUSCO |
| maker-000063F-augustus-gene-4.25-mRNA-1 | 8 | 000063F | 437307 | 439856 | BUSCO |
| maker-000014F-augustus-gene-0.18-mRNA-1 | 9 | 000014F | 22139 | 23733 | BUSCO |
| genemark-000014F-processed-gene-0.51-mRNA-1 | 9 | 000014F | 32030 | 37194 | BUSCO |
| maker-000014F-augustus-gene-1.37-mRNA-1 | 9 | 000014F | 131239 | 132072 | BUSCO |
| genemark-000014F-processed-gene-1.19-mRNA-1 | 9 | 000014F | 134236 | 145813 | BUSCO |
| augustus_masked-000014F-processed-gene-2.43-mRNA-1 | 9 | 000014F | 254379 | 260827 | BUSCO |
| genemark-000014F-processed-gene-2.36-mRNA-1 | 9 | 000014F | 259271 | 263803 | BUSCO |
| augustus_masked-000014F-processed-gene-2.50-mRNA-1 | 9 | 000014F | 271890 | 272445 | BUSCO |
| maker-000014F-augustus-gene-4.46-mRNA-1 | 9 | 000014F | 404869 | 405764 | BUSCO |
| maker-000014F-augustus-gene-7.32-mRNA-1 | 9 | 000014F | 725397 | 727758 | BUSCO |
| maker-000014F-snap-gene-8.69-mRNA-2 | 9 | 000014F | 874018 | 876297 | BUSCO |
| genemark-000014F-processed-gene-9.26-mRNA-1 | 9 | 000014F | 900496 | 906671 | BUSCO |
| genemark-000014F-processed-gene-9.23-mRNA-1 | 9 | 000014F | 933806 | 938075 | BUSCO |
| maker-000014F-augustus-gene-9.49-mRNA-1 | 9 | 000014F | 934257 | 935373 | BUSCO |
| maker-000014F-augustus-gene-13.48-mRNA-1 | 9 | 000014F | 1306086 | 1308109 | BUSCO |
| maker-000014F-augustus-gene-13.58-mRNA-3 | 9 | 000014F | 1356101 | 1358784 | BUSCO |
| maker-000014F-augustus-gene-17.85-mRNA-2 | 9 | 000014F | 1762212 | 1763689 | BUSCO |
| maker-000014F-snap-gene-18.10-mRNA-3 | 9 | 000014F | 1806978 | 1810474 | BUSCO |
| maker-000014F-snap-gene-20.46-mRNA-2 | 9 | 000014F | 1997164 | 1998968 | BUSCO |
| maker-000014F-augustus-gene-20.53-mRNA-1 | 9 | 000014F | 1999243 | 2000252 | BUSCO |
| maker-000014F-augustus-gene-20.63-mRNA-1 | 9 | 000014F | 2011442 | 2020142 | BUSCO |
| maker-000014F-snap-gene-20.50-mRNA-1 | 9 | 000014F | 2020510 | 2024283 | BUSCO |
| maker-000014F-augustus-gene-20.59-mRNA-1 | 9 | 000014F | 2076803 | 2078861 | BUSCO |
| maker-000014F-augustus-gene-21.28-mRNA-1 | 9 | 000014F | 2115729 | 2116584 | BUSCO |
| maker-000014F-augustus-gene-22.13-mRNA-1 | 9 | 000014F | 2244433 | 2247237 | BUSCO |
| maker-000014F-augustus-gene-23.14-mRNA-1 | 9 | 000014F | 2372360 | 2376562 | BUSCO |
| genemark-000014F-processed-gene-24.56-mRNA-1 | 9 | 000014F | 2456752 | 2457427 | BUSCO |
| maker-000014F-augustus-gene-25.40-mRNA-3 | 9 | 000014F | 2498533 | 2502717 | BUSCO |
| maker-000014F-augustus-gene-25.55-mRNA-2 | 9 | 000014F | 2505047 | 2508037 | BUSCO |
| maker-000014F-augustus-gene-25.58-mRNA-1 | 9 | 000014F | 2536888 | 2538893 | BUSCO |
| maker-000029F-augustus-gene-2.101-mRNA-1 | 9 | 000029F | 283964 | 284972 | BUSCO |
| maker-000029F-augustus-gene-5.72-mRNA-1 | 9 | 000029F | 488251 | 494317 | BUSCO |
| maker-000029F-augustus-gene-12.71-mRNA-1 | 9 | 000029F | 1224099 | 1224978 | BUSCO |
| maker-000029F-snap-gene-13.73-mRNA-1 | 9 | 000029F | 1436515 | 1437647 | BUSCO |
| maker-000029F-augustus-gene-13.131-mRNA-1 | 9 | 000029F | 1495251 | 1497044 | BUSCO |
| genemark-000042F-processed-gene-1.0-mRNA-1 | 9 | 000042F | 88036 | 94574 | BUSCO |
| augustus_masked-000042F-processed-gene-1.23-mRNA-1 | 9 | 000042F | 97003 | 105152 | BUSCO |
| maker-000042F-augustus-gene-1.63-mRNA-1 | 9 | 000042F | 118593 | 121798 | BUSCO |
| maker-000042F-augustus-gene-4.53-mRNA-1 | 9 | 000042F | 407113 | 410458 | BUSCO |
| maker-000042F-augustus-gene-5.104-mRNA-1 | 9 | 000042F | 492716 | 493975 | BUSCO |
| snap_masked-000042F-processed-gene-5.143-mRNA-1 | 9 | 000042F | 582035 | 583896 | BUSCO |
| augustus_masked-000059F-processed-gene-2.46-mRNA-1 | 9 | 000059F | 191440 | 200772 | BUSCO |
| maker-000059F-augustus-gene-2.59-mRNA-1 | 9 | 000059F | 278492 | 280833 | BUSCO |
| maker-000059F-augustus-gene-3.19-mRNA-3 | 9 | 000059F | 368750 | 371261 | BUSCO |
| augustus_masked-000066F-processed-gene-0.63-mRNA-1 | 9 | 000066F | 10273 | 19137 | BUSCO |
| maker-000066F-augustus-gene-0.119-mRNA-1 | 9 | 000066F | 33941 | 36430 | BUSCO |
| maker-000066F-augustus-gene-0.124-mRNA-1 | 9 | 000066F | 60064 | 63493 | BUSCO |
| augustus_masked-000066F-processed-gene-3.35-mRNA-1 | 9 | 000066F | 302953 | 308100 | BUSCO |
| maker-000079F-augustus-gene-0.76-mRNA-1 | 9 | 000079F | 8979 | 9766 | BUSCO |
| genemark-000102F-processed-gene-0.120-mRNA-1 | 9 | 000102F | 72665 | 73760 | BUSCO |
| augustus_masked-000107F-processed-gene-0.85-mRNA-1 | 9 | 000107F | 63297 | 72399 | BUSCO |
| augustus_masked-000169F-processed-gene-0.35-mRNA-1 | 9 | 000169F | 36453 | 38874 | BUSCO |
| maker-000169F-augustus-gene-0.60-mRNA-1 | 9 | 000169F | 39633 | 40613 | BUSCO |
| maker-000006F-augustus-gene-8.97-mRNA-1 | 10 | 000006F | 809163 | 812606 | BUSCO |
| maker-000006F-augustus-gene-9.81-mRNA-1 | 10 | 000006F | 946445 | 947525 | BUSCO |
| maker-000006F-snap-gene-11.37-mRNA-1 | 10 | 000006F | 1105751 | 1113243 | BUSCO |
| maker-000006F-augustus-gene-18.45-mRNA-1 | 10 | 000006F | 1860276 | 1861591 | BUSCO |
| genemark-000006F-processed-gene-21.51-mRNA-1 | 10 | 000006F | 2084010 | 2085762 | BUSCO |
| genemark-000006F-processed-gene-22.40-mRNA-1 | 10 | 000006F | 2186663 | 2192959 | BUSCO |
| maker-000006F-augustus-gene-26.50-mRNA-1 | 10 | 000006F | 2589616 | 2597571 | BUSCO |
| genemark-000006F-processed-gene-26.43-mRNA-1 | 10 | 000006F | 2604639 | 2610172 | BUSCO |
| maker-000006F-augustus-gene-28.21-mRNA-1 | 10 | 000006F | 2816228 | 2820218 | BUSCO |
| maker-000006F-snap-gene-29.44-mRNA-2 | 10 | 000006F | 2934818 | 2937002 | BUSCO |
| maker-000006F-augustus-gene-29.17-mRNA-1 | 10 | 000006F | 2937011 | 2941156 | BUSCO |
| maker-000006F-augustus-gene-29.18-mRNA-1 | 10 | 000006F | 2948639 | 2951416 | BUSCO |
| maker-000006F-augustus-gene-29.20-mRNA-1 | 10 | 000006F | 2957187 | 2959052 | BUSCO |
| maker-000013F-augustus-gene-3.27-mRNA-2 | 10 | 000013F | 299895 | 306144 | BUSCO |
| maker-000013F-snap-gene-3.68-mRNA-4 | 10 | 000013F | 322131 | 325693 | BUSCO |
| maker-000013F-snap-gene-6.44-mRNA-2 | 10 | 000013F | 643648 | 646193 | BUSCO |
| maker-000013F-augustus-gene-10.67-mRNA-1 | 10 | 000013F | 996146 | 997317 | BUSCO |
| genemark-000013F-processed-gene-12.55-mRNA-1 | 10 | 000013F | 1286798 | 1290856 | BUSCO |
| maker-000013F-augustus-gene-12.26-mRNA-1 | 10 | 000013F | 1290844 | 1292695 | BUSCO |
| maker-000013F-augustus-gene-14.81-mRNA-1 | 10 | 000013F | 1463350 | 1467622 | BUSCO |
| maker-000013F-augustus-gene-15.45-mRNA-1 | 10 | 000013F | 1552044 | 1559777 | BUSCO |
| maker-000013F-augustus-gene-17.66-mRNA-1 | 10 | 000013F | 1763953 | 1765443 | BUSCO |
| snap_masked-000013F-processed-gene-17.89-mRNA-1 | 10 | 000013F | 1767180 | 1769506 | BUSCO |
| maker-000013F-snap-gene-18.69-mRNA-4 | 10 | 000013F | 1802324 | 1802842 | BUSCO |
| augustus_masked-000013F-processed-gene-20.32-mRNA-1 | 10 | 000013F | 2004932 | 2009491 | BUSCO |
| maker-000013F-snap-gene-24.51-mRNA-1 | 10 | 000013F | 2426765 | 2427855 | BUSCO |
| maker-000013F-augustus-gene-24.45-mRNA-1 | 10 | 000013F | 2481640 | 2483988 | BUSCO |
| genemark-000013F-processed-gene-25.28-mRNA-1 | 10 | 000013F | 2535543 | 2537667 | BUSCO |
| maker-000013F-augustus-gene-26.6-mRNA-1 | 10 | 000013F | 2629320 | 2632836 | BUSCO |
| augustus_masked-000013F-processed-gene-27.71-mRNA-1 | 10 | 000013F | 2673182 | 2693929 | BUSCO |
| maker-000013F-augustus-gene-27.90-mRNA-1 | 10 | 000013F | 2699030 | 2702075 | BUSCO |
| genemark-000034F-processed-gene-2.42-mRNA-1 | 10 | 000034F | 198530 | 201124 | BUSCO |
| maker-000034F-augustus-gene-2.33-mRNA-1 | 10 | 000034F | 200168 | 204444 | BUSCO |
| maker-000034F-augustus-gene-2.30-mRNA-1 | 10 | 000034F | 252243 | 254513 | BUSCO |
| maker-000034F-augustus-gene-5.20-mRNA-2 | 10 | 000034F | 510740 | 511923 | BUSCO |
| maker-000034F-augustus-gene-11.111-mRNA-1 | 10 | 000034F | 1126456 | 1128830 | BUSCO |
| maker-000034F-augustus-gene-11.113-mRNA-2 | 10 | 000034F | 1149962 | 1151176 | BUSCO |
| maker-000035F-augustus-gene-0.70-mRNA-2 | 10 | 000035F | 13983 | 18568 | BUSCO |
| maker-000035F-augustus-gene-0.69-mRNA-3 | 10 | 000035F | 78780 | 81065 | BUSCO |
| maker-000035F-augustus-gene-1.76-mRNA-5 | 10 | 000035F | 92670 | 96380 | BUSCO |
| maker-000035F-augustus-gene-1.88-mRNA-2 | 10 | 000035F | 104685 | 105714 | BUSCO |
| maker-000035F-augustus-gene-1.96-mRNA-1 | 10 | 000035F | 174072 | 175815 | BUSCO |
| genemark-000035F-processed-gene-3.37-mRNA-1 | 10 | 000035F | 339840 | 348406 | BUSCO |
| maker-000035F-augustus-gene-4.107-mRNA-1 | 10 | 000035F | 434084 | 435027 | BUSCO |
| snap_masked-000035F-processed-gene-6.74-mRNA-1 | 10 | 000035F | 622682 | 637802 | BUSCO |
| snap_masked-000035F-processed-gene-6.70-mRNA-1 | 10 | 000035F | 680925 | 688489 | BUSCO |
| maker-000035F-snap-gene-7.78-mRNA-1 | 10 | 000035F | 708654 | 709954 | BUSCO |
| maker-000035F-augustus-gene-7.60-mRNA-2 | 10 | 000035F | 757150 | 759874 | BUSCO |
| augustus_masked-000035F-processed-gene-8.14-mRNA-1 | 10 | 000035F | 860990 | 863124 | BUSCO |
| genemark-000035F-processed-gene-9.49-mRNA-1 | 10 | 000035F | 907850 | 911468 | BUSCO |
| maker-000035F-augustus-gene-9.118-mRNA-1 | 10 | 000035F | 985266 | 987124 | BUSCO |
| maker-000035F-augustus-gene-9.119-mRNA-1 | 10 | 000035F | 999009 | 1000141 | BUSCO |
| maker-000078F-augustus-gene-0.25-mRNA-1 | 10 | 000078F | 20048 | 23070 | BUSCO |
| maker-000078F-augustus-gene-1.56-mRNA-1 | 10 | 000078F | 138162 | 139217 | BUSCO |
| maker-000136F-augustus-gene-0.8-mRNA-1 | 10 | 000136F | 1962 | 2752 | BUSCO |
| maker-000136F-augustus-gene-0.10-mRNA-1 | 10 | 000136F | 22386 | 23178 | BUSCO |
| maker-000041F-augustus-gene-0.17-mRNA-5 | n.a. | 000041F | 3592 | 12618 | BUSCO |
| maker-000105F-augustus-gene-0.63-mRNA-1 | n.a. | 000105F | 45367 | 54181 | BUSCO |
| maker-000168F-augustus-gene-0.75-mRNA-1 | n.a. | 000168F | 4397 | 4854 | BUSCO |

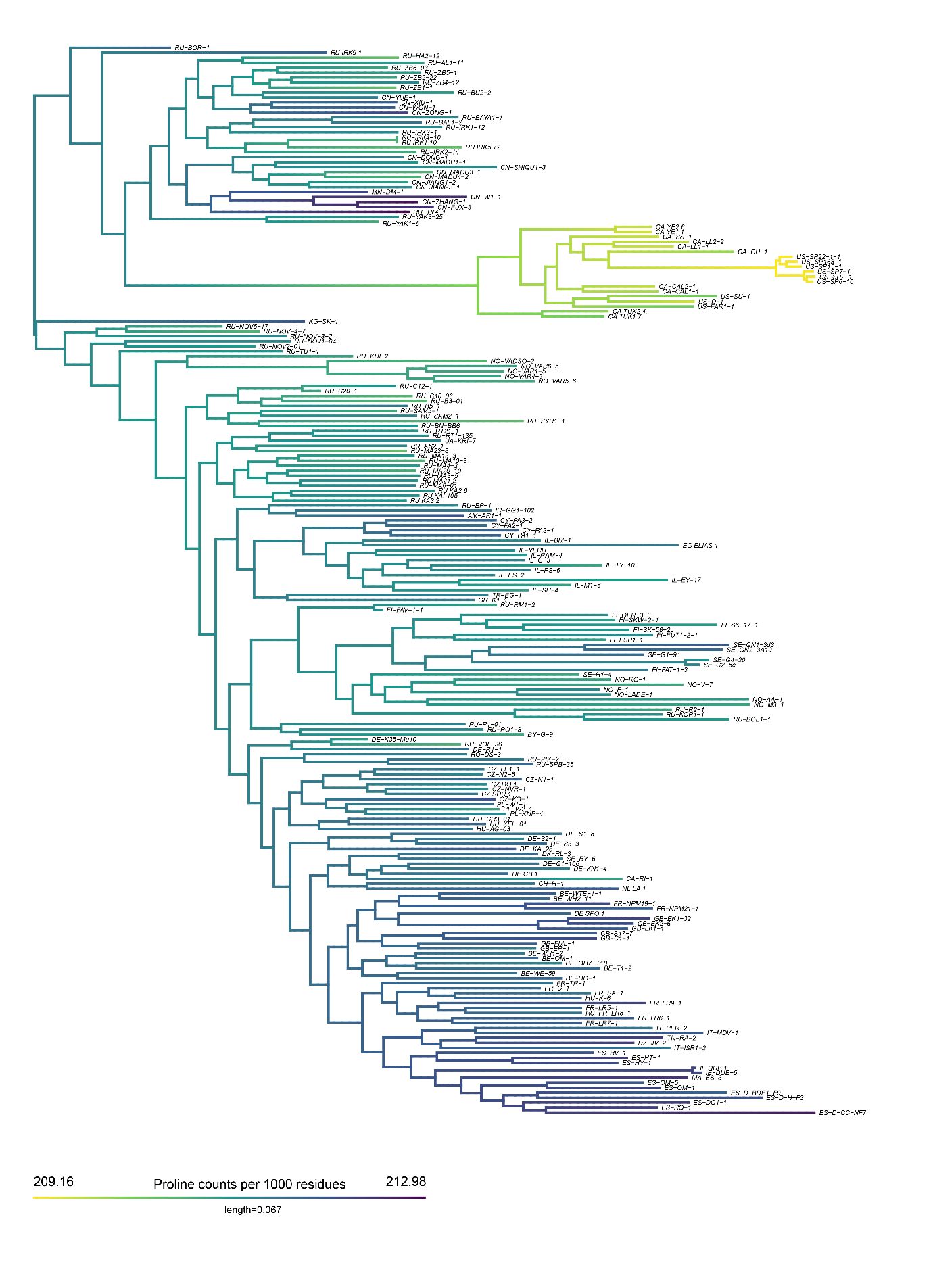

### **Figure S1.** Phylogenetic comparative tree for proline content variation in collagen across the 215 D. magna genotypes used in this study.

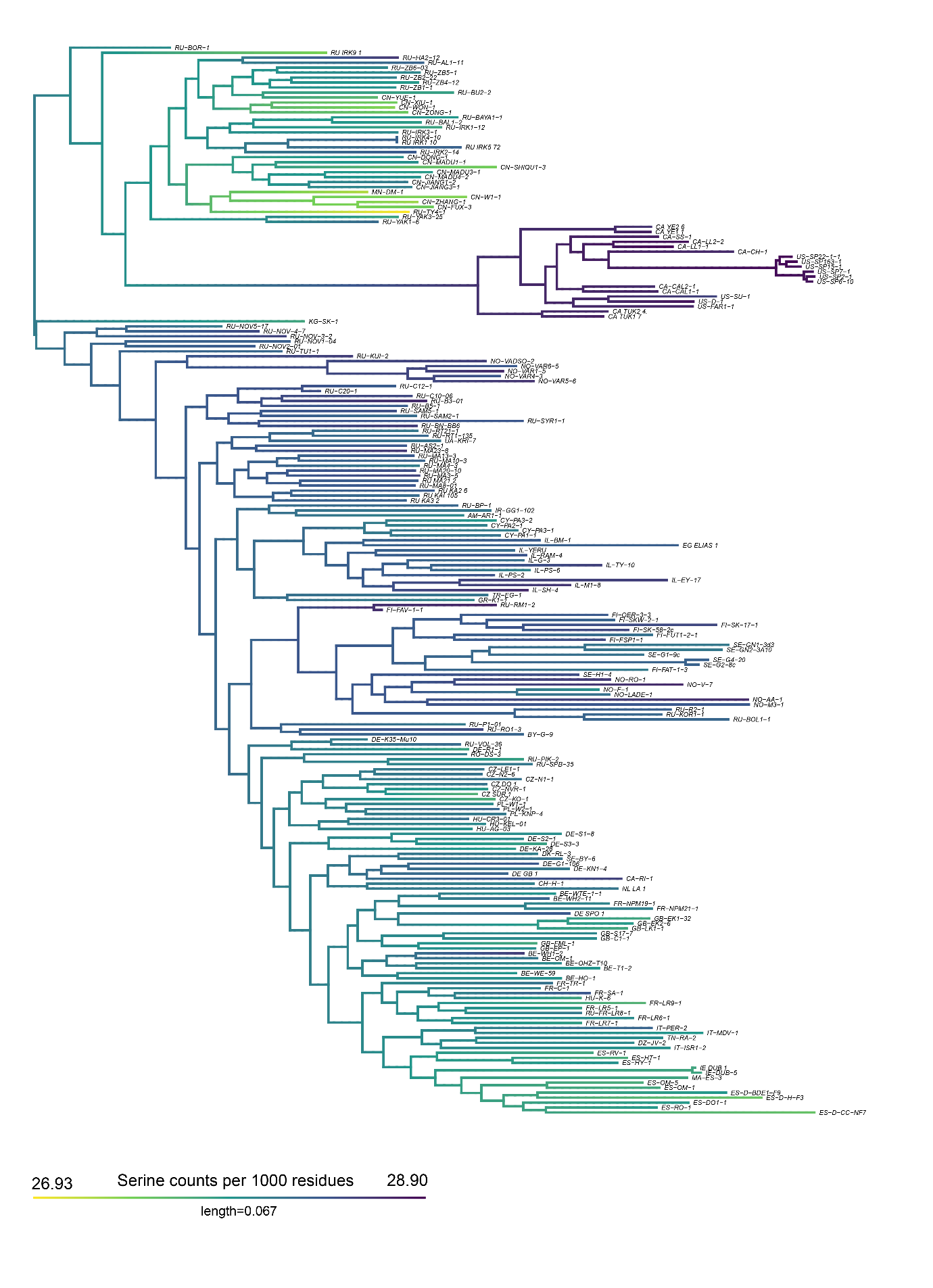

### **Figure S2**. Phylogenetic comparative tree for serine content variation in collagen across the 215 D. magna genotypes used in this study.
